# Peri-centrosomal localization of small interfering RNAs in *C. elegans*

**DOI:** 10.1101/2024.07.11.603161

**Authors:** Qile Jin, Xuezhu Feng, Minjie Hong, Ke Wang, Xiangyang Chen, Jiewei Cheng, Yan Kuang, Xiaoyue Si, Mingjing Xu, Xinya Huang, Shouhong Guang, Chengming Zhu

## Abstract

The centrosome is the microtubule-organizing center and a crucial part of cell division. Centrosomal RNAs (cnRNAs) have been reported to enable precise spatiotemporal control of gene expression during cell division in many species. Whether and how cnRNAs exist in *C. elegans* are unclear. Here, using the nuclear RNAi Argonaute protein NRDE-3 as a reporter, we observed potential peri-centrosome localized small interfering (si)RNAs in *C. elegans*. NRDE-3 was previously shown to associate with pre-mRNAs and pre-rRNAs via a process involving the presence of complementary siRNAs. We generated a GFP-NRDE-3 knock-in transgene through CRISPR/Cas9 technology and observed that NRDE-3 formed peri-centrosomal foci neighboring the tubulin protein TBB-2, other centriole proteins and pericentriolar material (PCM) components in *C. elegans* embryos. The peri-centrosomal accumulation of NRDE-3 depends on RNA-dependent RNA polymerase (RdRP)-synthesized 22G siRNAs and the PAZ domain of NRDE-3, which is essential for siRNA binding. Mutation of *eri-1, ergo-1*, or *drh-3* significantly increased the percentage of pericentrosome-enriched NRDE-3. At the metaphase of the cell cycle, NRDE-3 was enriched in both the peri-centrosomal region and the spindle. Moreover, the integrity of centriole proteins and pericentriolar material (PCM) components is also required for the peri-centrosomal accumulation of NRDE-3. Therefore, we concluded that siRNAs could accumulate in the peri-centrosomal region in *C. elegans* and suggested that the peri-centrosomal region may also be a platform for RNAi-mediated gene regulation.

## Introduction

Centrosomes are nonmembrane organelles. The core of the centrosome consists of a pair of orthogonally oriented centrioles surrounded by a dynamic assembly of proteins known as the pericentriolar material (PCM). Centrosomes are responsible for microtubule nucleation and organization, spindle assembly, cell division, and cell polarity (1).

Centrosomal RNAs (cnRNAs) were identified by studying the oocytes of the surf clam *(Spisula solidissima)* as early as 2006 (2). Scientists have used purification methods and labeling approaches to discover cnRNAs. With the development of new technologies, localization-based approaches such as single molecular FISH (smFISH) and transcriptomic approaches such as single-cell sequencing have accelerated the characterization of cnRNAs (3). Centrosomal RNAs have since been identified in diverse model systems, including *Ilyanassa, Spisula, Drosophila, Xenopus,* zebrafish, mollusk, and mammalian cell lines (2, 4–10), suggesting that the localization of mRNAs to the centrosome is an evolutionarily conserved phenomenon. However, it is unclear whether and how cnRNAs exist in *C. elegans*.

Centrosomal RNAs are proposed to have three main functions. First, RNAs localize to the centrosome to regulate centrosome function and mitotic integrity via co-translational mechanisms. Since cell division is a highly dynamic process, cnRNAs and co-translational centrosome proteins may effectively respond to cell cycle demands (11). Mislocalized cnRNAs have been shown to disrupt microtubule organization and induce mitotic errors (12, 13). Second, asymmetrically localized cnRNAs may contribute to asymmetric cell division and lead to embryonic patterning and selective inheritance of specific transcripts (5, 8, 10, 14). Third, cnRNAs may support the structural integrity of centrosomes and promote phase separation within the centrosome (15).

RNA interference (RNAi) is a conserved mechanism that silences complementary mRNAs via Argonaute/siRNA complexes at the posttranscriptional level (16). siRNAs are generated by the conserved ribonuclease Dicer and bound by Argonaute (Ago) proteins to degrade targeted RNA or inhibit its translation (17–19). In addition, in many organisms, RNA-dependent RNA polymerases (RdRPs) can synthesize secondary siRNAs to amplify silencing signals (20, 21). In *C. elegans*, siRNAs silence nucleus-localized RNAs via the nuclear RNAi defective (Nrde) pathway (22, 23). The somatic nuclear Argonaute NRDE-3, or HRDE-1 in the germline, transports 22G siRNAs from the cytoplasm to the nucleus, where they bind to nascent transcripts and recruit other NRDE factors to inhibit RNA polymerase I/II elongation and induce H3K9 and H3K27 trimethylation (23–26). NRDE-3 has been used to reveal tissue-specific gene expression (22), antisense ribosomal siRNAs (risiRNAs) (27, 28), and transcriptional dynamics *in vivo* (29, 30)

Small RNAs have been shown to localize to many special subcellular organelles, including membrane compartments (mitochondria, secreted exosomes and the endoplasmic reticulum) and membraneless compartments (stress granules, processing bodies and germ granules) (27, 28, 31–33). Small RNAs play vital roles in the maintenance of organelle homeostasis. Dysfunction of small RNAs leads to various diseases, such as neurodegenerative diseases and cancer (34, 35).

Here, using CRISPR/Cas9 technology, we generated a GFP-tagged NRDE-3 knock-in transgene. NRDE-3 was expressed in oocytes, early and late embryos, and somatic cells and was enriched in the nucleus. Strikingly, we observed that NRDE-3 could also accumulate in the peri-centrosomal foci in a manner dependent on its PAZ domain and RdRPs. Therefore, our findings suggest that the peri-centrosomal region may also be a platform for RNAi-mediated gene regulation.

## Results

### NRDE-3 is widely expressed in germlines, oocytes, early and late embryos and soma

Previously, we identified the nuclear Argonaute protein NRDE-3 by forward genetic screening to search for factors required for nuclear RNAi (22). We constructed a low-copy green fluorescent protein (GFP) and full-length NRDE-3 fusion transgene driven by the *nrde-3* promoter through a microparticle-mediated bombardment method (Fig. 1A), which rescued the nuclear RNAi defects of *nrde-3*(*-*) animals (22). GFP::NRDE-3(ggIS1, bombardment) was expressed in somatic cells after the ∼80-cell stage of embryogenesis in embryos and larvae and was predominantly localized to the nucleus in a small RNA-dependent manner. The GFP::NRDE-3(ggIS1, bombardment) transgenic animals contained approximately 11 copies of the GFP::NRDE-3 fusion gene in the genome (Fig. S1A).

**Figure 1.**
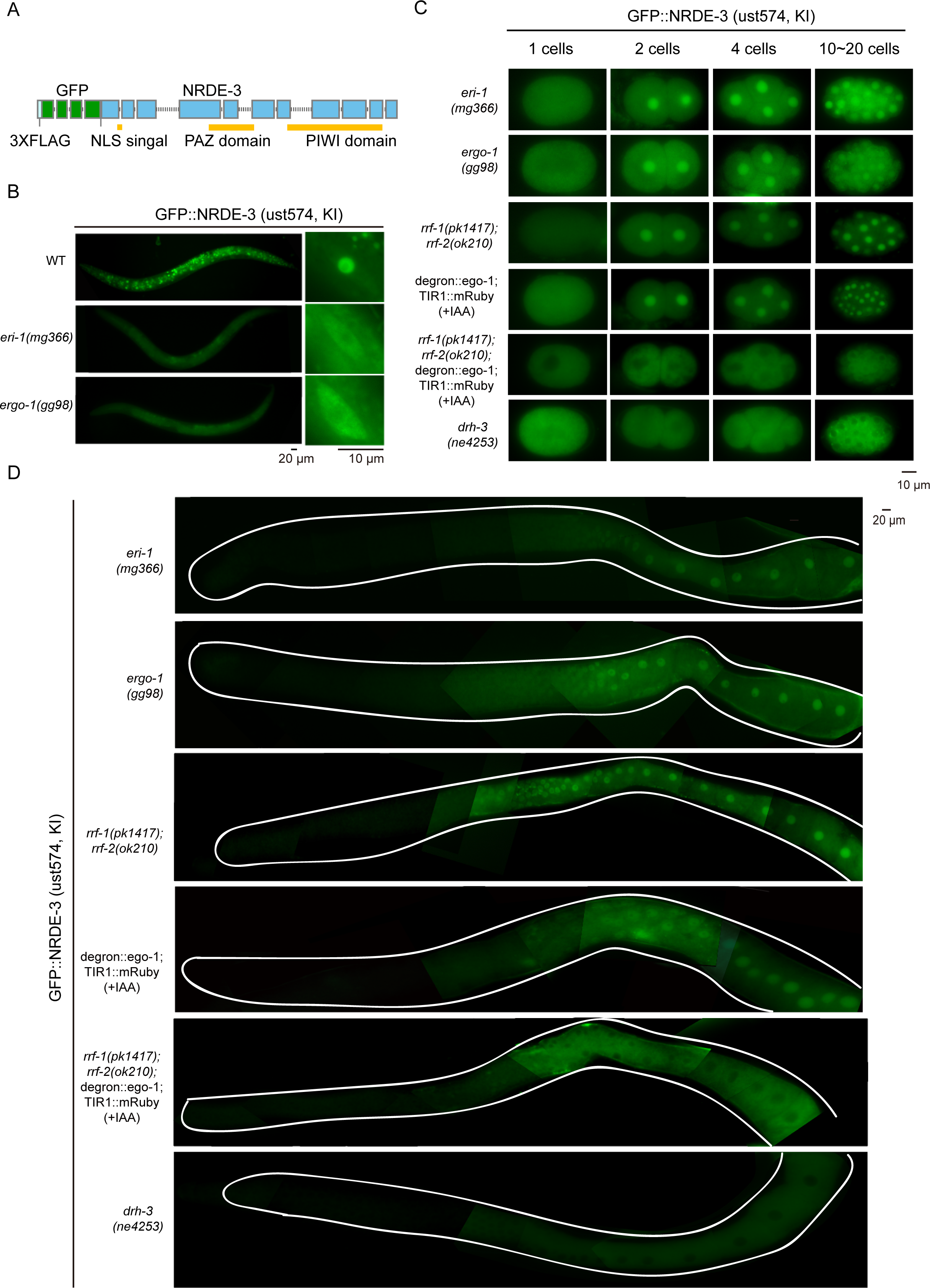
NRDE-3 is widely expressed in germlines, oocytes, early and late embryos and somatic cells. (A) Schematic of NRDE-3 and the 3xFLAG::GFP tag. NLS, nuclear localization signal. (B) Fluorescence microscopy images of GFP::NRDE-3(ust574, KI) in L4 stage animals (left) and seam cells (right) of the indicated genotypes. (C) Fluorescence microscopy images of GFP::NRDE-3 (ust574, KI) in embryonic stages in the indicated background. For *degron::ego-1* animals, synchronized embryos were grown on NGM plates seeded with OP50 supplemented with 1 mM IAA. For the *rrf-1;rrf-2;degron::ego-1* animals, synchronized animals were grown on OP50 plates and shifted to NGM plates containing 1 mM IAA at the L4 stage. Pictures were taken 48 hours after IAA treatment. (D) Fluorescence microscopy images of the dissected gravid adult germline.

Interestingly, in addition to rescuing the nuclear RNAi defects of *nrde-3*(*-*) animals, the GFP::NRDE-3(ggIS1, bombardment) strain also exhibited an enhanced RNAi (Eri) phenotype (36). *lir-1* and *lin-26* are cotranscribed in an operon, and *lir-1* mutant is viable, whereas loss of *lin-26* results in larval lethality. RNAi targeting *lir-1* also caused larval lethality via nuclear RNAi machinery-mediated gene silencing of nucleus-localized *lir-1–lin-26* polycistronic RNA (22, 37, 38). For instance, growing L1 wild-type N2 animals on bacteria expressing dsRNAs targeting *the lir-1* gene caused early larval arrest in F1 progeny (Fig. S1B). *nrde-3*(*-*) animals are completely resistant to *lir-1* RNAi. Strikingly, introducing the GFP::NRDE-3(ggIS1, bombardment) transgene not only rescued *lir-1* lethality in the F1 generation but also caused larval arrest in the P0 generation, a phenotype that is much more severe than that of wild-type N2 animals upon *lir-1* RNAi (22, 36). The extra Eri phenotype caused by the GFP::NRDE-3(ggIS1, bombardment) transgene prompted us to reinvestigate the expression and function of NRDE-3 under endogenous expression levels (39, 40).

We used CRISPR/Cas9 gene editing technology to knock in a 3xFLAG::GFP tag onto the N-terminus of the endogenous gene locus of NRDE-3 (Fig. 1A) and confirmed its copy number by real-time PCR (Fig. S1A). The response of GFP::NRDE-3(ust574, KI) animals to *lir-1* RNAi was similar to that of wild-type N2 animals (Fig. S1B), suggesting that the GFP::NRDE-3(ust574, KI) transgene largely recapitulated the endogenous functions of NRDE-3 without causing an extra Eri phenotype, as did the GFP::NRDE-3(ggIS1, bombardment) transgene. In addition, *dpy-11* RNAi caused nuclear RNAi-dependent gene silencing in both the P0 and F1 generations (36). Single-copy GFP::NRDE-3(ust574, KI) transgenic animals exhibited a similar response to RNAi to that of wild-type N2 animals upon *dpy-11* RNAi (Fig. S1C).

The expression of the previous GFP::NRDE-3(ggIS1, bombardment) transgene was restricted to late embryos (>80 cell stages) and larval somatic cells (22), whereas the GFP::NRDE-3(ust574, KI) transgene was expressed in the germline, oocytes, early and late embryos and larval soma (Figs. 1B-D, S1D-E), which is consistent with recent research from other laboratories (39, 40) and suggests that the original GFP::NRDE-3(ggIS1, bombardment) transgene may have been subjected to gene silencing in the germline, oocytes and early embryos.

Then, we compared the small RNA binding of the *in situ* GFP::NRDE-3(ust574, KI)-associated siRNAs with that of the ectopically expressed GFP::NRDE-3(ggIS1, bombardment). NRDE-3 was immunoprecipitated from the embryo lysates of the two strains, and the associated small RNAs were subjected to deep sequencing via a 5’-end phosphate-independent method(22, 41). In both strains, the NRDE-3-associated embryo-specific siRNAs were 22 nt in length, and the majority started with G at their 5′ ends, which is consistent with the properties of 22G RNAs (Fig. S2A). Approximately 79% of NRDE-3-associated 22G RNAs target protein-coding genes. We selected potential NRDE-3-specific targets that had greater than 25 reads per million small RNAs and identified 602 GFP::NRDE-3(ust574, KI) targets. The ectopically expressed GFP::NRDE-3(ggIS1, bombardment)-associated siRNA targets were largely a subset of the GFP::NRDE-3(ust574, KI) targets (Fig. S2B), which is consistent with the expression stage of both GFP::NRDE-3 strains in embryos.

The embryo-specific GFP::NRDE-3(ust574, KI) targets exhibited extensive overlap with those of HRDE-1 (Fig. S2C), the other germline-expressed nuclear Argonaute protein (42). The NRDE-3 embryo-specific targets also showed a large overlap with *glp-4-*dependent genes (Fig. S2D) (42), suggesting that NRDE-3 siRNAs may target germline-enriched genes. The WAGO-1 and WAGO-4 targets exhibited modest overlap with the NRDE-3 targets (Figs. S2E-F). However, CSR-1 targets exhibited only a slight overlap with NRDE-3 targets (42) (Fig. S2G).

Overall, GFP::NRDE-3(ust574, KI) was expressed in the germline, oocytes, early and late embryos, and larval somatic cells.

## The nuclear localization of NRDE-3 depends on RNA-dependent RNA polymerases

Previous work has shown that NRDE-3 is localized to the nucleus or nucleolus in somatic cells, depending on the presence of 22G RNAs (22). Consistent with these findings, mutations in *eri-1* and *ergo-1*, which are required for the biogenesis of endogenous siRNAs (22, 43), decreased the nuclear localization of NRDE-3 in the soma, as shown by both GFP::NRDE-3(ggIS1, bombardment) (22) and GFP::NRDE-3(ust574, KI) (Fig. 1B).

Interestingly, mutations in *eri-1* and *ergo-1* did not significantly deplete the nuclear accumulation of NRDE-3 in the germline, oocytes or early embryos. NRDE-3 remained in the nucleus in the mutants (Figs. 1C and 1D), suggesting that NRDE-3 may bind alternative cohorts of siRNAs in cells even when *eri-1-* and *ergo-1-*dependent siRNAs are depleted. The *C. elegans* genome encodes four RNA-dependent RNA polymerases, namely, RRF-1, RRF-2, RRF-3 and EGO-1(44, 45). Among them, RRF-1 localizes to Mutator foci, where it synthesizes WAGO class 22G RNAs using pUGylated RNA fragments as templates (46, 47). EGO-1 is a germline-specific RdRP that is essential for viability and synthesizes CSR-1 class 22G RNAs in E granules (18). We generated a *degron::ego-1* strain by the CRISPR/Cas9 method to degrade EGO-1 proteins upon treatment with IAA (48). The function of RRF-2 remains unclear.

In both the *rrf-1;rrf-2* double mutant and the *degron::ego-1(+IAA)* mutant, NRDE-3 was enriched in the nucleus in the germline, oocytes and early embryos (Figs. 1C-D). However, in the *rrf-1;rrf-2;degron::ego-1(+IAA)* triple mutant, NRDE-3 accumulated in the cytoplasm in the germline, oocytes and early embryos at 20°C (Figs. 1C-D). DRH-3 interacts with EGO-1, is enriched in both the E-granule and mutator foci and is required for both RRF-1- and EGO-1-dependent 22G RNA biogenesis (49). In the *drh-3* mutant, NRDE-3 accumulates in the cytoplasm. The depletion of NRDE-3 from the nucleus in the *rrf-1;rrf-2;degron::ego-1(+IAA)* triple mutant and in the *drh-3* mutant suggested that NRDE-3 associated with 22G RNAs generated by both RRF-1- and EGO-1-dependent production machineries.

### NRDE-3 accumulated in the perinuclear foci in embryos

Interestingly, although *eri-1* and *ergo-1* mutation did not abolish the nuclear accumulation of NRDE-3 in the germline and early embryos, we noticed that GFP::NRDE-3 was able to accumulate to the distinct perinuclear foci in embryos in *eri-1*(*-*) and *ergo-1*(*-*) animals (Figs. 2A-B). In *drh-3* mutant, although NRDE-3 was depleted from the nucleus and accumulated in the cytoplasm (Figs. 1C-D), we also observed the perinuclear foci-accumulation of NRDE-3 in embryos (Figs. 2A-B). However, no perinuclear foci accumulation of NRDE-3 were identified in any of the three mutant background: *degron::ego-1(+IAA)*, *rrf-1;rrf-2* double mutant*; rrf-1;rrf-2;degron::ego-1(+IAA)* triple mutant.

**Figure 2.**
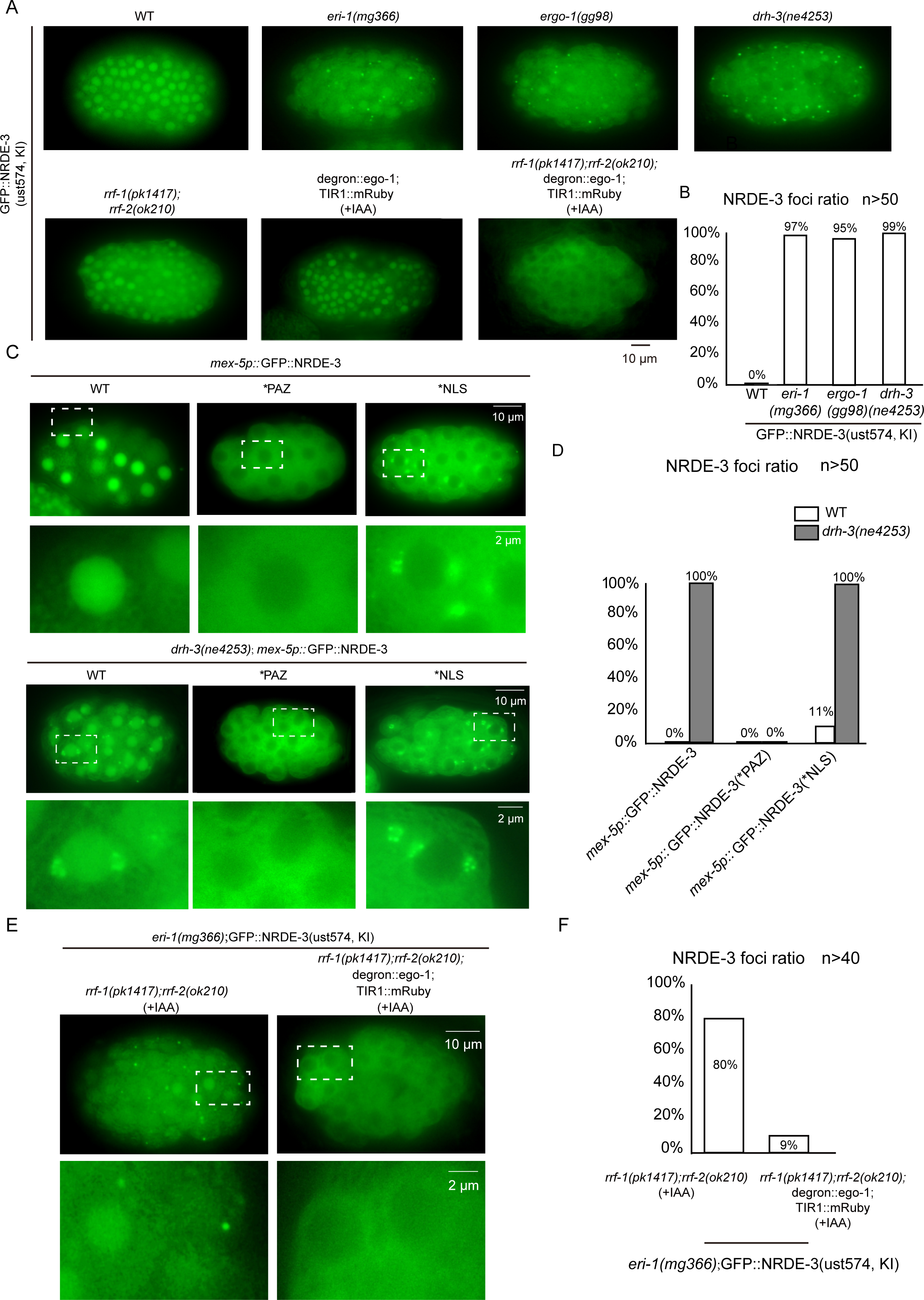
NRDE-3 accumulates in the perinuclear foci in embryos. (A) Fluorescence microscopy images of GFP::NRDE-3(ust574, KI) in late embryos in the indicated genotypes. (B) The bar graph quantifies the percentage of NRDE-3 foci in embryos. An embryo containing at least one NRDE-3 foci was counted as positive. At least 50 embryos were imaged for each genotype. (C) Fluorescence microscopy images of *mex-5* promoter-driven GFP::NRDE-3, GFP::NRDE-3(*PAZ) and GFP::NRDE-3(*NLS) in embryonic stages in indicated background. *PAZ, (Y463A, Y464A); *NLS, (K80A, R81A, K82A). (D) The bar graph quantifies the percentage of NRDE-3 foci-positive embryos. The embryo with one or more NRDE-3 foci in an embryo was defined as positive. At least 50 embryos were imaged for each condition. (E) Fluorescence microscopy images of *eri-1(mg366);*GFP::NRDE-3(ust574, KI) in embryos of the indicated genotypes. In *rrf-1;rrf-2* background, synchronized embryos were grown on NGM plates seeded with OP50 and 1 mM IAA.. For *rrf-1;rrf-2;degron::ego-1* animals, synchronized animals were grown on OP50 and at L4 stage shifted to NGM plates containing 1 mM IAA. Pictures were taken 48 hours later after IAA treatment. (F) The bar graph quantifies the percentage of NRDE-3 foci. An embryo containing at least one NRDE-3 foci was counted as positive. n>40.

To facilitate fluorescence detection of the perinuclear foci-enriched NRDE-3, we generated a much brighter GFP-NRDE-3 transgene using the *mex-5* promoter. The *mex-5* promoter-driven single-copy *mex-5p::3xFLAG::GFP::NRDE-3* transgene was inserted into the ttTi5605 site on chromosome II via CRISPR/Cas9 method. Similarly, the *mex-5* promoter-driven GFP::NRDE-3 was strongly enriched in the perinuclear foci in *drh-3* background (Figs. 2C-D).

The nuclear localization of NRDE-3 requires its ability to bind 22G RNAs (22). We tested whether the perinuclear foci-accumulation of NRDE-3 also requires it’s the presence of siRNA binding. PAZ domain is required for Arognaute proteins, including NRDE-3, to bind small RNAs (22). We genearated a *mex-5p::3xFLAG::GFP::NRDE-3(*PAZ, Y463A, Y464A)* mutant transgene (Fig. S3A). Consistent with previous reports (22), *NRDE-3(*PAZ, Y463A, Y464A)* protein accumulated in the cytoplasm as well. Notably, we did not observe the perinuclear foci accumulation of GFP::NRDE-3(*PAZ) in *drh-3* mutant, suggesting that small RNA binding is essential for NRDE-3 to enrich in the perinuclear foci (Figs. 2C-D).

The nuclear localization signal (NLS) sequence is important for NRDE-3 to translocate to the nucleus upon binding to siRNAs (22). The mutation in NLS(K80A, R81A, K82A) in NRDE-3 abrogated the nuclear accumulation of NRDE-3 and prohibited its ability to rescue nuclear RNAi defects in *nrde-3* animals (22). We generated *mex-5p::GFP::NRDE-3(*NLS, K80A, R81A, K82A, KI)* transgenic animals (Fig. S3A) and observed the depletion from the nucleus in both wild type N2 background and *drh-3* mutant embryos. Strikingly, in wild type N2 animals, approximately 11% early embryos revealed the perinuclear foci accumulation of NRDE-3(*NLS). In addition, the percentage increased to approximately 100% in *drh-3* mutants (Figs. 2C-D).

In the nucleus, siRNAs guide NRDE-3 to targeted pre-mRNAs and further recruit NRDE-2 to induce epigenetic modifications and gene silencing (23). However, we did not observe the perinuclear foci accumulation of NRDE-2 in both wild type N2 background and *drh-3* mutant animals (Fig. S3B), suggesting that NRDE-3 may target the nucleic acids in the perinuclear foci independent of other downstream NRDE factors.

We also tested whether other germline Argonaute proteins could accumulate in the perinuclear foci in embryos. However, neither HRDE-1 nor WAGO-4 showed the perinuclear foci accumulation in both wild type N2 background and *drh-3* mutant animals (Fig. S3C).

To further confirm that the presence of 22G siRNAs contributes to the perinuclear foci accumulation of NRDE-3, we investigated the NRDE-3 foci in RdRP mutant animals. We introduced *rrf-1;rrf-2;degron::ego-1(+IAA)* into *eri-1*(*-*) animals (Fig. 2D). The mutation of RRF-1, RRF-2 and EGO-1 together may deplete most, if not all, of the endogenous 22G RNAs (18). In the *rrf-1;rrf-2* double mutant, NRDE-3 still accumulated to distinct perinuclear foci in *eri-1*(*-*) animals (Figs. 2D-E). However, in *rrf-1;rrf-2;degron::ego-1(+IAA)* triple mutant, the perinuclear foci accumulation of NRDE-3 was drastically reduced. Interestingly, the perinuclear foci accumulation of NRDE-3 is temperature sensitive. Growing animals at 25℃ for 12 hours could deplete the perinuclear foci-enriched NRDE-3 (Figs. S4A-D).

Therefore, NRDE-3 could accumulate in distinct perinuclear foci in embryos, which depends on the binding of siRNAs.

### NRDE-3 accumulates in peri-centrosomal foci

We noticed that NRDE-3 accumulated in either one perinuclear focus or two perinuclear foci which were on the opposite side of the nucleus, in *eri-1, ergo-1* and *drh-3* animals. Therefore, we suspected that the focus is the centrosome or at least in peri-centrosomal region. To test this idea, we generated a number of tagRFP- and mCherry-tagged centriole proteins and pericentriolar material (PCM) and examined their relative subcellular localization with GFP::NRDE-3.

SAS-4, SAS-5 and SAS-6 are components of the centriole, AIR-1 is a component of the pericentriolar material, γ-tubulin is required for both centriole and PCM (1). In *eri-1, ergo-1* and *drh-3* mutants, GFP::NRDE-3 consistently accumulated in foci neighboring to the γ-tubulin, AIR-1, SAS-4, SAS-5 and SAS-6-marked centrosome (Figs. 3A, S5A-B). *tbb-2* encodes a *C. elegans* β-tubulin, which is an important part of the centrosome center-centriole, and also a component of the centrosome-based spindle assembly process in the early embryo (1). We generated single-copy *mex-5* promoter-driven tagRFP-tagged TBB-2 transgenic strains (termed tagRPF::TBB-2). NRDE-3 foci accumulated neighboring the microtubule marker TBB-2 as well (Fig. 3B). Although shifting animals to 25℃ could deplete the peri-centrosomal foci-enriched NRDE-3, the temperature change did not significantly alter the TBB-2-marked centrosome (Fig. S6A). Therefore, we concluded that NRDE-3 may accumulate in the peri-centrosomal foci.

**Figure 3.**
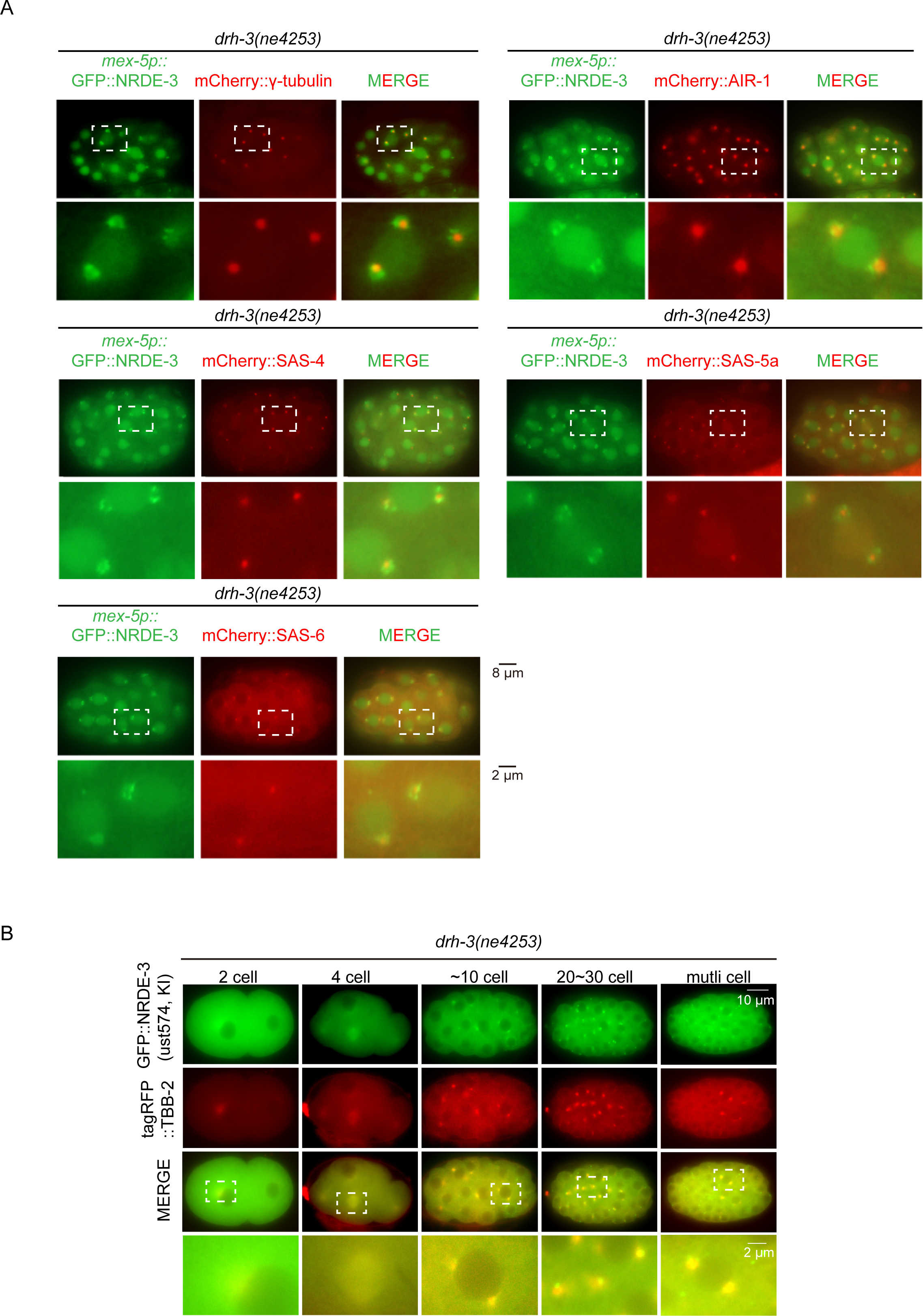
NRDE-3 accumulates in the peri-centrosomal foci. (A) Images of *mex-5p::* GFP::NRDE-3 with indicated mCherry-tagged centrosome proteins in *drh-3(ne4253)* background. (B) Images of GFP::NRDE-3(ust574, KI) with tagRFP::TBB-2 in embryos in *drh-3(ne4253)* background.

CSR-1 is the only Argonaute protein essential for fertility and forms diverse granules in embryos (39), yet we failed to detect the colocalization of CSR-1 with γ-tubulin and AIR-1-marked PCM (Fig. S6B).

### NRDE-3 accumulates in the peri-centrosomal foci and spindle during the cell cycle

The centrosome is highly dynamic and required for faithful cell division during the cell cycle (1). To investigate whether NRDE-3 also accumulates in centrosome in a cell cycle-dependent manner during embryogenesis, we used tagRFP::TBB-2 as centrosome marker and BFP::H2B as chromatin marker and quantified the proportion of NRDE-3 foci at different cell cycle phases. NRDE-3 was significantly enriched at the peri-centrosomal region throughout all mitotic phases (Figs. 4A-B). Especially, NRDE-3 also colocalized with the spindle during cell division. However, the NRDE-3(*PAZ) mutant did not exhibit centrosome and spindle enrichment, suggesting that siRNA binding is essential for NRDE-3 to accumulate in the peri-centrosomal region and spindle structure (Figs. S7A-B). The NRDE-3(*NLS) also revealed enrichment in the peri-centrosomal region and spindle during the cell cycle in wild type N2 background animals. The mutation in *drh-3* strongly enhanced the centrosome and spindle accumulation of NRDE-3(*NLS) (Figs. S8A-B). Therefore, we conclude that NRDE-3 could accumulate in the peri-centrosomal region and spindle during the cell cycle, which depends on the binding of siRNAs.

**Figure 4.**
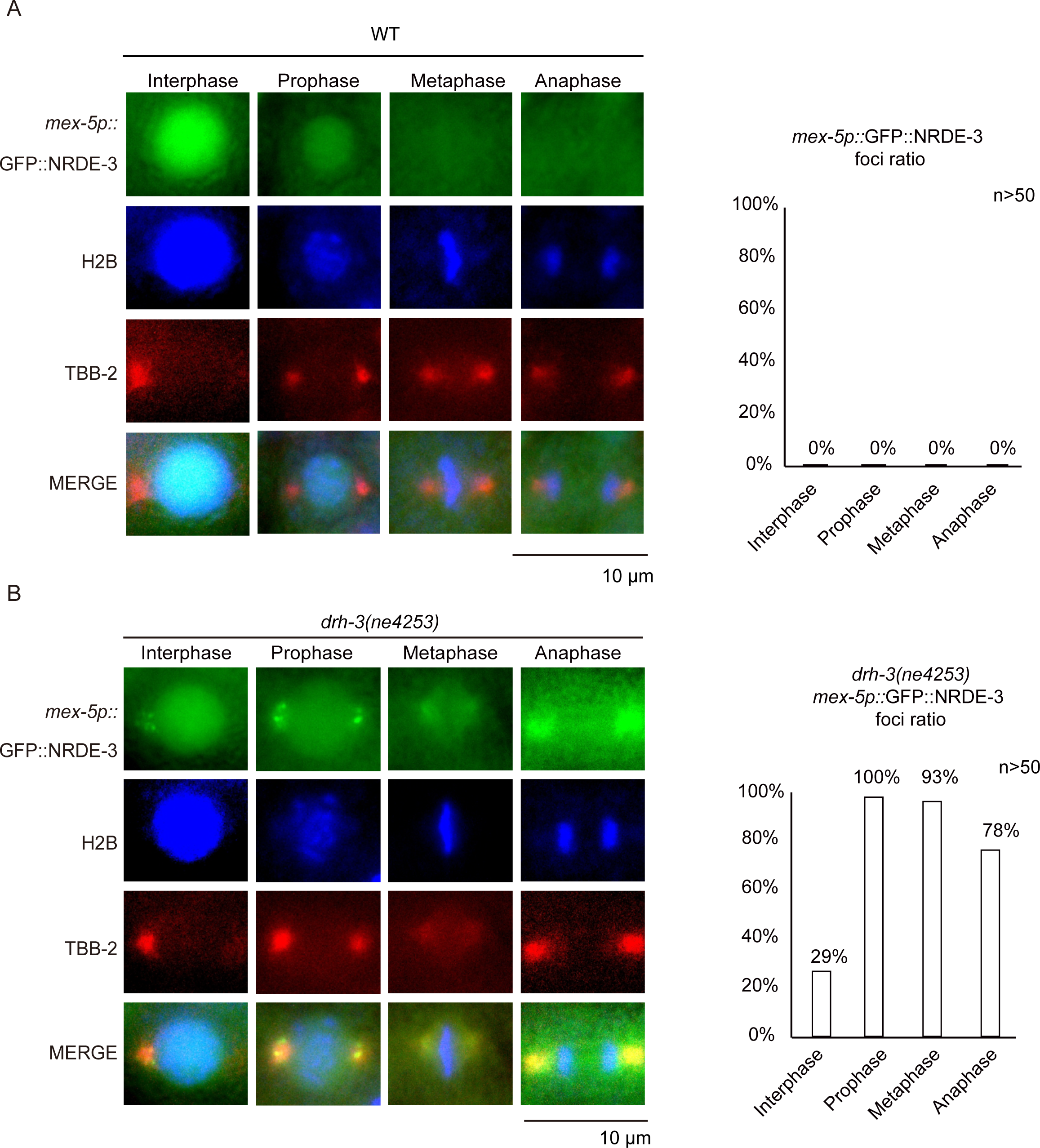
NRDE-3 accumulates in the peri-centrosomal foci and spindle during the cell cycle. (A, B) Left: Fluorescence microscopy images of indicated animals in interphase and mitosis. Right: Bar graph depicting the percentage of peri-centrosomal localization of NRDE-3 foci in each phase. Embryos at 10-30 cell stage were selected for quantification. For each embryo, each cell was assigned to different mitosis phases using BFP::H2B as a marker. We defined that one or more NRDE-3 foci in one cell as positive and then counted the percentage of positive cells in each phase, at least 50 cells were quantified for each phase.

### Identification of peri-centrosomal enriched siRNAs in *C. elegans*

The small size of the centrosome and the shell of embryos hindered us from biochemically purifying centrosomes of *C. elegans*. To specifically identify peri-centrosomal-enriched siRNAs, we sought to compare NRDE-3(*NLS)- and TBB-2-associated siRNAs in *drh-3* animals vs wild type animals and focused on the upregulated siRNAs, because *drh-3* mutation significantly increased the peri-centrosomal accumulation of NRDE-3 (Fig. 5A).

**Figure 5.**
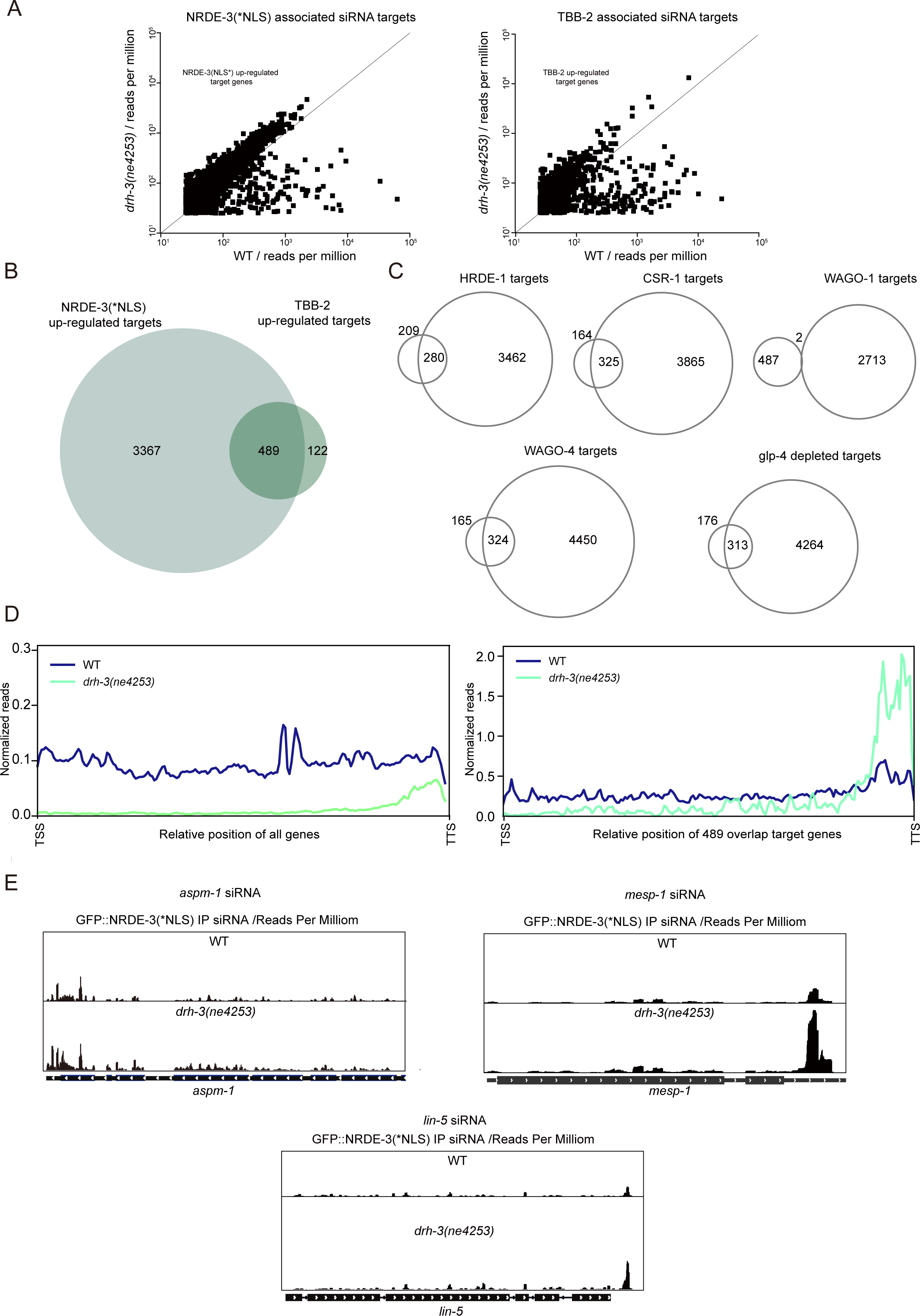
Identification of peri-centrosomal-enriched siRNAs in *C. elegans.* (A) Scatterplots showing gene-by-gene comparisons of NRDE-3(*NLS)- and TBB-2-associated small RNA targets reads between *drh-3(ne4253)* and wild-type animals. Each dot represents a gene. Cutoff > 25 reads per million. (B) Venn diagrams showing the overlap between upregulated siRNAs in the gene sets of NRDE-3(*NLS) (*drh-3*(*-*) vs wild type) and TBB-2 (*drh-3*(*-*) vs wild type). *drh-3(ne4253)*/wild type fold change >1.3. The overlap is annotated as potentially peri-centrosomal-enriched siRNAs. (C) Venn diagrams showing comparisons between the 489 overlap targets and other known siRNA categories. (D) Metaprofile analysis showing the distribution of normalized 22G RNA reads (RPM) along all genes (left) and the 489 overlap genes (right) in the indicated animals. (E) The distribution of normalized NRDE-3(*NLS)-associated small RNA reads across *aspm-1, mesp-1* and *lin-5* genomic loci in wild type and *drh-3(ne4253)* animals.

We immunoprecipitated NRDE-3 from NRDE-3(*NLS) strain and TBB-2 from tagRFP::TBB-2;NRDE-3(*NLS) strain at embryonic stage in both *drh-3(+)* and *drh-3*(*-*) animals. The associated siRNAs were isolated and subjected to deep sequencing in a 5’-end phosphate-independent manner. Approximately 90% of NRDE-3(*NLS)-associated small RNAs targeted protein-coding genes in both wild type background and *drh-3* animals. For TBB-2-immunoprecipitated small RNAs, the vast majority were miRNAs and siRNAs targeting protein-coding genes. The upregulated siRNAs in the gene sets of NRDE-3(*NLS) (*drh-3*(*-*) vs wild type) and TBB-2 (*drh-3*(*-*) vs wild type) were subsequently compared and 489 shared target genes were identified, which were potentially peri-centrosomal-enriched siRNAs (Fig. 5B, Table S1). These target genes revealed large overlap with HERD-1, CSR-1 and WAGO-4 targets and *glp-4*-dependent targets, but minimal overlap with WAGO-1 targets (Fig. 5C).

The 22G RNAs generated by EGO-1 have been classed to E-class siRNAs (49). Then, E-class siRNAs were further classified to E-class 5’ siRNAs and E-class 3’ siRNAs, which predominantly mapped to the 5’ portion of the E-class genes and 3’ end, respectively. The siRNAs targeting the 3’ most of the E-class genes relied on EGO-1 and typically include the last exon of the E-class genes (49). Metagene analysis revealed that the peri-centrosomal-enriched siRNAs predominantly mapped to the 3’ portion of the target genes (Fig. 5D).

A number of mRNAs have been reported to localize in centrosomes in human cells, *Drosophila* and *C. elegans* (14, 15). Three of the genes, *aspm-1*, *mesp-1* and *lin-5*, were identified among the top candidates in the overlap 489 target gene set, revealed abundant siRNA reads and increased siRNA association in *drh-3*(*-*) mutant vs *drh-3(+)* animals (Fig. 5E). *aspm-1* mRNA was reported to be enriched on mitotic centrosomes across all phases of cell division in *Drosophila* to human cells (10, 14). Although *mesp-1* mRNA has not been reported to be enriched in the centrosome, MESP-1 functions to sort microtubules of mixed polarity into a configuration in which the minus ends are away from the chromosomes, enabling the formation of nascent poles (50). LIN-5 protein binds the dynein complex, localizes in the spindle and invovles in microtubule cytoskeleton organization(51). *lin-5* mRNA was previously shown to be enriched in centrosomes by FISH assay (52). We noticed the increase of siRNAs targeting the 3’ portion of the *aspm-1*, *mesp-1* and *lin-5* genes (Fig. 5E) as well as in other peri-centrosomal targets (Figs. S9), as shown by IGV.

### The integrity of the centrosome is required for the peri-centrosomal accumulation of NRDE-3

We then tested whether the peri-centrosomal accumulation of NRDE-3 depends on the integrity of centriole proteins and pericentriolar material (PCM). GFP::NRDE-3 animals were fed with bacteria expressing dsRNAs targeting centriole, pericentriolar material and their interactors and F1 embryos were examined. Most of these proteins are essential for worm fertility or embryogenesis.

In F1 embryos, knocking down both PCM components and centriole proteins not only abolished the centrosome structure, as shown by tagRFP::AIR-1 and tagRFP::SAS-4 markers but also depleted the peri-centrosomal foci localization of GFP::NRDE-3 (Figs. 6A-C, S10), suggesting that centrosome integrity is essential for the NRDE-3 accumulation in the peri-centrosomal foci.

**Figure 6.**
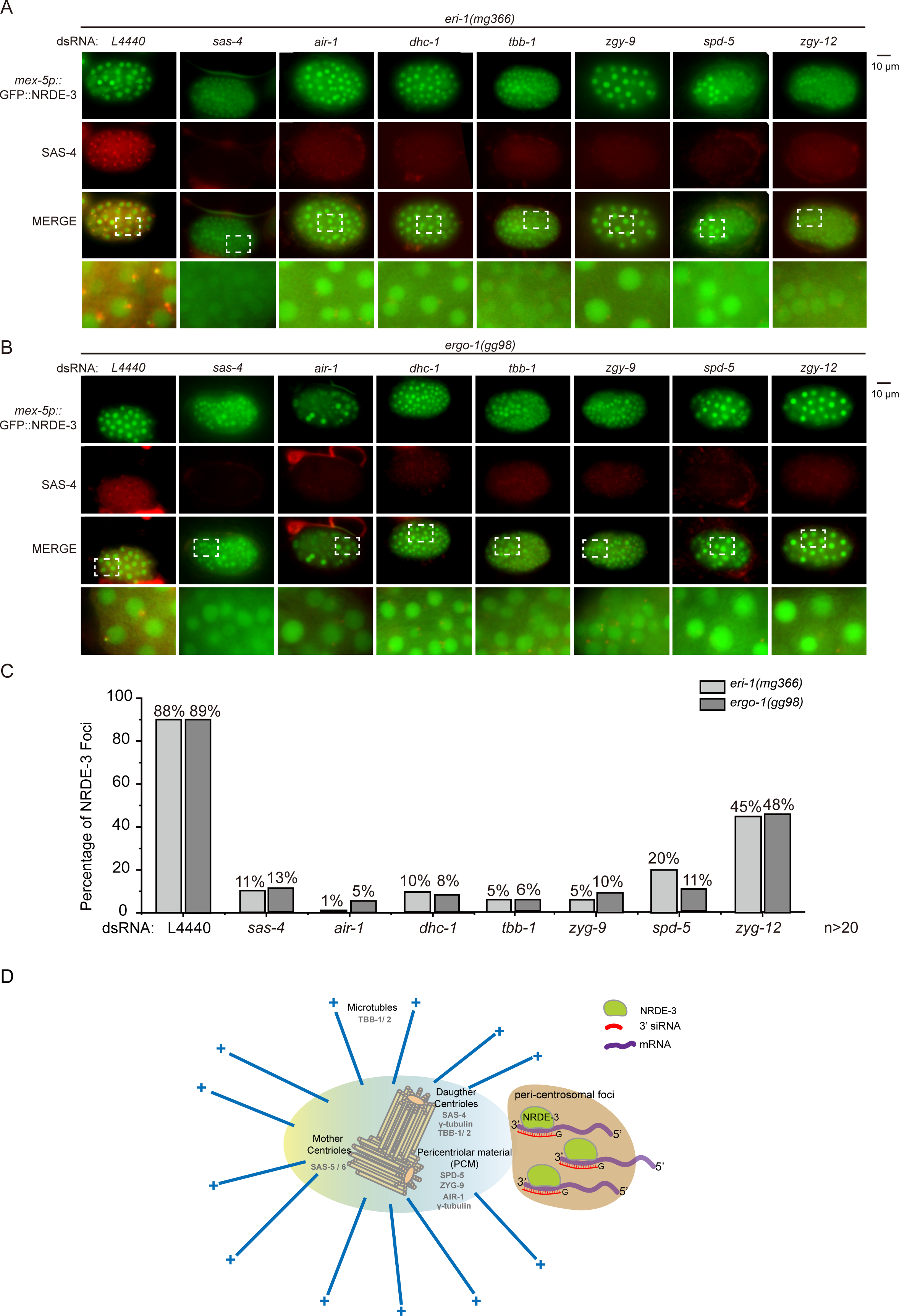
The integrity of the centrosome is required for the peri-centrosomal accumulation of NRDE-3. (A, B) Images of *eri-1*(*-*);GFP::NRDE-3;mCherry::SAS-4 (A) and *ergo-1*(*-*);GFP::NRDE-3;mCherry::SAS-4 (B) animals after feeding RNAi targeting indicated centriole, and PCM genes. Synchronized embryos were hatched and cultured on NGM plates for 41 hours and then at L4 stage transferred to RNAi plates. F1 embryos were imaged. (C) The bar graph quantified the percentage of NRDE-3 foci-positive embryos. The embryos with one or more NRDE-3 foci were counted as positive. At least 20 embryos were imaged for each RNAi experiment. (D) A working model for peri-centrosomal localization of NRDE-3/siRNAs in *C. elegans.* The nuclear Argonaute protein NRDE-3 accumulates in peri-centrosomal foci, a process depending on its ability of siRNA binding. NRDE-3-bound 22G RNAs map to the 3’ regions of the target genes.

## Discussion

Taken together, the results of this study revealed that the nuclear Argonaute protein NRDE-3 is expressed in the germline, oocytes, early and late embryos, and larval soma. Specifically, NRDE-3 can accumulate in the peri-centrosomal foci through a process dependent on its ability to bind siRNA and the integrity of centriole proteins and pericentriolar material (PCM) components (Fig. 6D), which implies that the pericentrosomal region may also be a platform for RNAi-mediated gene regulation. Moreover, this work suggested that the subcellular localization of NRDE-3 may be used to track the dynamics and transport of RNAs in distinct subcellular organelles.

### Centrosome-localized RNAs in *C. elegans*

Centrosomal RNAs (cnRNAs) have been identified in a number of model organisms through diverse purification and labeling approaches. However, the function and regulation of cnRNAs are largely mysterious. The lack of a genetic trackable system may have hindered the understanding of cnRNAs.

Several outstanding questions include how and why RNAs localize to the centrosome and how certain RNAs are selected to specifically accumulate in the centrosome. Specific RNA sequences or structural motifs may account for RNA localization via the recognition of distinct RNA-binding proteins (53). However, the key sequence motifs and particular binding partners of cnRNAs are still unknown. The identification of these sequences and binding partners will help to further understand the regulation and functions of cnRNAs. The localization of cnRNAs is cell cycle-dependent (3, 12, 13). It has been hypothesized that centrosomes are larger in interphase and that mRNAs are more likely to localize to interphase centrosomes (12, 54, 55). The mechanism of the spatiotemporal regulation of cnRNAs is ambiguous. Additionally, intrinsically disordered proteins and specific RNAs are important for the formation of membraneless organelles through phase separation processes, which may also contribute to the structural formation of centrosomes (56, 57). Identification of the RNA components in centrosomes and pericentrosomal regions will help to clarify the phase separation properties of centrosomes and their surrounding regions and how RNAs influence centrosome composition and biological consequences.

Recent literature has suggested that RNA localization is frequently coupled with translational control for a number of mRNAs (5, 58, 59). The local translation of cnRNAs is likewise required for normal centrosome function (5, 10, 14, 60). Consistently, our data suggested that *aspm-1*, *mesp-1* and *lin-5,* which have centrosome-related functions, likely enrich for their mRNAs in the peri-centrosomal foci. Further experiments, including MS2-GFP reporter system or single-molecular FISH, may help to confirm the subcellular localization of these mRNAs.

A number of Argonaute proteins exhibit slicer activity, via which small regulatory RNAs guide the Argonaute protein to complement nucleic acid sequences and slice targets. NRDE-3 is a worm-specific Argonaute protein that has no noticeable slicer residues but is able to inhibit transcription elongation and induce histone modification in the nucleus (22, 24, 61). Whether and how siRNAs direct NRDE-3 to pericentrosome-localized mRNAs and inhibit translation are intriguing questions.

### Centrosomes may participate in RNAi-mediated gene regulation

Due to technical limitations, the subcellular localization of small regulatory RNAs and their specialized functions have largely been neglected. With the development of various technologies, including the isolation of subcellular organelles and immunofluorescence localization analysis, an increasing number of studies have yielded broad insights into the subcellular localization and functions of small RNAs. Small RNAs have been shown to localize to mitochondria, secreted exosomes, the endoplasmic reticulum, stress granules, the processing body, germ granules, etc. (27, 28, 31–33). These organelle-localized small RNAs may mediate transcriptional regulation and modulate homeostasis for many biological and physiological processes, which are necessary for cell fitness. In organelles, AGO or PIWI proteins may be recruited to form the RISC complex to posttranscriptionally modulate gene expression (62–64). For example, Ago2 has been reported to interact with MitomiRs and regulate the mitochondrial transcriptome (65). In addition, Ago2 and Dicer in stress granules can regulate miRNA biogenesis (66). Some small regulatory RNAs may further exchange and traffic through different organelles to direct mutual regulation.

Here, we showed that the centrosome or pericentrosomal region may provide a platform for RNAi-mediated gene regulation. The Argonaute protein NRDE-3 can bind a certain class of siRNAs and accumulate in distinct peri-centrosomal foci, especially in *eri-1, ergo-1* and *drh-3* mutants. When endogenous 22G siRNAs were completely depleted in the RdRP triple mutants, the accumulation of peri-centrosomal foci of NRDE-3 was reduced.

The 22G RNAs generated by EGO-1 have been classified as E-class siRNAs (49). E-class siRNAs were further classified into E-class 5’ siRNAs and E-class 3’ siRNAs, which predominantly mapped to the 5’ portion of the E-class genes and 3’ end, respectively. siRNAs mapped to the 3*’* regions of CSR-1 class mRNAs may be produced by EGO-1 in the cytosol (49). Interestingly, the peri-centrosomal foci-localized NRDE-3 may preferentially bind 3’ siRNAs. DRH-3(ne4253) is a temperature-sensitive mutant that is required for the production of 5’ siRNAs of E-class siRNAs but not for 3’ siRNA accumulation (also see Fig. 5) (49, 67). In the *drh-3(ne4253)* mutant, NRDE-3 may therefore preferentially bind 3’ siRNAs and accumulate in the peri-centrosomal foci. Whether peri-centrosome-localized siRNAs contribute to centrosome homeostasis during mitosis and gene expression regulation in an RNAi-mediated manner requires further investigation.

### NRDE-3 was used as a reporter to track the subcellular localization of RNAs *in vivo*

The introduction of green fluorescent protein (GFP) has revolutionized the investigation of cellular proteins *in vivo* (68, 69). However, although a great number of methods have been developed to assay RNAs *in vitro*, it is still difficult to directly visualize RNAs in live cells. Many methods, including FISH and fluorophore aptamer-based RNA imaging, frequently require cell fixation and are not applicable in live cells to spatiotemporally track RNAs. The MS2-MCP system has been successfully used in living organisms, yet long repetitive MS2 sequences may interfere with the normal function of RNAs. A series of small, monomeric and stable fluorescent RNAs with large Stokes shifts, including Pepper, Clivias and Okra, have been developed to enable simple and robust imaging of RNAs with minimal perturbation of the target RNA’s localization and functionality (70–73). The development of SunTag technology has enabled the detection of cotranslational centrosomal RNA as well (14, 60).

NRDE-3 is an Argonaute protein that associates with 22G RNAs and usually translocates to the nucleus to target pre-mRNAs containing complementary nucleic acid sequences. Thus, the nuclear accumulation of NRDE-3 has been used to visualize tissue-specific gene expression (22). NRDE-3 has also been used to track transcription dynamics in live *C. elegans* (29, 30). Here, by using fluorescence-tagged NRDE-3, we first showed that siRNAs could accumulate in the eri-centrosomal foci, which suggests that a particular cohort of siRNAs and mRNAs may accumulate and play certain mysterious roles in centrosome and spindle regulation. With the aid of proximity RNA labeling technology, NRDE-3 may also be used to identify the transcriptome in particular subcellular organelles.

## Materials and methods

### Strains

The Bristol strain N2 was used as the standard wild-type strain. All strains were grown at 20°C unless otherwise specified. For heat stress treatment, worms were cultured at 25°C for 12 hours. The strains used in this study are listed in Supplementary Table S2.

### Construction of plasmids

All plasmids were generated through the recombinational cloning of PCR-amplified fragments. A ClonExpress MultiS One Step Cloning Kit (Vazyme) was used to connect these fragments to the vector. The construction process is described in detail in the supplemental materials and methods.

### Gene editing by the CRISPR/Cas9 method

For the *in situ* knock-in transgene 3xFLAG::GFP::NRDE-3(ust574, KI), the injection mixture contained pDD162 (50 ng/µl), the NRDE-3 repair plasmid (50 ng/µl), pCFJ90 (5 ng/µl), and three sgRNAs (30 ng/µl). The mixture was injected into young adult N2 animals. The transgenes were integrated by the CRISPR/Cas9 system.

For the *in situ* knock-in *degron::ego-1(ust614)* transgene, the injection mixture contained PDD162 (50 ng/µl), the *degron::ego-1* repair plasmid (50 ng/µl), pCFJ90 (5 ng/µl), and three sgRNAs (30 ng/µl). The mixture was injected into young adult CA1199 [*sun-1p*::TIR1::mRuby::sun-1 3’UTR] animals. The transgenes were integrated via the CRISPR/Cas9 method.

For ectopic transgenes, the injection mixture contained pDD162 (50 ng/µl), an ectopic transgene homology-directed repair plasmid (50 ng/µl), pCFJ90 (5 ng/µl), and three sgRNAs (30 ng/µl). The mixture was injected into young adult N2 animals. The transgenes were integrated by the modified counterselection (cs)-CRISPR method (74, 75).

The sgRNAs used in this study for transgene construction are listed in Supplementary Table S3.

### Imaging and quantification

Images were collected on a Leica DM4 B microscope.

### RNAi

RNAi experiments were performed at 20°C by placing synchronized embryos on feeding plates as previously described (76). HT115 bacteria expressing the empty vector L4440 (a gift from A. Fire) were used as controls. Bacterial clones expressing dsRNAs were obtained from the Ahringer RNAi library and sequenced to verify their identity. Images were collected using a Leica DM4 B microscope.

### Quantitative real-time PCR

Worm samples from the indicated animals were collected and digested at 65°C with proteinase K to extract genomic DNA. qRT‒PCR was performed using a MyIQ2 real-time PCR machine (Bio-Rad) with AceQ SYBR Green Master mix (Vazyme). The primers used for qRT‒PCR are listed in Supplementary Table S4.

### Auxin treatment

Unless otherwise indicated, auxin treatment was performed by transferring worms to bacteria-seeded plates containing auxin. The natural auxin indole-3-acetic acid (IAA) was purchased from Sigma‒Aldrich (#12886). A 400 mM stock solution in ethanol was prepared and stored at 4°C for up to one month. IAA was diluted into NGM agar and cooled to approximately 50°C before being added to the plates. Fresh OP50 spreading plates. The plates were left at room temperature for 1-2 days to allow bacterial lawn growth. For all IAA treatments, 0.25% ethanol was used as a control.

For all strains containing *degron::ego-1(ust614); sun-1p::*TIR1::mRuby::sun-13’UTR, animals exposed to IAA treatment exhibited degradation of degron-tagged EGO-1 in the germline, leading to embryonic arrest in F1 embryos. For *degron::ego-1(ust614)*;*sun-1p::*TIR1::mRuby::sun-13’UTR;GFP::NRDE-3(ust574, KI) and *eri-1(mg366);rrf-1(pk1417);rrf-2(ok210);*GFP::NRDE-3(ust574, KI), synchronized embryos were exposed onto NGM plates seeded with OP50 and containing 1 mM IAA. For *rrf-1(pk1417);rrf-2(ok210);degron::ego-1(ust614);sun-1p::*TIR1::mRuby::sun-13’UTR; GFP::NRDE-3(ust574, KI) and *eri-1(mg366);rrf-1(pk1417);rrf-2(ok210;degron::ego-1(ust614);sun-1p::*TIR1::mRuby::sun-13’UTR;GFP::NRDE-3(ust574, KI), synchronized animals were grown on OP50 on NGM plates and at the L4 stage, they were transferred to NGM plates containing 1 mM IAA. Pictures were taken 48 hours after IAA treatment.

### RNA immunoprecipitation (RIP) assay

3xFLAG::GFP::NRDE-3(ust574, KI), *nrde-3p::3Xflag::gfp::nrde-3(ggIS1, bombardment)* and NRDE-3(*NLS)-associated siRNAs were isolated from embryo lysates. The embryos were cryogrolded in lysis buffer (20 mM Tris-HCl, pH 7.5, 200 mM NaCl, 2.5 mM MgCl_2_, 0.5% NP-40, and 10% glycerol) supplemented with proteinase inhibitor tablets (Roche). The lysate was precleared with protein G-agarose beads (Roche) and then incubated with anti-FLAG M2 magnetic beads (Sigma #M8823). The beads were washed extensively and eluted with 3xFLAG peptide (Sigma #F4799). The eluates were incubated with DNase I followed by TRIzol purification and isopropanol precipitation. The precipitated RNA was treated with RNA 5′ polyphosphatase (Biosearch) and re-extracted with TRIzol reagent.

TBB-2-associated siRNAs were isolated from *mex5p*::3xHA::tagRFP::TBB-2;GFP::NRDE-3(*NLS) embryo lysates and incubated with anti-HA magnetic beads (Thermo Scientific #88836). The beads were washed extensively and eluted with HA peptide (Thermo Scientific #26184).

### siRNA deep sequencing

NRDE-3- and TBB-2-bound siRNAs were subjected to deep sequencing using an Illumina platform by Novogene Bioinformatics Technology Co., Ltd., for library preparation and sequencing. Briefly, 3’ and 5’ adaptors were ligated to the 3’ and 5’ ends of small RNAs, respectively. After purification and size selection, libraries with insertions between 18∼30 bp were subjected to deep sequencing.

The Illumina-generated raw reads were first filtered to remove adaptors, low-quality tags, and contaminants to obtain clean reads at BGI Shenzhen. Clean reads ranging from 18 to 30 nt were mapped to the unmasked *C. elegans* genome and the transcriptome assembly WS243, respectively, using Bowtie2 with default parameters. The number of reads targeting each transcript was counted using custom Perl scripts and displayed by IGV.

### Metagene analysis

The metagene profiles were generated according to a method described previously (77). The BigWig files were generated using the Snakemake workflow (https://gitlab.Pasteur.fr/bli/bioinfo_utils). Briefly, the 3′ adapters and 5′ adapters were trimmed from the raw reads using Cutadapt v.1.18 with the following parameters: -a AGATCGGAAGAGCACACGTCT -g GTTCAGAGTTCTACAGTCCGACGATC – discard-untrimmed. After adapter trimming, the reads containing 18 to 26 nt were selected using Bioawk. The selected 18–26 nucleotide reads were aligned to the *C. elegans* genome (ce11, *C. elegans* Sequencing Consortium WBcel235) using Bowtie2 v.2.3.4.3 with the following parameters: -L 6 -i S,1,0.8 -N 0. The resulting alignments were used to generate bigwig files with a custom bash script using BEDtools version v2.27.1, BEDOPS version 2.4.35, and bedGraphToBigWig version 4. Read counts in the bigwig file were normalized to million “nonstructural” mappers—that is, reads containing 18 to 26 nt and mapped to annotations not belonging to “structural” (tRNA, snRNA, snoRNA, rRNA, ncRNA) categories—and counted using featureCounts77 v.1.6.3. These bigwig files were used to generate “metaprofiles” files with a shell script.

### Statistics

The means and standard deviations of the results are presented in bar graphs with error bars. All experiments were conducted with independent *C. elegans* animals at the indicated number (N) of times. Statistical analysis was performed with two-tailed Student’s t tests.

## Data availability

The data that support this study are available from the corresponding author upon request. All the high-throughput data generated by this work have been deposited in the Genome Sequence Archive (Genomics, Proteomics & Bioinformatics 2021) (78) in National Genomics Data Center (Nucleic Acids Res 2022) (79), China National Center for Bioinformation / Beijing Institute of Genomics, Chinese Academy of Sciences (GSA: CRA017153) that are publicly accessible at https://ngdc.cncb.ac.cn/gsa/ CRA017153.

## Acknowledgments

We are grateful to members of the Guang laboratory and Dr. Carolyn Phillips’ laboratory for their comments. We are grateful to the International *C. elegans* Gene Knockout Consortium and the National Bioresource Project for providing the strains. Some strains were provided by the CGC, which is funded by the National Institutes of Health (NIH) Office of Research Infrastructure Programs (P40 OD010440).

## Author contributions

S.G. and C.Z. conceptualized the research; X.F., X.H., C.Z. and S.G. designed the research; Q.J., C.Z. and M.H. performed the research; Q.J., X.F., M.H., X.C., K.W., J.C., Y.K., X.S., and M.X. contributed new reagents; K.W. and C.Z. contributed analytic tools and performed bioinformatics analysis; and S.G. wrote the paper.

## Supplemental materials and methods

### Construction of plasmids

All plasmids were generated through the recombinational cloning of PCR-amplified fragments. A ClonExpress MultiS One Step Cloning Kit (Vazyme) was used to connect these fragments to the vector.

For the *in situ* knock-in transgene 3xFLAG::GFP::NRDE-3(ust574, KI), the 3xFLAG::GFP sequence was PCR amplified with the primers 5’-ATGGACTACAAAGACCATGACGGT-3’ and 5’-AGCTCCACCTCCACCTCC-3’ from YY178 genomic DNA. The homologous left arm (1.2 kb) was PCR amplified with the primers 5’-ATAACAATTTCACAGGGCCCCCGTCGATCAAGTTTGCCGG −3’ and 5’-ATAACAATTTCACAGGGCCCCCGTCGATCAAGTTTGCCGG −3’ from N2 genomic DNA. The homologous right arms (1.3 kb) were PCR amplified from N2 genomic DNA with the primers AAGGAGGTGGAGGTGGAGCTATGGATCTCCTAGACAAAGTAATG-3’ and CCAGTCACGACGTCACGTGAATCAGAGTAACCTCGTCGGG-3’. The backbone was PCR amplified with the primers 5’-CACGTGACGTCGTGACTGGG −3’ and 5’-GGGCCCTGTGAAATTGTTATCC −3’ from the plasmid pCFJ151.

For the *in situ* knock-in *degron::ego-1(ust614)* transgene, the homologous left arm (1.4 kb) was PCR amplified with the primers 5’-ATAACAATTTCACAGGGCCCCGTCCAAATCTTGGTTTCTGGC-3’ and 5’-CGGTATGATCTCACCGGTGGCCATCCCACAACTTGTGCCTTGGCCGGAGGTTTGGCTGGATCTTTAGGCATTGTTGCGAGGATTCGGGATA-3’ from N2 genomic DNA. The homologous right arms (1.5 kb) were PCR amplified with the primers 5’-CCACCGGTGAGATCATACCGGAAGAACGTGATGGTTTCCTGCCAAAAATCAAGCGGTGGCCCGGAGGCGGCGGCGTTCGTGAAGGGAGGTGGAGGTGGAGCTATGGGGGACGAAGGTTATCG-3’ and 5’-CCCAGTCACGACGTCACGTGGATCCTTAACCACTGCTCCTCTC −3’ from N2 genomic DNA. The degron sequence was added to the abovementioned PCR primers. The backbone was PCR amplified with the primers 5’-CACGTGACGTCGTGACTGGG −3’ and 5’-GGGCCCTGTGAAATTGTTATCC −3’ from the plasmid pCFJ151.

For the ectopic transgene *mex-5p*:: 3xFLAG::GFP::NRDE-3(*PAZ):tbb-2 UTR inserted in chr II, the *mex-5p*::3xFLAG::GFP::NRDE-3(*PAZ)::tbb-2 UTR was amplified with the primers 5’-CTTCCATCACAGAGGCTGCCTTACAACGGTACAACTATCGTCTC −3’ and 5’-GGCAGCCTCTGTGATGGAAGTGAGCTTC −3’ from shg750 genomic DNA. The vector fragment was the same as that used for the chr II vector above.

For the ectopic transgene *mex-5p*::GFP::NRDE-3(m)(*NLS):tbb-2 UTR inserted in chr II, the *mex5-p*:: 3xFLAG::GFP::NRDE-3(*NLS)::tbb-2 UTR was amplified with the primers 5’-GCAGCAGCGCCACTGGGAGGATGTGGGT −3’ and 5’-CCTCCCAGTGGCGCTGCTGCTGGATCGGGACCTGTGCGAT −3’ from 750 genomic DNA. The vector fragment was the same as that used for the chr II vector above.

For the ectopic transgene *mex-5p*::tagRFP::TBB-2 inserted in chr I, the MEX-5 promoter was amplified with the primers 5’-CCTCCCTGTCAATTCCCAAAATACAAATATCAGTTTTTAAAAAATTAAACCAT −3’ and 5’-TCTTCTCCCTTGGACACCATTCTCTGTCTGAAACATTCAATTG −3’ from N2 genomic DNA. The tagRFP sequence was PCR amplified with the primers 5’-ATGGTGTCCAAGGGAGAAGA-3’ and 5’-GTTTGAGCTTGTGCCCGAGC-3’ from yy967 genomic DNA. The coding sequence (CDS) region and 3′ untranslated region (UTR) sequence were PCR amplified with the primers 5’-GCTCGGGCACAAGCTCAACGGAGGTGGAGGTGGAGCTATGAGAGAGATCGTCCACGT −3’ and 5’-TTCAAAGAAATCGCCGACTTCCAACCACATATGTTTCTCTTAGGC −3’ fromN2 genomic DNA. The chr I vector fragment was PCR amplified with the primers 5’-AAGTCGGCGATTTCTTTGAAGTT-3’ and 5’-GTATTTTGGGAATTGACAGGGAGG-3’ from the plasmid pSG274.

For *mex-5p*::3xHA::tagRFP::TBB-2 inserted in chr I, *mex-5p*::3xHA::tagRFP::TBB-2 was PCR amplified with the primers 5’-TCCATATGACGTGCCGGACTATGCATACCCATACGATGTTCCAGATTACGCTTACCCATACGATGTTCCAGATTACGCTATGGTGTCCAAGGGAGAAGA −3’ and 5’-AGTCCGGCACGTCATATGGATACATTCTCTGTCTGAAACATTCAATTG-3’ from shg1494 genomic DNA. The vector fragment used was the same as that used for the chrI vector above.

For *mex-5p*::Cherry::TBG-1::tbb-2 utr inserted in chr I, the MEX-5 promoter was amplified with the primers 5’-CCTCCCTGTCAATTCCCAAAATACAAATATCAGTTTTTAAAAAATTAAACCAT −3’ and 5’-TCCTCTCCCTTGGAGACCATTCTCTGTCTGAAACATTCAATTG −3’ from N2 genomic DNA. The mCherry sequence was PCR amplified with the primers 5’-ATGGTCTCCAAGGGAGAGGA-3’ and 5’-ATAGCTCCACCTCCACCTCCCTTGTAAAGCTCATCCATTCCTCC-3’ from the SX2650 genomic DNA. The coding sequence (CDS) region sequence was PCR amplified with the primers 5’-GGAGGTGGAGGTGGAGCTATGTCCGGTACGGGTGCC-3’ and 5’-TGCTTGAAAGGATCTTGCATCTAAAGCCCTCTTGTCAGATAG-3’ from N2 genomic DNA. The tbb-2 untranslated region (UTR) was PCR amplified with the primers 5’-ATGCAAGATCCTTTCAAGCA-3’ and 5’-TTCAAAGAAATCGCCGACTTCCAACCACATATGTTTCTCTTAGGC-3’ from N2 genomic DNA. The vector fragment used was the same as that used for the chrI vector above.

For *mex-5p*::Cherry::AIR-1::tbb-2 utr inserted in chr I, the coding sequence (CDS) region sequence was PCR amplified with the primers 5’-AGGGAGGTGGAGGTGGAGCTATGAGCGGAAAGGAAAATACTGC-3’ and 5’-TGCTTGAAAGGATCTTGCATTTATTGATTGGCTGTAGAATTATTGCGAC-3’ from N2 genomic DNA. The mex-5p::Cherry::tbb-2 utr vector was PCR amplified with the primers 5’-ATGCAAGATCCTTTCAAGCA-3’ and 5’-ATAGCTCCACCTCCACCTCCCTTGTAAAGCTCATCCATTCCTCC-3’ from shg1064 genomic DNA.

For *mex-5p*::mCherry::SAS-4::tbb-2 utr inserted in chr I, the coding sequence (CDS) region sequence was PCR amplified with the primers 5’-AGGGAGGTGGAGGTGGAGCTATGGCTTCCGATGAAAATATCGG −3’ and 5’-TGCTTGAAAGGATCTTGCATTCATTTTTTCCACTGGAACAAAGTTG −3’ from N2 genomic DNA. The mex-5p::mCherry::tbb-2 utr backbone fragment is the same as that of *mex-5p*::mCherry::AIR-1::tbb-2 utr above.

For *mex-5p*::mCherry::SAS-5a::tbb-2 utr inserted in chr I, the coding sequence (CDS) region sequence was PCR amplified with the primers 5’-AGGGAGGTGGAGGTGGAGCTATGAATAATTACGACGACTTACCCT −3’ and 5’-TGCTTGAAAGGATCTTGCATTCATTTCCTGCGAGCGTATTTTTC −3’ from N2 genomic DNA. The mex-5p::mCherry::tbb-2 utr backbone fragment is the same as that of *mex-5p*::mCherry::AIR-1::tbb-2 utr above.

For *mex-5p*::mCherry::SAS-6::tbb-2 utr inserted in chr I, the coding sequence (CDS) region sequence was PCR amplified with the primers 5’-AGGGAGGTGGAGGTGGAGCTATGACTAGCAAAATTGCATTATTCG −3’ and 5’-TGCTTGAAAGGATCTTGCATTTATCGTTGAGCGGGTGGGG −3’ from N2 genomic DNA. The mex-5p::mCherry::tbb-2 utr backbone fragment is the same as that of *mex-5p*::mCherry::AIR-1::tbb-2 utr above.

For *mex-5p*::BFP::H2B::tbb-2 utr inserted in chr IV, the MEX-5 promoter was amplified with the primers 5’-CATCCCGTTAGAAACAATCTAAATATCAGTTTTTAAAAAATTAAACCAT −3’ and 5’-TCCTTAATAAGCTCTGACATTCTCTGTCTGAAACATTCAATTG −3’ from N2 genomic DNA. The BFP sequence was PCR amplified with the primers 5’-ATGTCAGAGCTTATTAAGGAGAATATG-3’ and 5’-ATAGCTCCACCTCCACCTCCATTAAGCTTGTGACCCAGTTTGC-3’ from shg1902 genomic DNA. The coding sequence (CDS) region and tbb-2 untranslated region (UTR) sequence were PCR amplified with the primers GGAGGTGGAGGTGGAGCTA-3 and GCTAACTTACATTTAGCTAGCCAACCACATATGTTTCTCTTAGGC-3 from shg366 genomic DNA. The chr IV vector fragment was PCR amplified with the primers 5’-CTAGCTAAATGTAAGTTAGCGACC −3’ and 5’-AGATTGTTTCTAACGGGATGC −3’ from the plasmid pCZGY2729.

---

**Figure S1.**
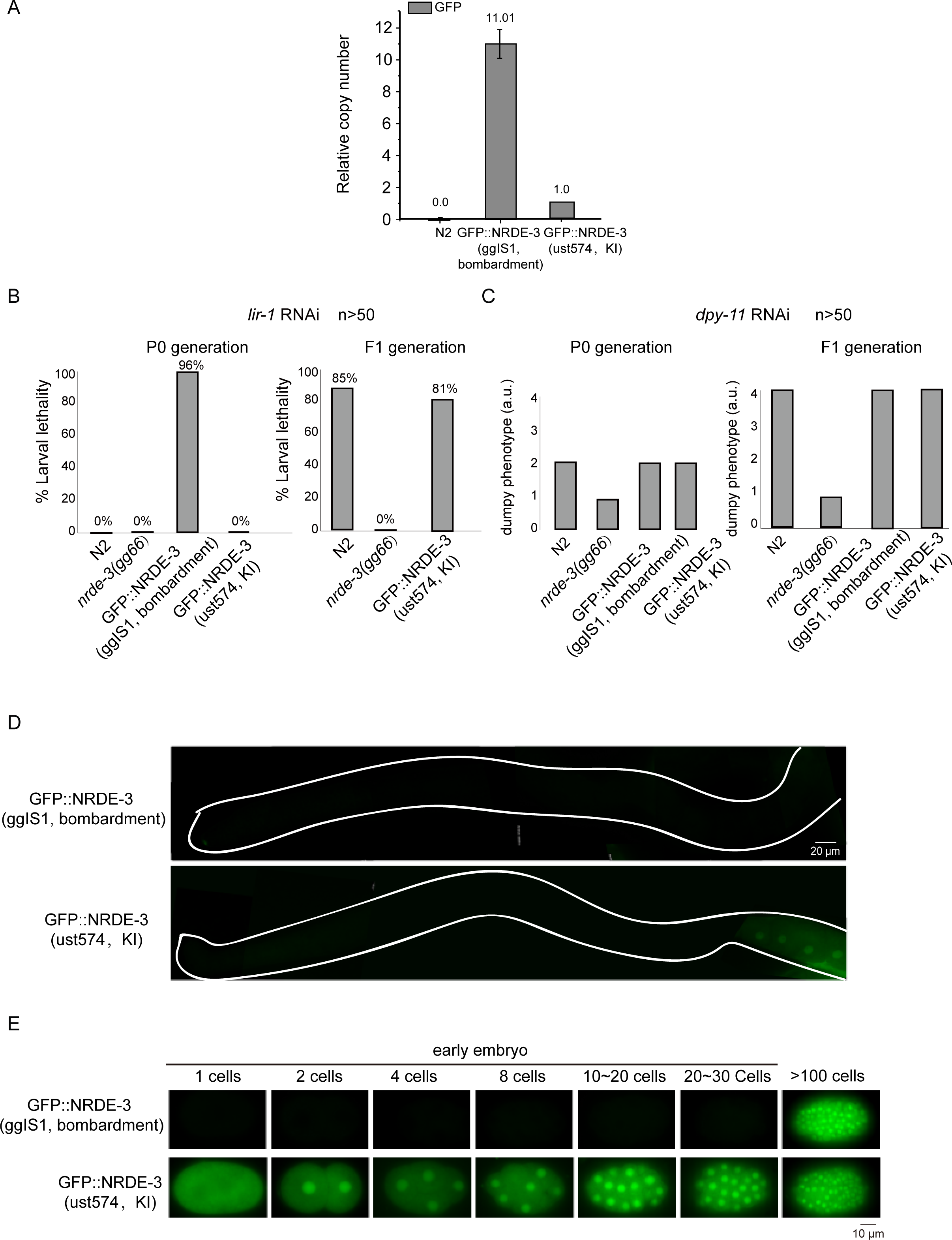
NRDE-3 is widely expressed in the germline, oocytes, early and late embryos and soma. (A) The copy number of GFP in the indicated animals was quantified by real-time PCR. *eft-3* genomic DNA was used as an internal control for normalization. Mean ± SD. n = 3. (B) The percentage of larval arrest after feeding RNAi targeting the *lir-1* gene in the indicated animals (n > 50). Left: P0 generation, Right: F1 generation. (C) The percentage of dumpy animals after *dyp-11* RNAi in the indicated groups (n > 50). Left: P0 generation, Right: F1 generation. (D) Fluorescence micrographs of dissected gravid adult germlines from the indicated animals. The germline outline is marked. (E) Fluorescence micrographs of embryos from the indicated animals.

**Figure S2.**
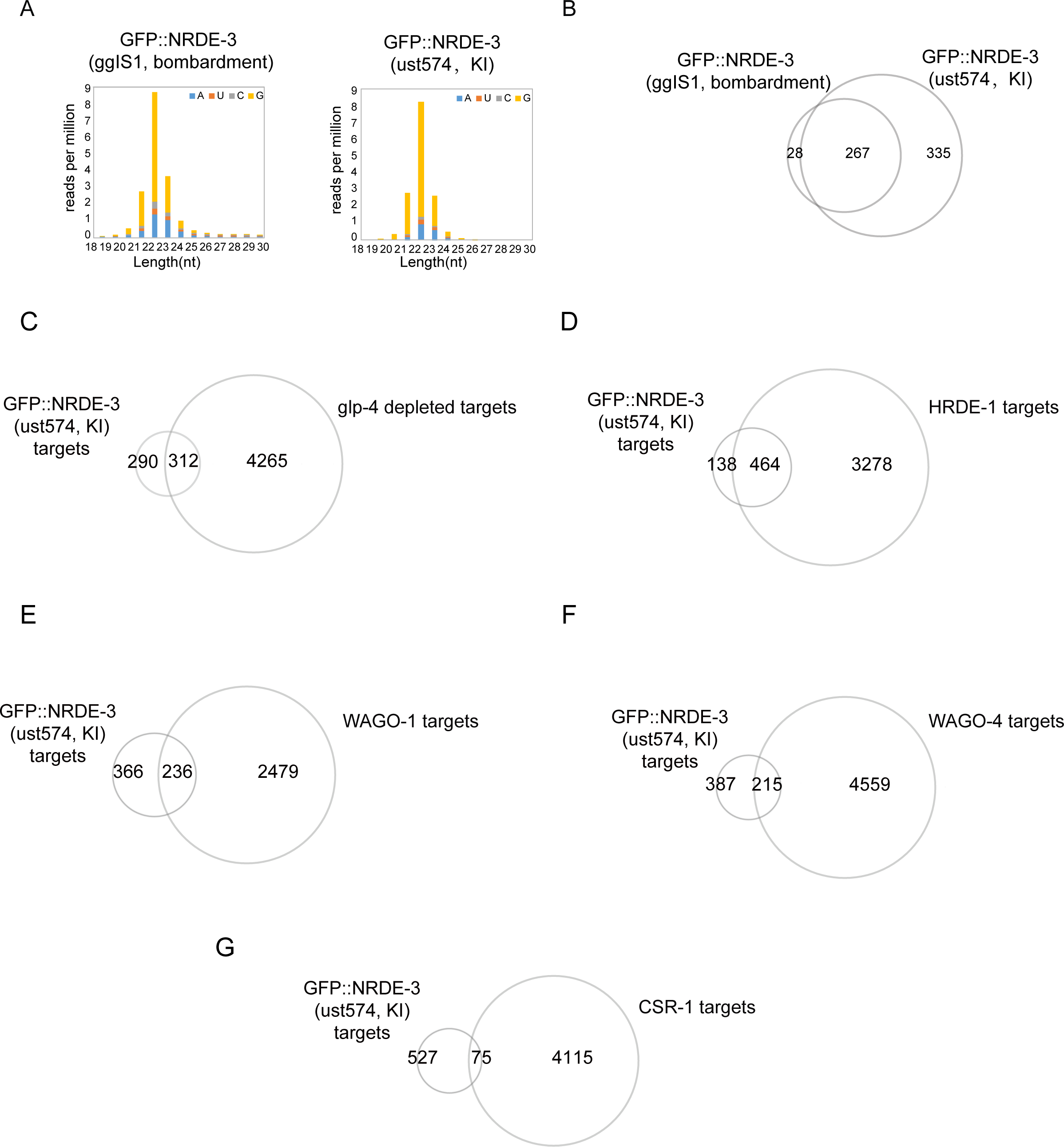
NRDE-3 binds 22G RNAs. (A) Length and first letter distribution of NRDE-3-associated siRNAs in embryos from the indicated animals. (B) Venn diagram of the overlapping gene targets of GFP::NRDE-3(ggIS1, bombardment) and GFP::NRDE-3(ust574, KI) in embryos. Cutoff > 25 reads per million. (C-G) Venn diagram showing the overlap between GFP::NRDE-3(ust574, KI) targets and other known siRNA categories.

**Figure S3.**
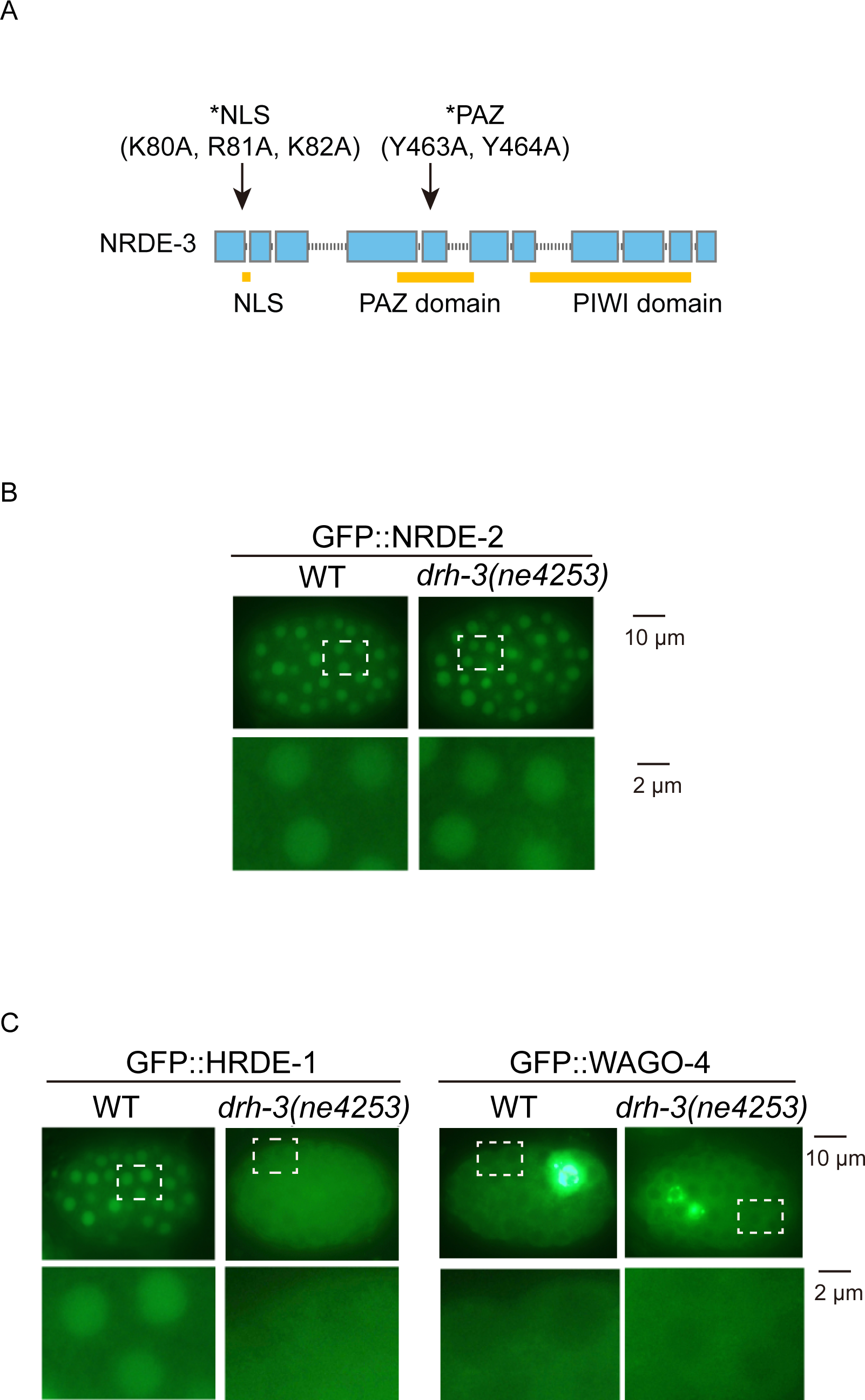
siRNA binding was required for NRDE-3 accumulation in the perinuclear foci. (A) Schematic of NRDE-3 and its variants. NLS, nuclear localization signal. (B, C) Fluorescence micrographs of the indicated embryos.

**Figure S4.**
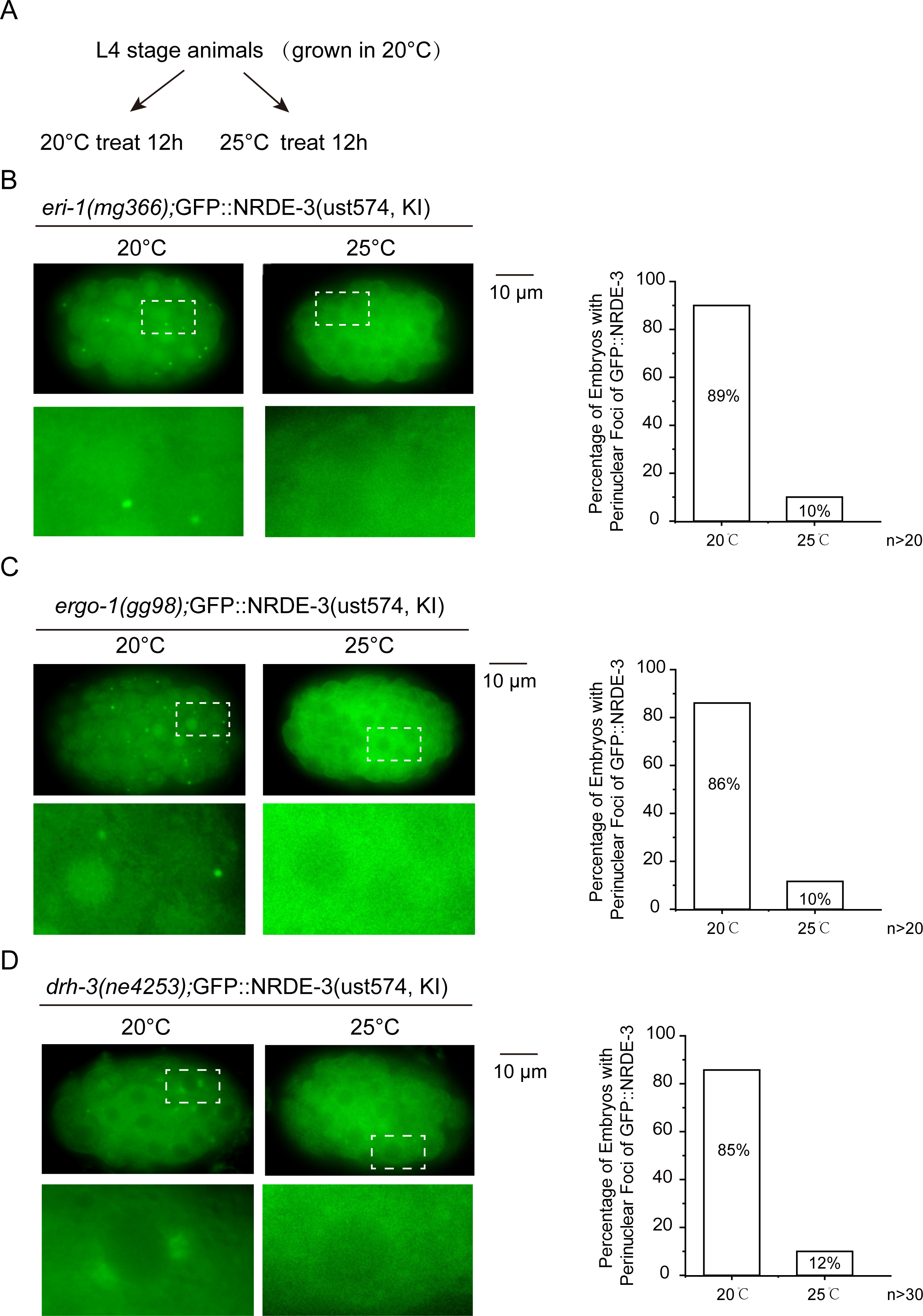
Culturing animals at 25°C depleted the perinuclear accumulation of NRDE-3. (**A**) Workflow of the temperature shift assay. Briefly, L4 stage animals grown at 20°C were divided onto new NGM plates and grown at 20°C or 25°C. Images were then taken 12 hours later, and the NRDE-3 foci were scored. (B, C, D) Left: Fluorescence micrographs of GFP::NRDE-3 embryos from the indicated animals. Right: The bar graph shows the percentages of NRDE-3 foci-positive embryos. An embryo with one or more NRDE-3 foci was defined as positive. At least 20 embryos were imaged for each condition.

**Figure S5.**
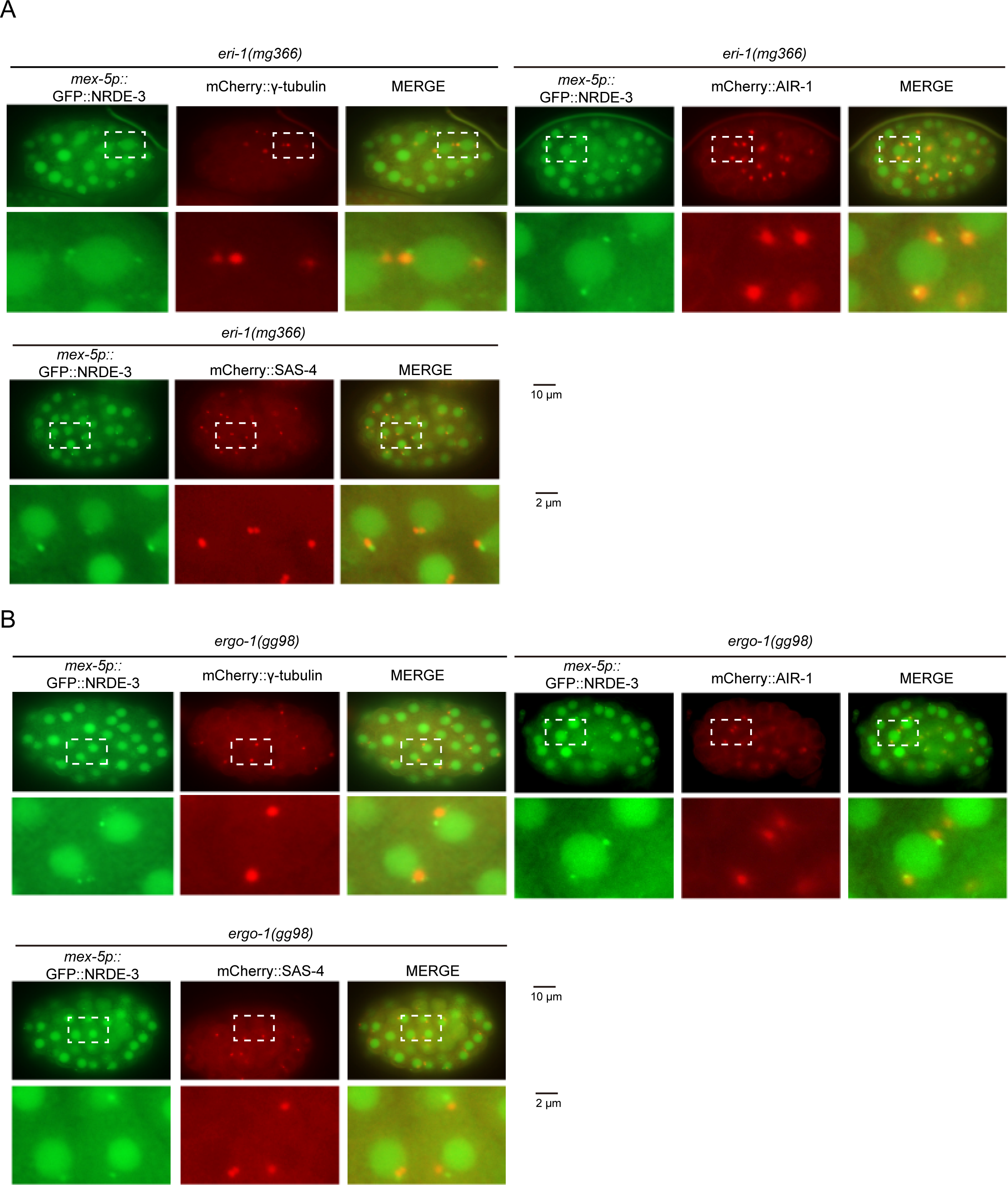
NRDE-3 accumulates in the peri-centrosomal foci. (A, B) Images of *mex-5p::*GFP::NRDE-3 with the indicated mCherry-tagged centrosome proteins in the *eri-1* (A) *and ergo-1* (B) backgrounds.

**Figure S6.**
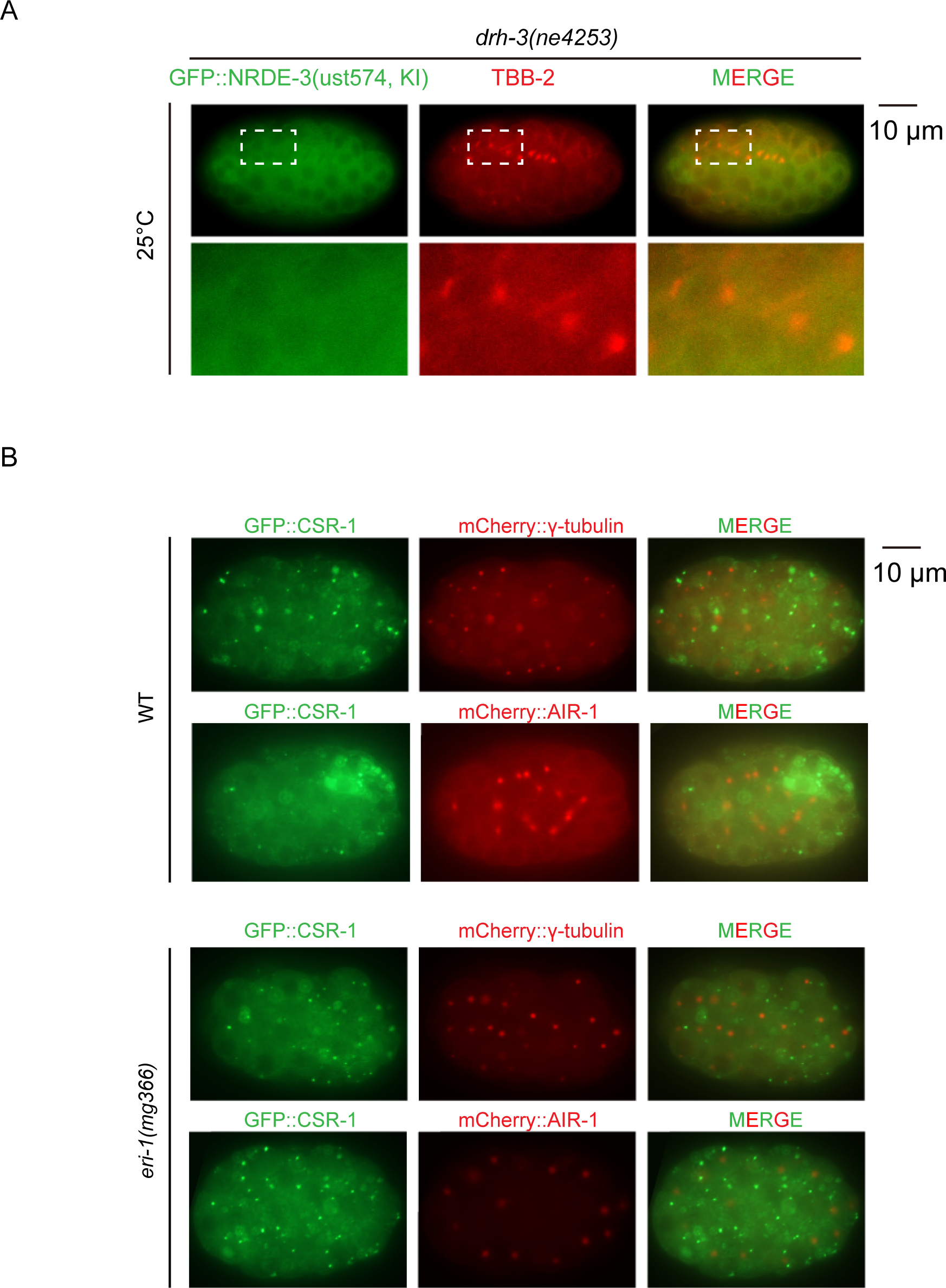
Culturing animals at 25°C depleted the peri-centrosomal accumulation of NRDE-3. (A) Indicated animals were grown at 20°C and shifted to 25°C at the L4 stage. Images were taken after 12 hours of treatment. (B) Images of the indicated GFP::CSR-1 embryos.

**Figure S7.**
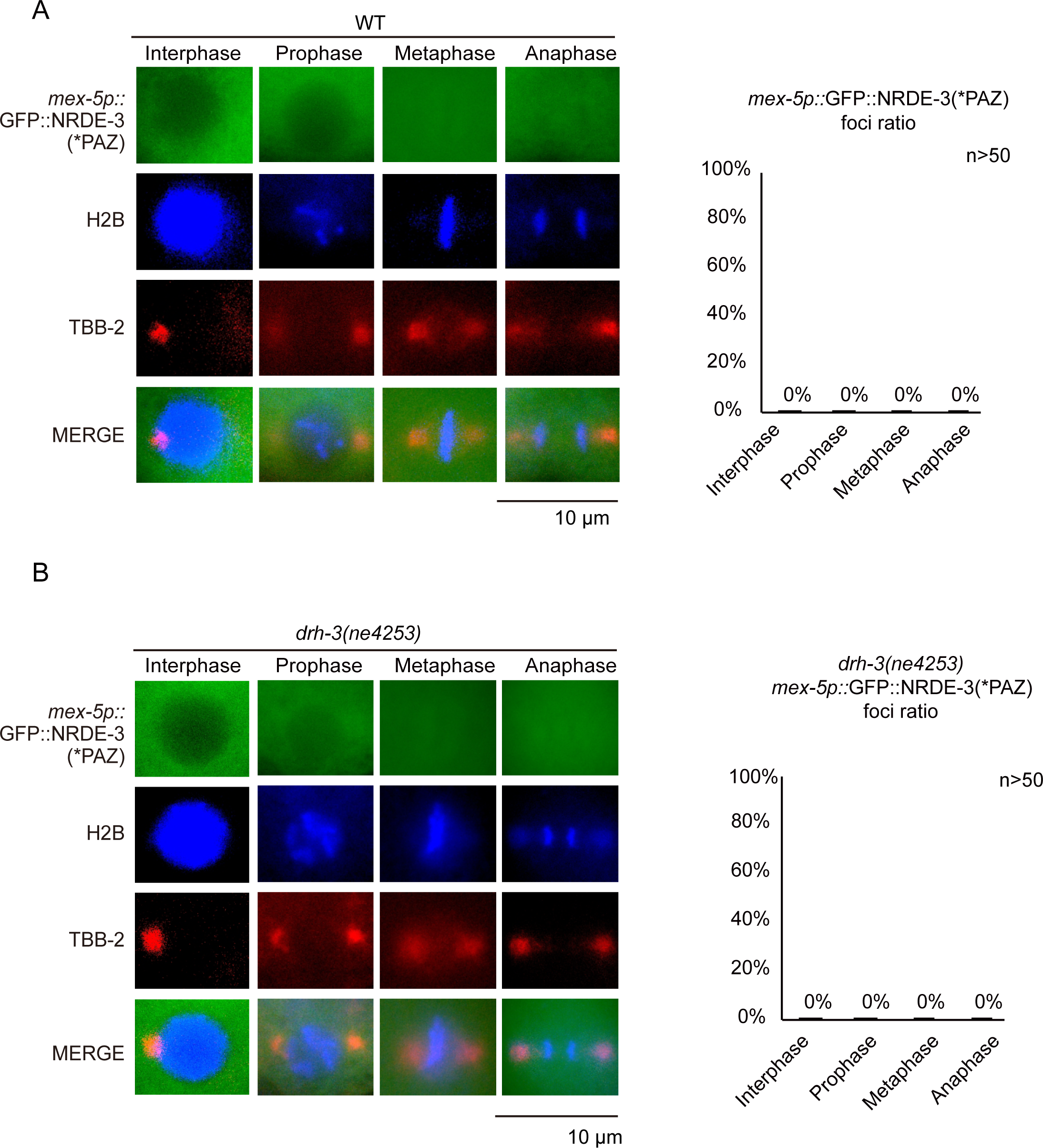
siRNA binding is essential for NRDE-3 accumulation in the peri-centrosomal foci and spindles during the cell cycle. (A, B) Left: Fluorescence microscopy images of interphase and mitosis in the indicated animals. Right: Bar graph depicting the percentage of peri-centrosomal localization of NRDE-3(*PAZ) foci in each phase. Embryos at the 10-30-cell stage were selected for quantification. For each embryo, each cell was assigned to different mitosis phases using BFP::H2B as a marker. We defined one or more NRDE-3(*PAZ) foci in one cell as positive and then counted the percentage of positive cells in each phase. At least 50 cells were quantified for each phase.

**Figure S8.**
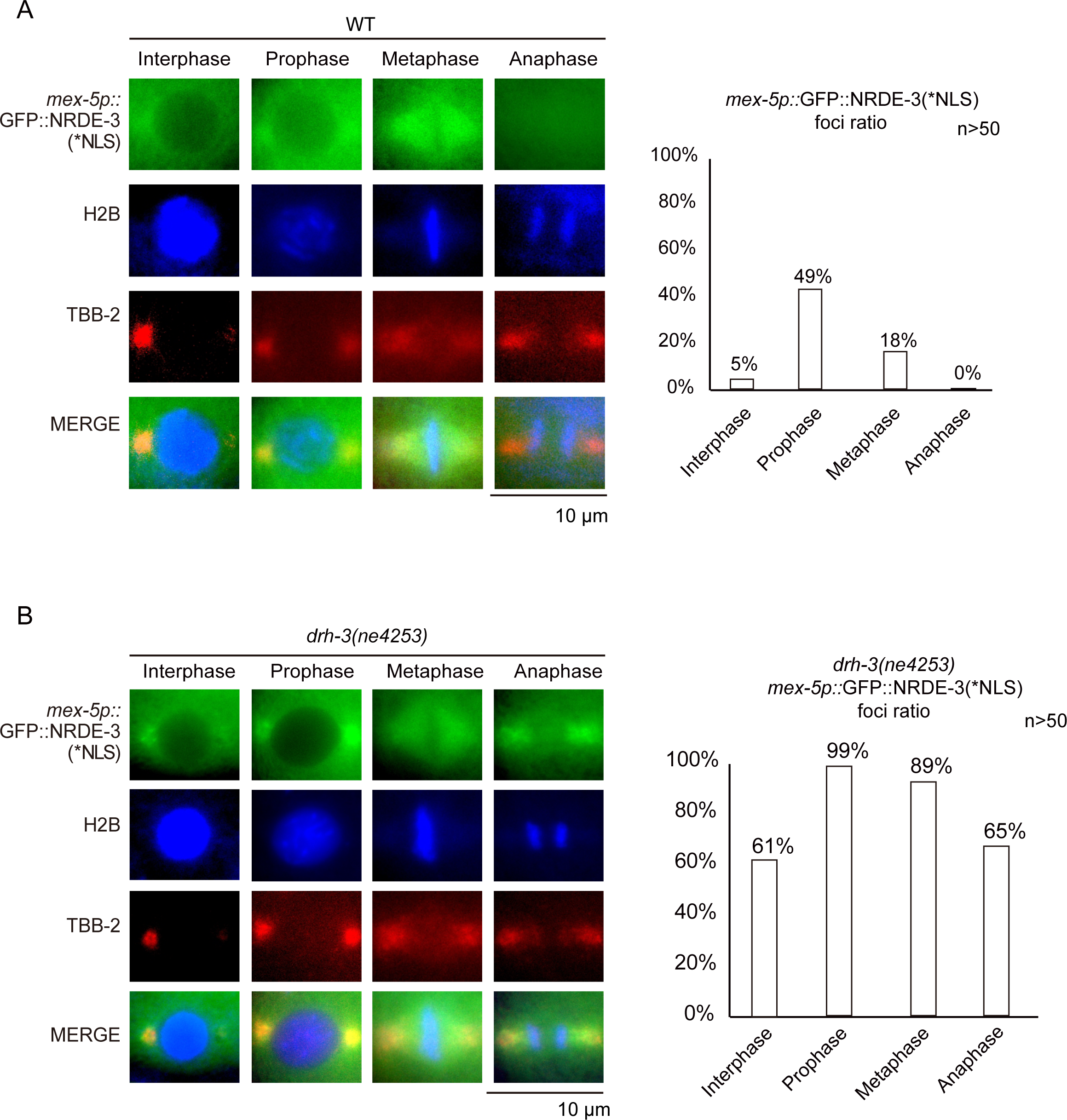
NRDE-3 (*NLS) accumulates in the peri-centrosomal foci and spindles during the cell cycle. (A, B) Left: Fluorescence microscopy images of interphase and mitosis in the indicated animals. Right: Bar graph depicting the percentage of peri-centrosomal localization of NRDE-3(*NLS) foci in each phase. Embryos at the 10-30-cell stage were selected for quantification. For each embryo, each cell was assigned to different mitosis phases using BFP::H2B as a marker. We defined one or more NRDE-3(*NLS) foci in one cell as positive and then counted the percentage of positive cells in each phase. At least 50 cells were quantified for each phase.

**Figure S9.**
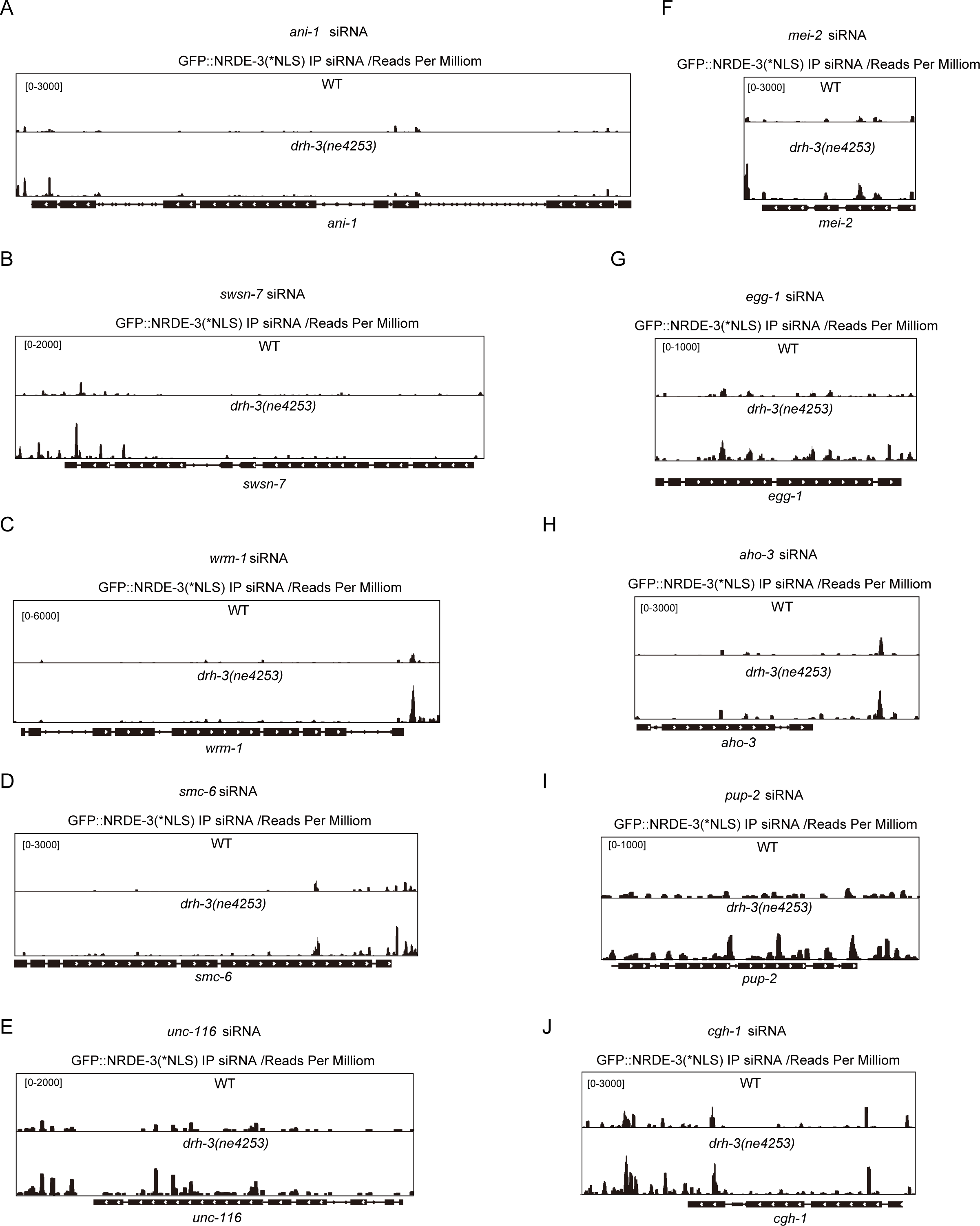
The peri-centrosomal-enriched siRNAs predominantly mapped to the 3’ portion of the target genes. (A-J) The distribution of normalized NRDE-3(*NLS)-associated small RNA reads across indicated genomic loci in wild type and *drh-3(ne4253)* animals.

**Figure S10.**
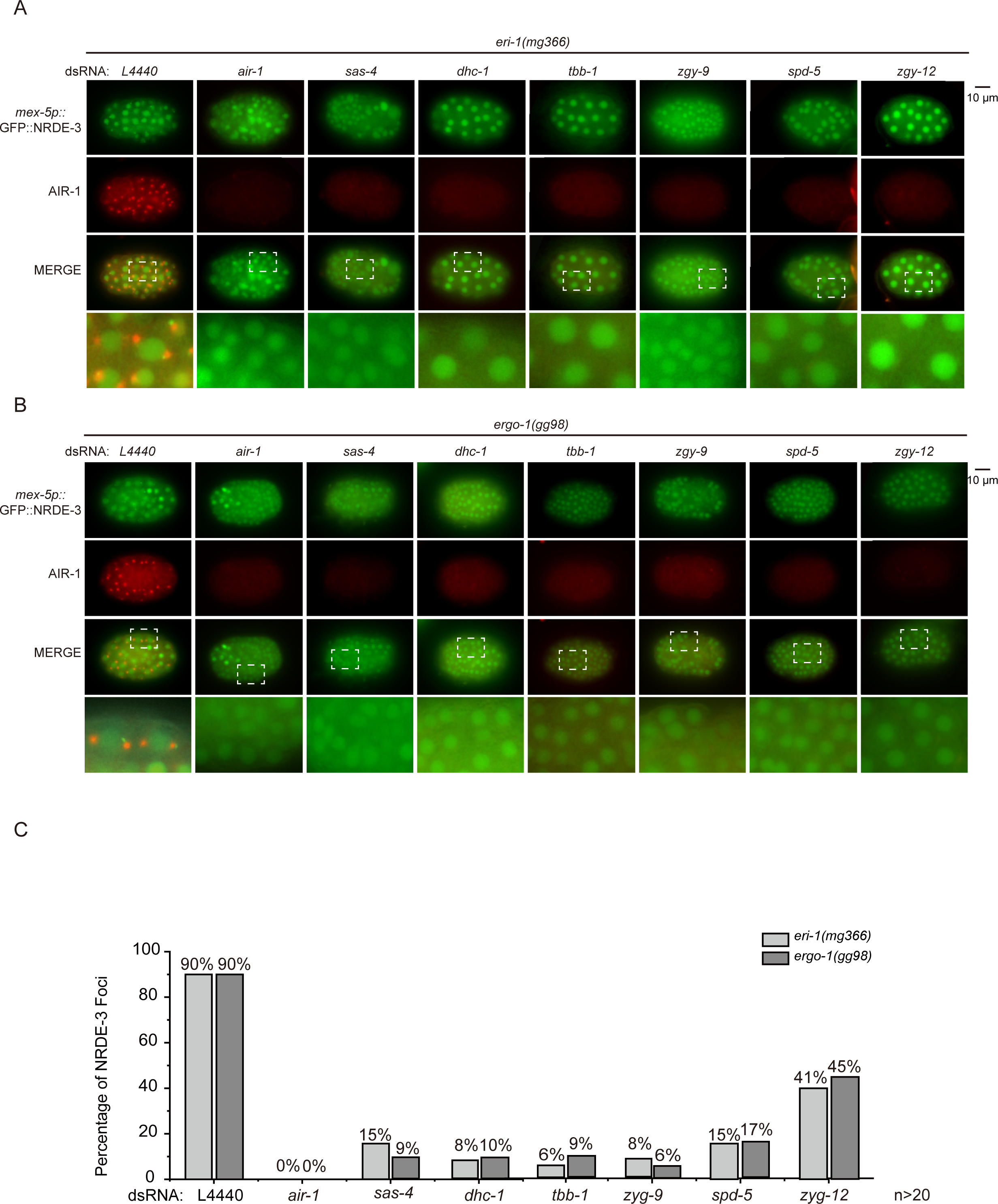
The integrity of the centrosome is required for the peri-centrosomal accumulation of NRDE-3. (A, B) Images of *eri-1*(*-*);GFP::NRDE-3;mCherry::AIR-1 (A) and *ergo-1*(*-*);GFP::NRDE-3;mCherry::AIR-1 (B) animals after RNAi targeting of the centriole, and PCM genes. Synchronized embryos were hatched and cultured on NGM plates for 41 hours and then transferred to RNAi plates at the L4 stage. F1 embryos were imaged. (C) Bar graph quantifying the percentage of NRDE-3 foci-positive embryos. The embryos with one or more NRDE-3 foci were considered positive. At least 20 embryos were imaged for each RNAi experiment.

**Table S1.**
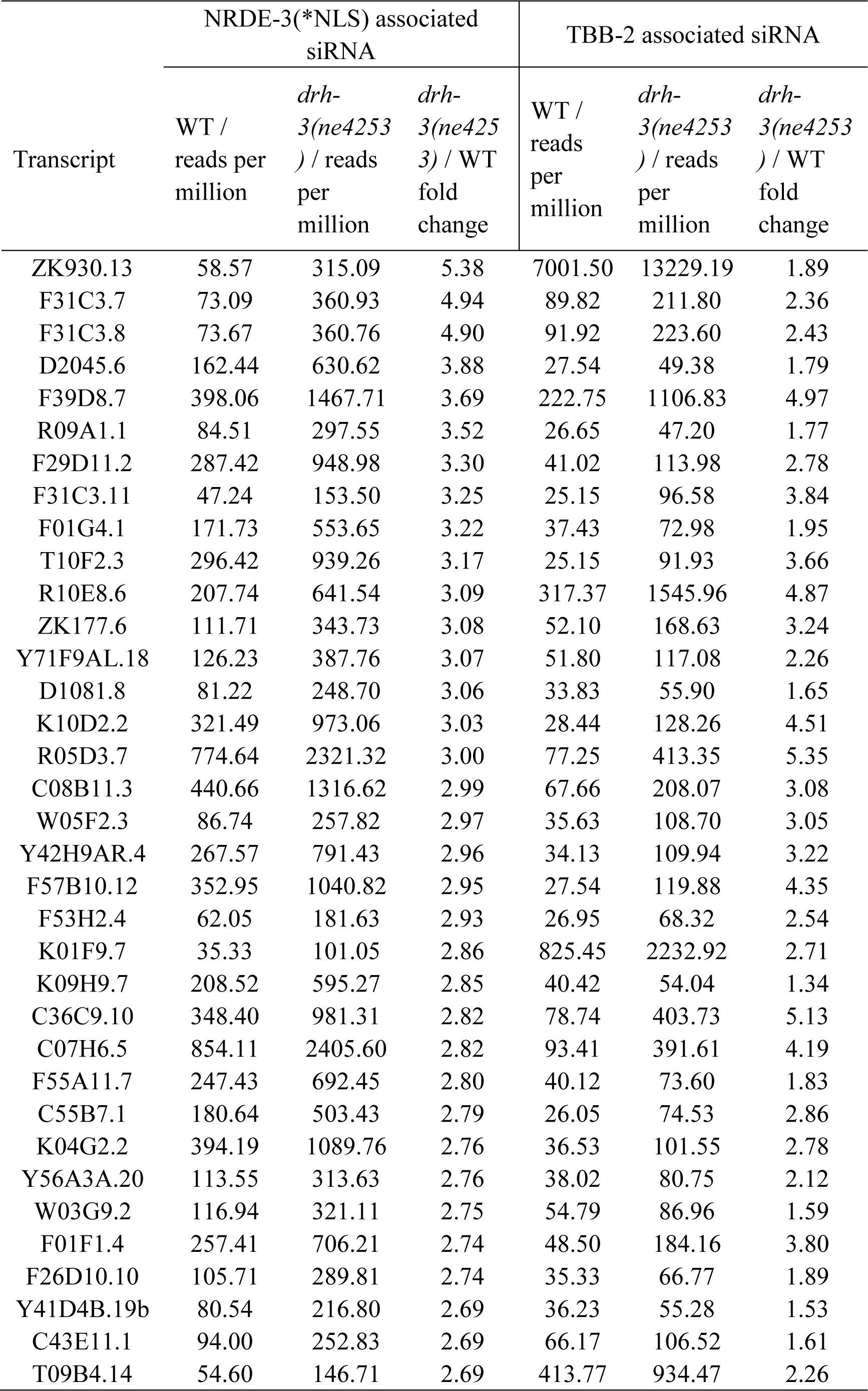
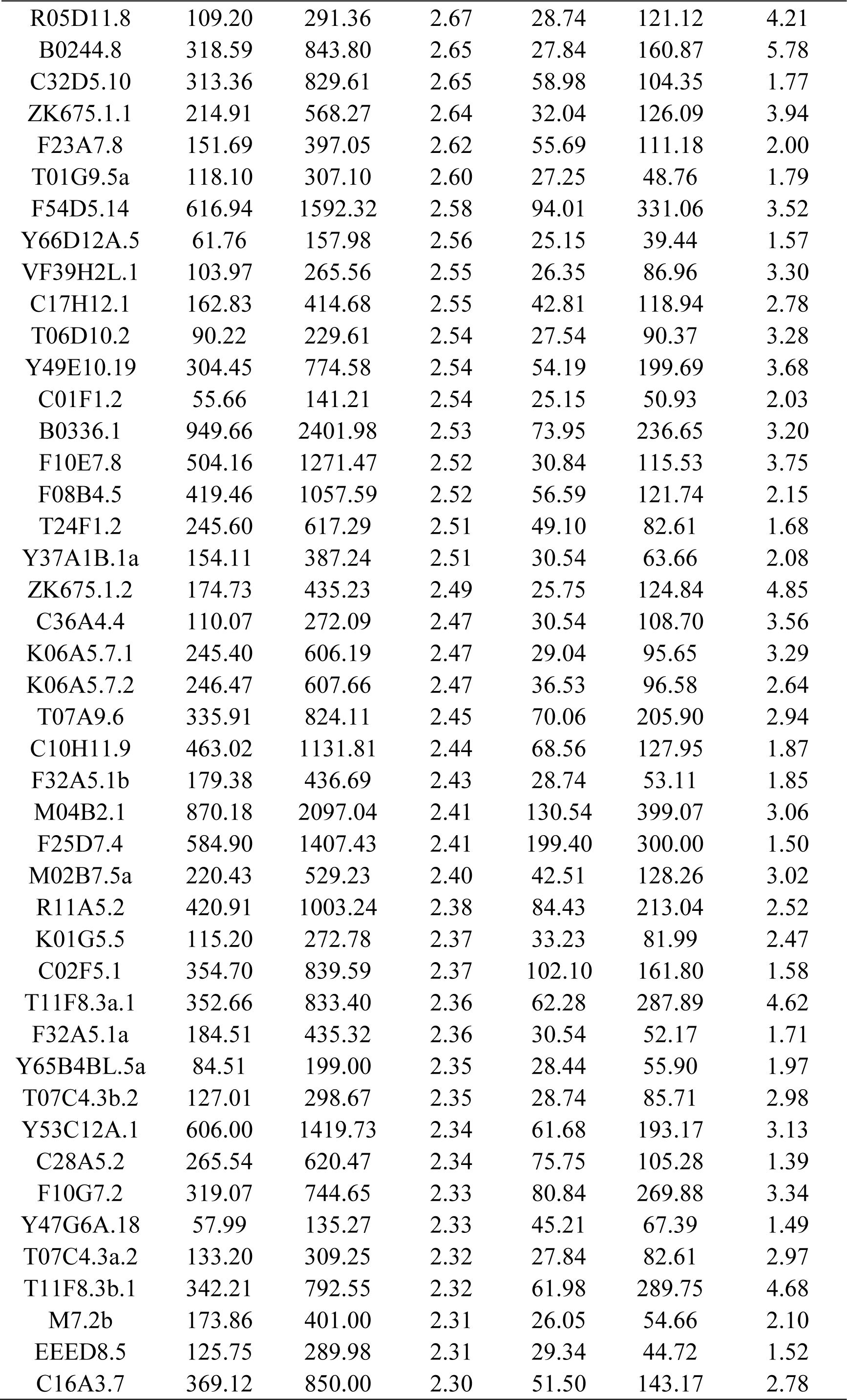
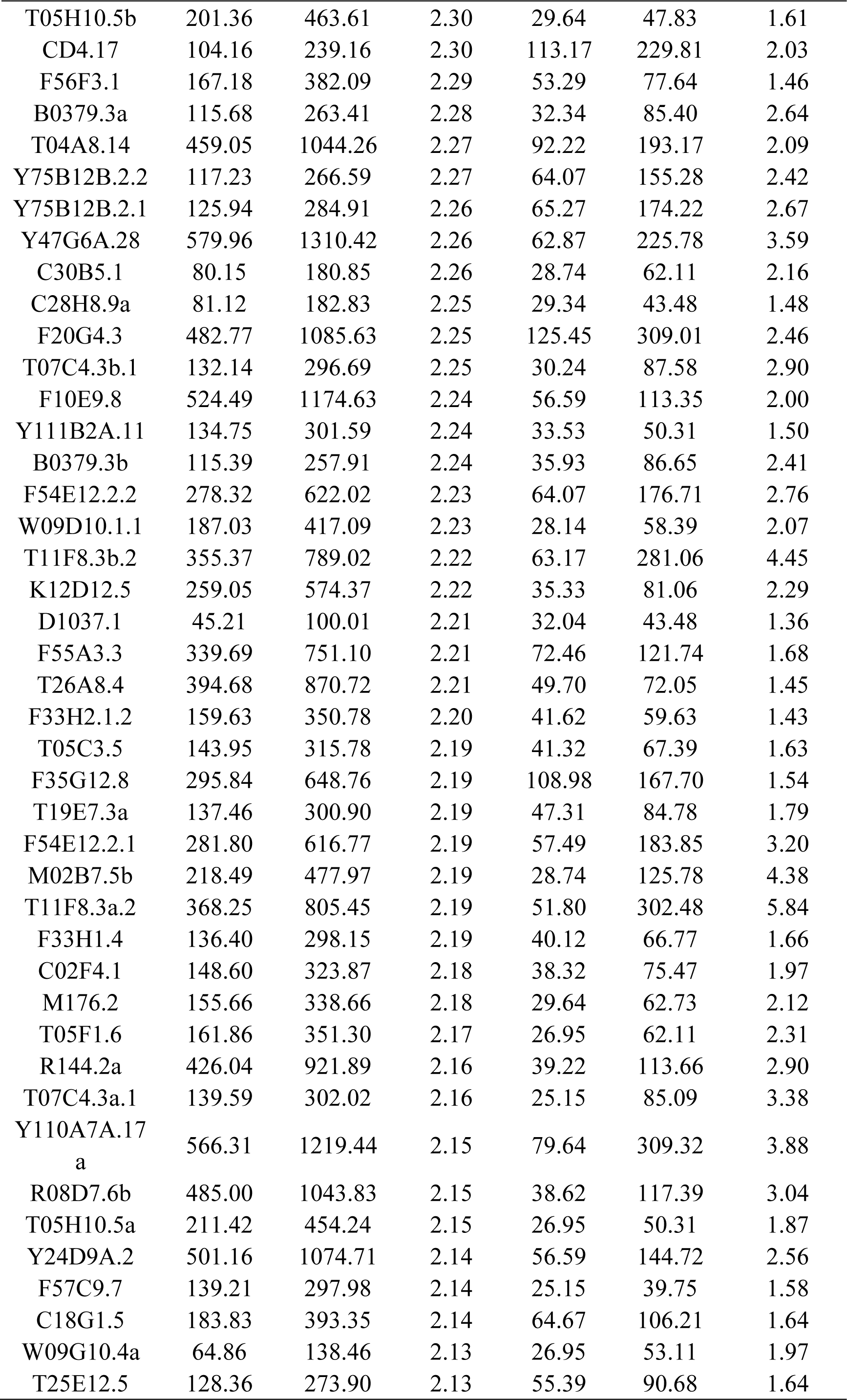
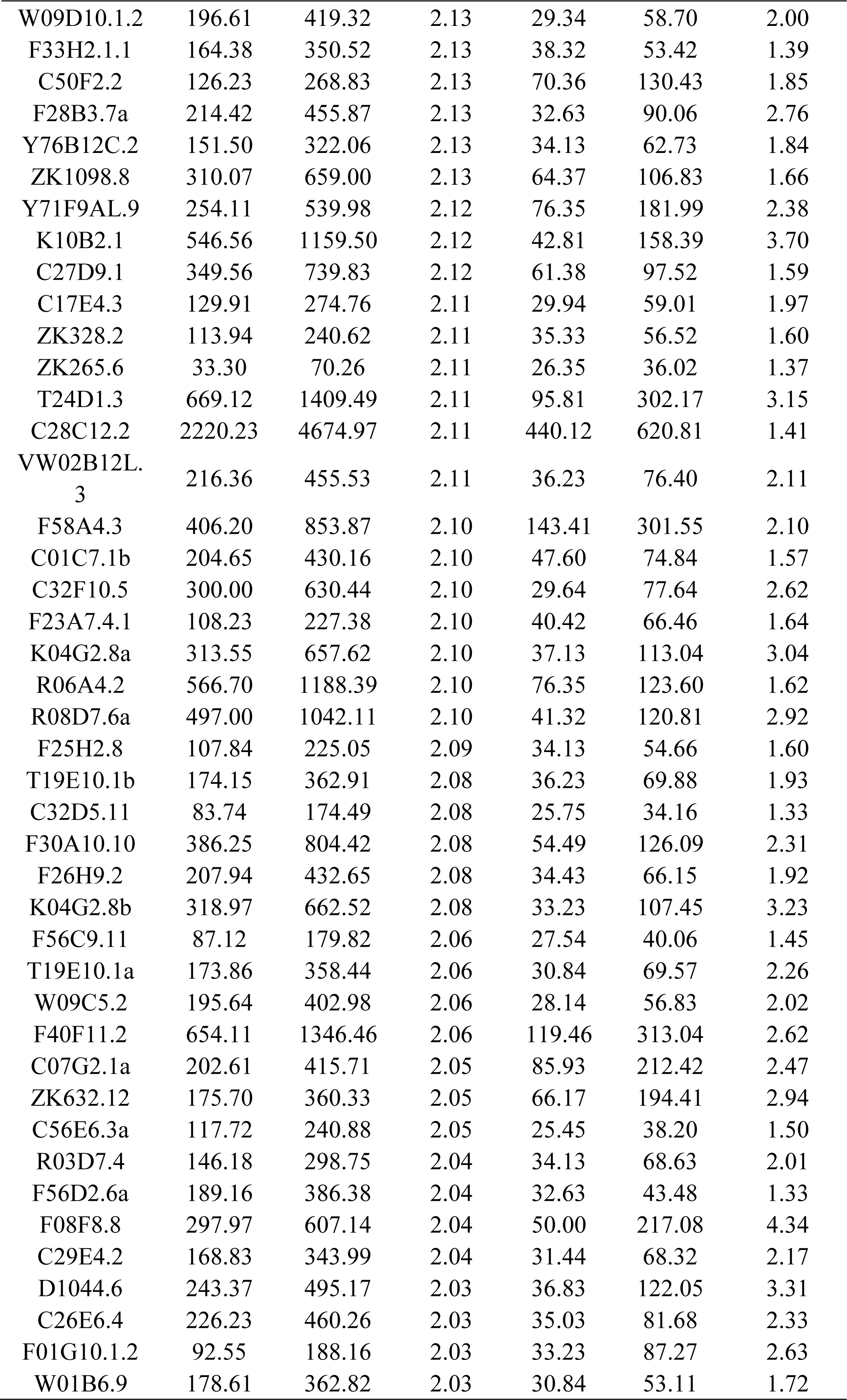
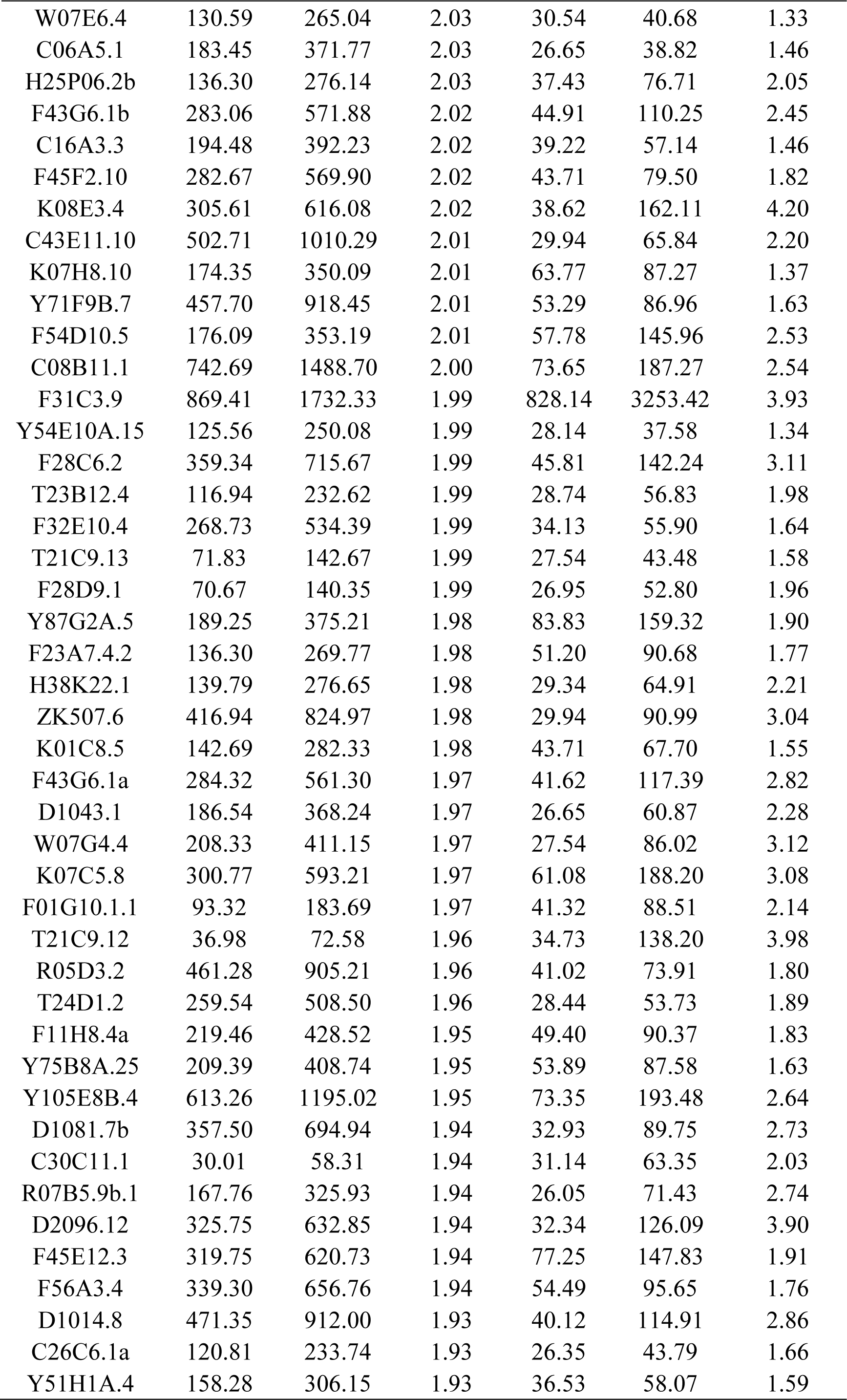
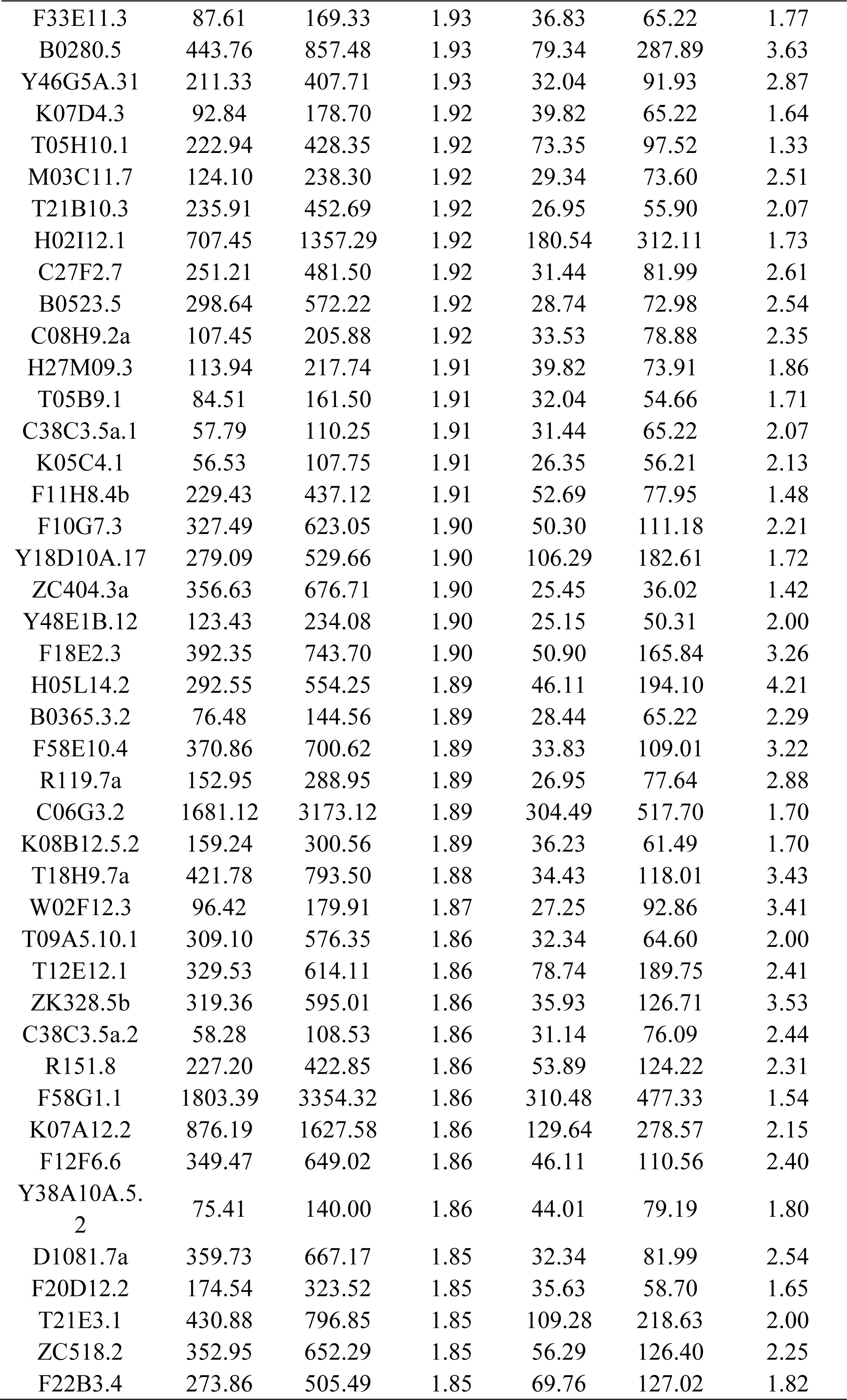
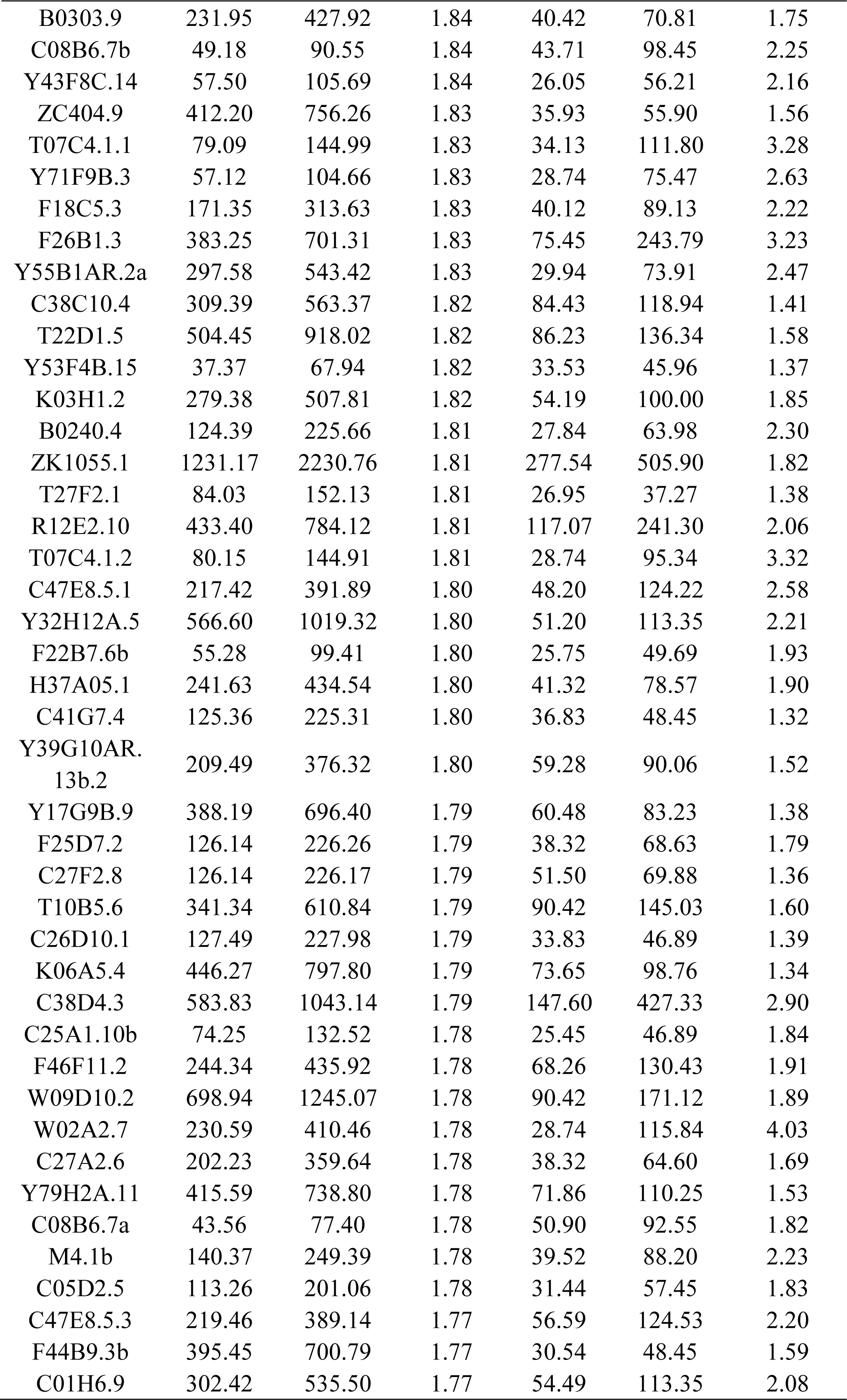
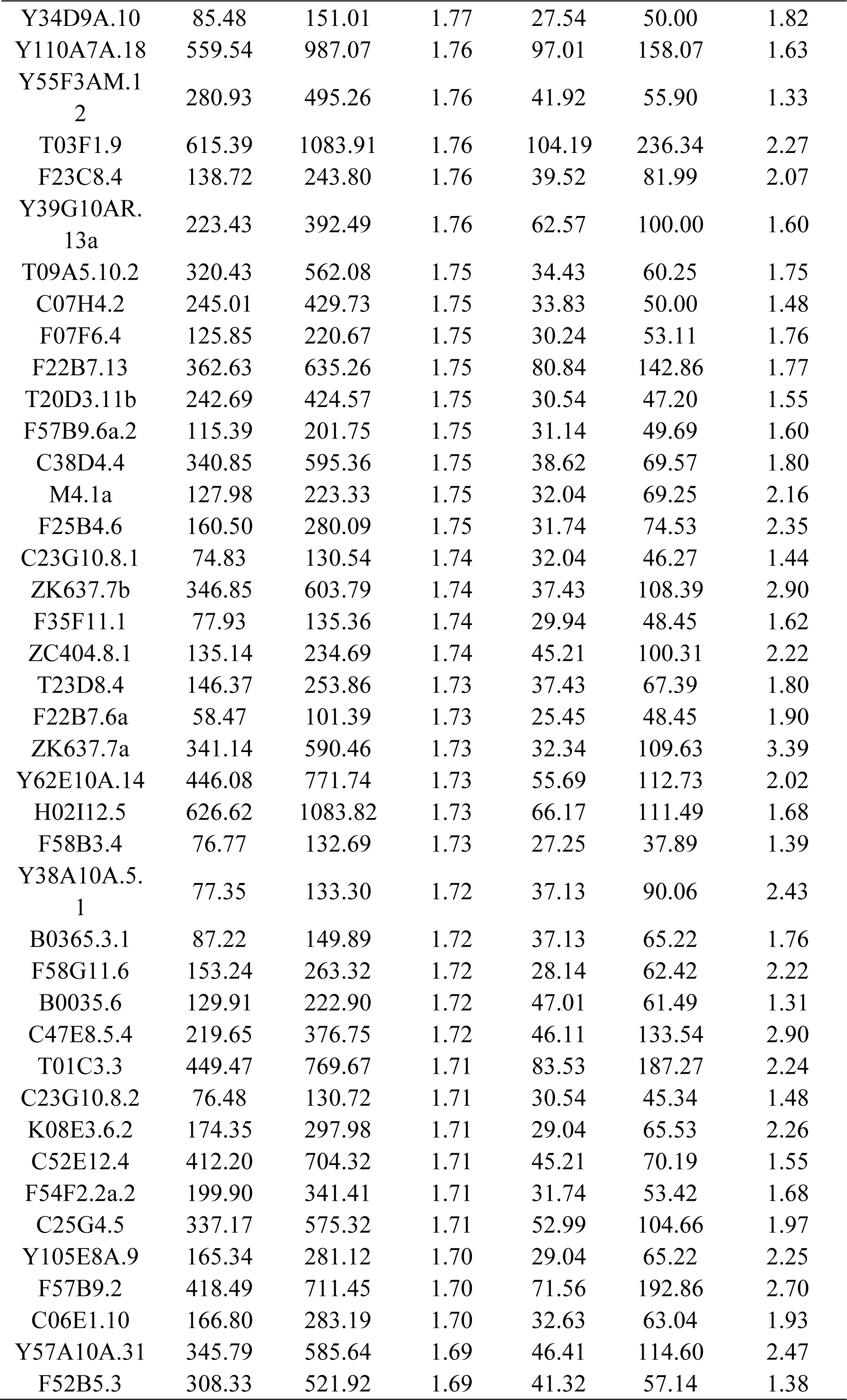
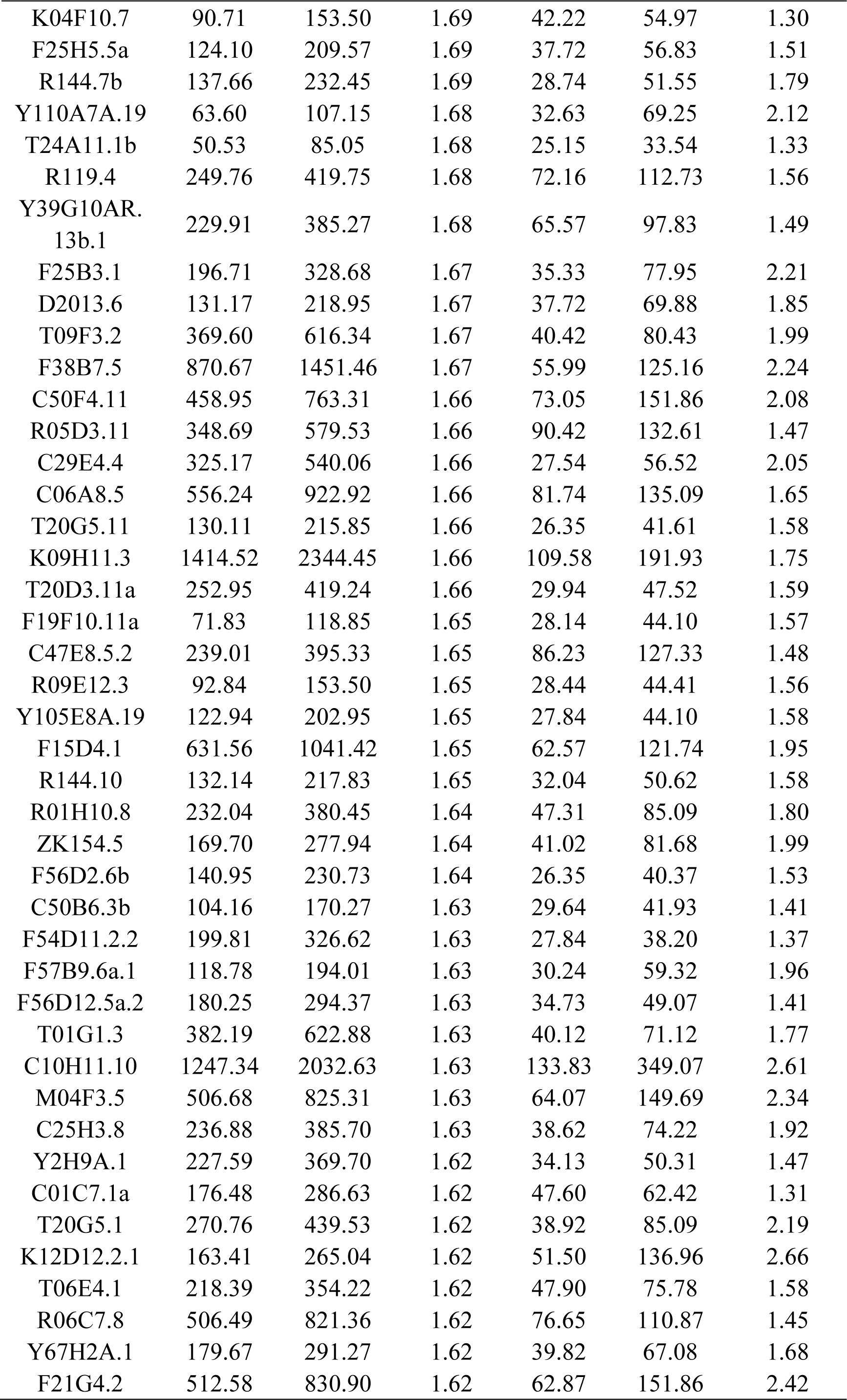
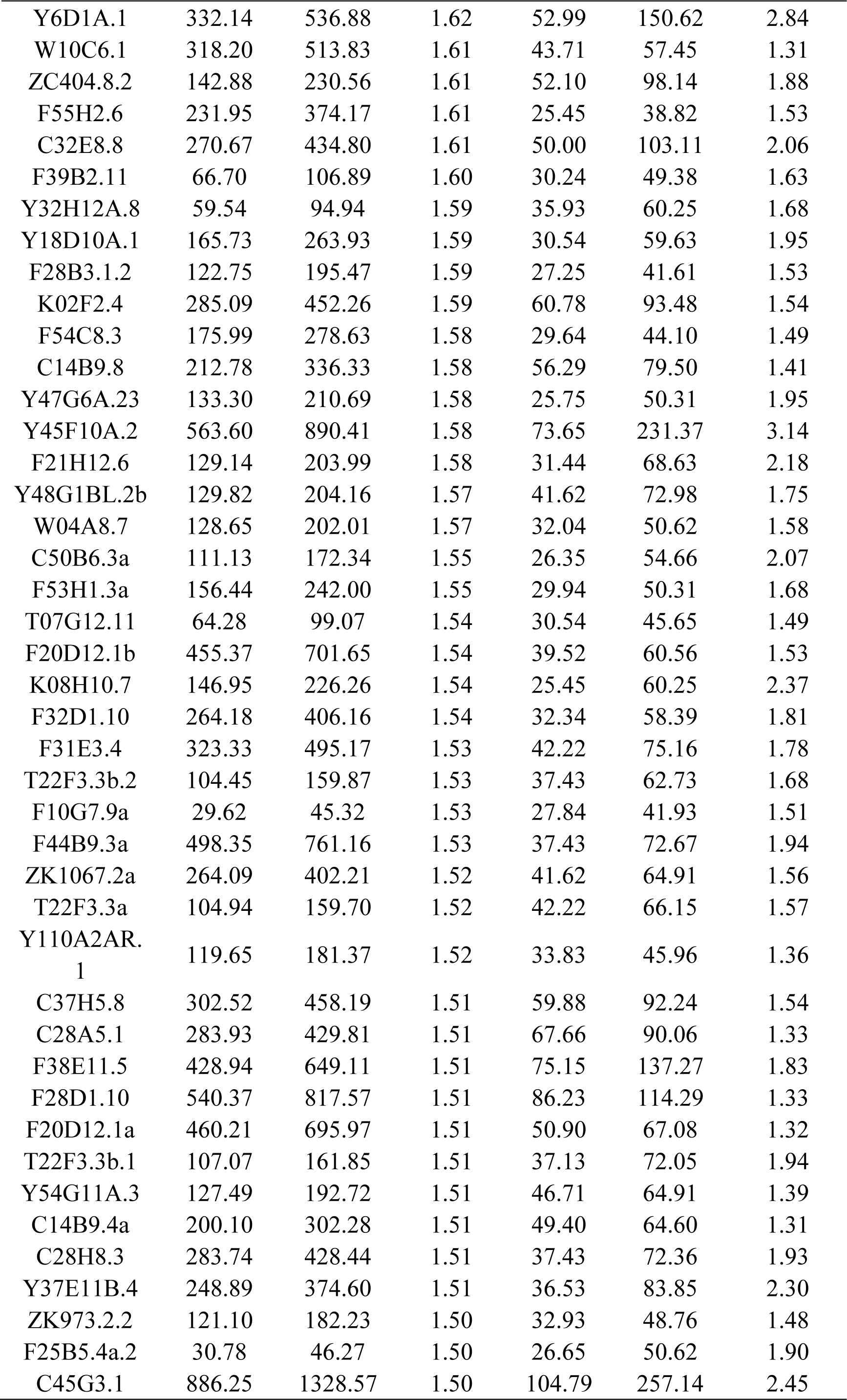
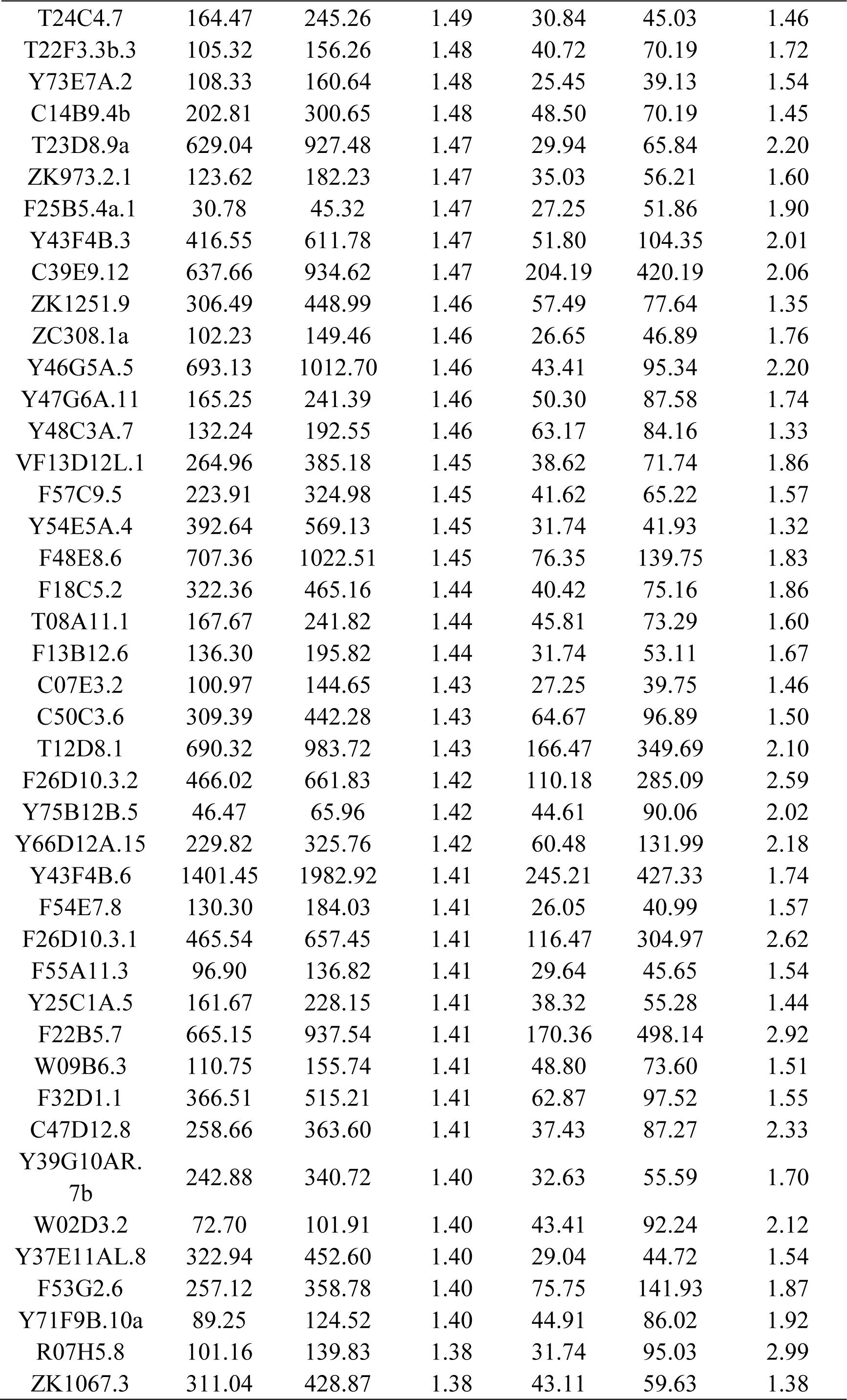
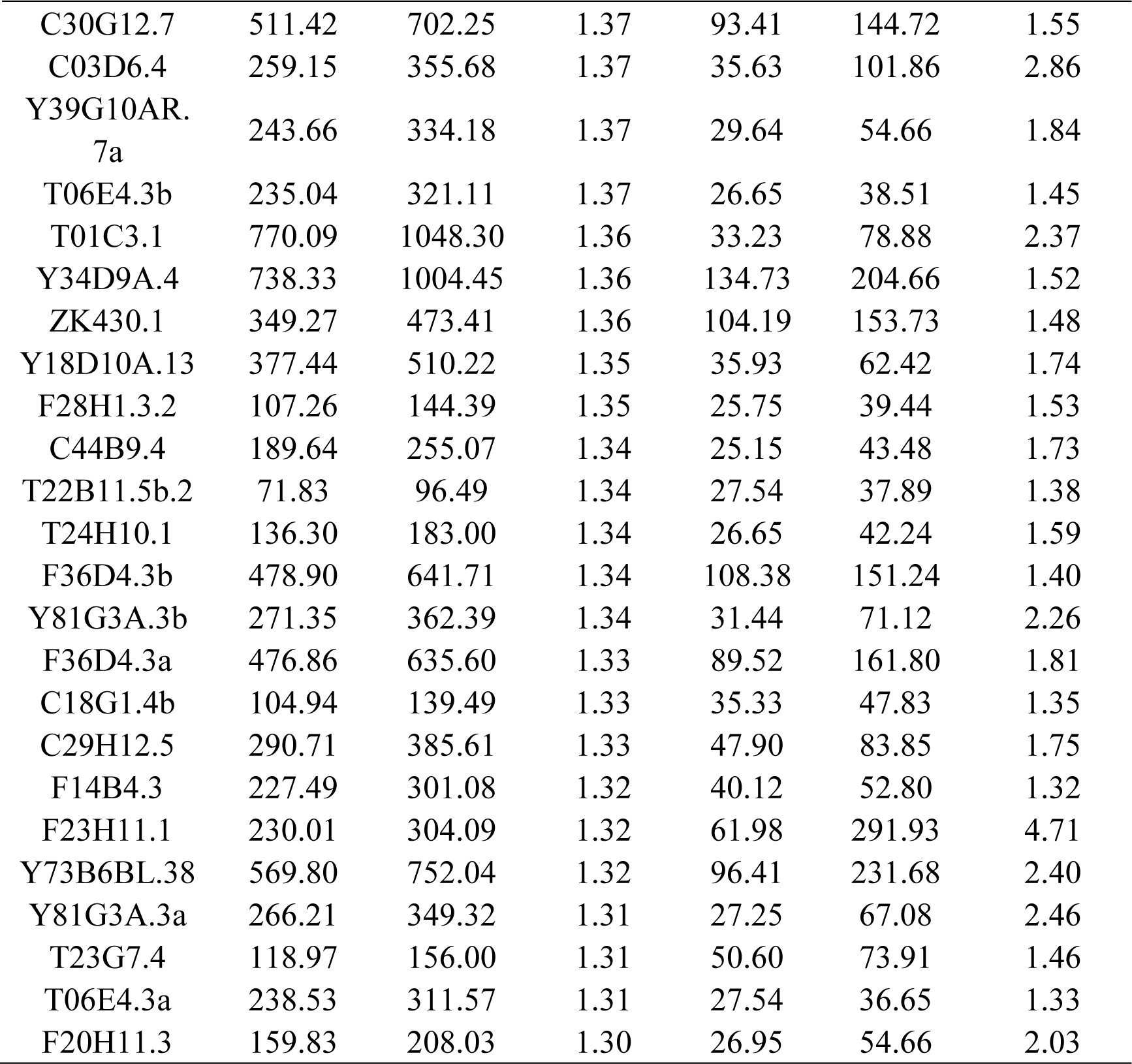
Peri-centrosomal-enriched siRNA targets.

**Table S2.**
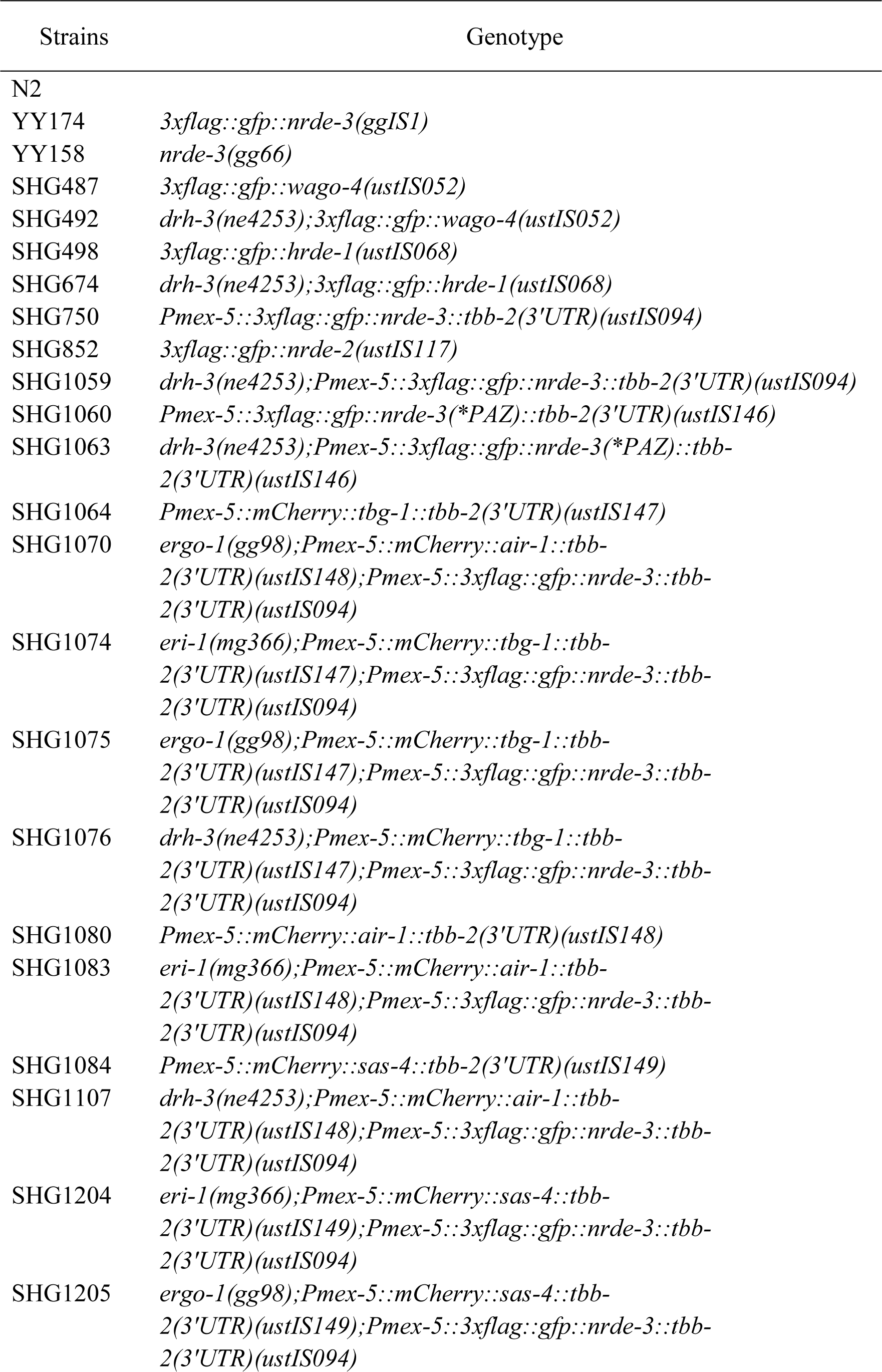
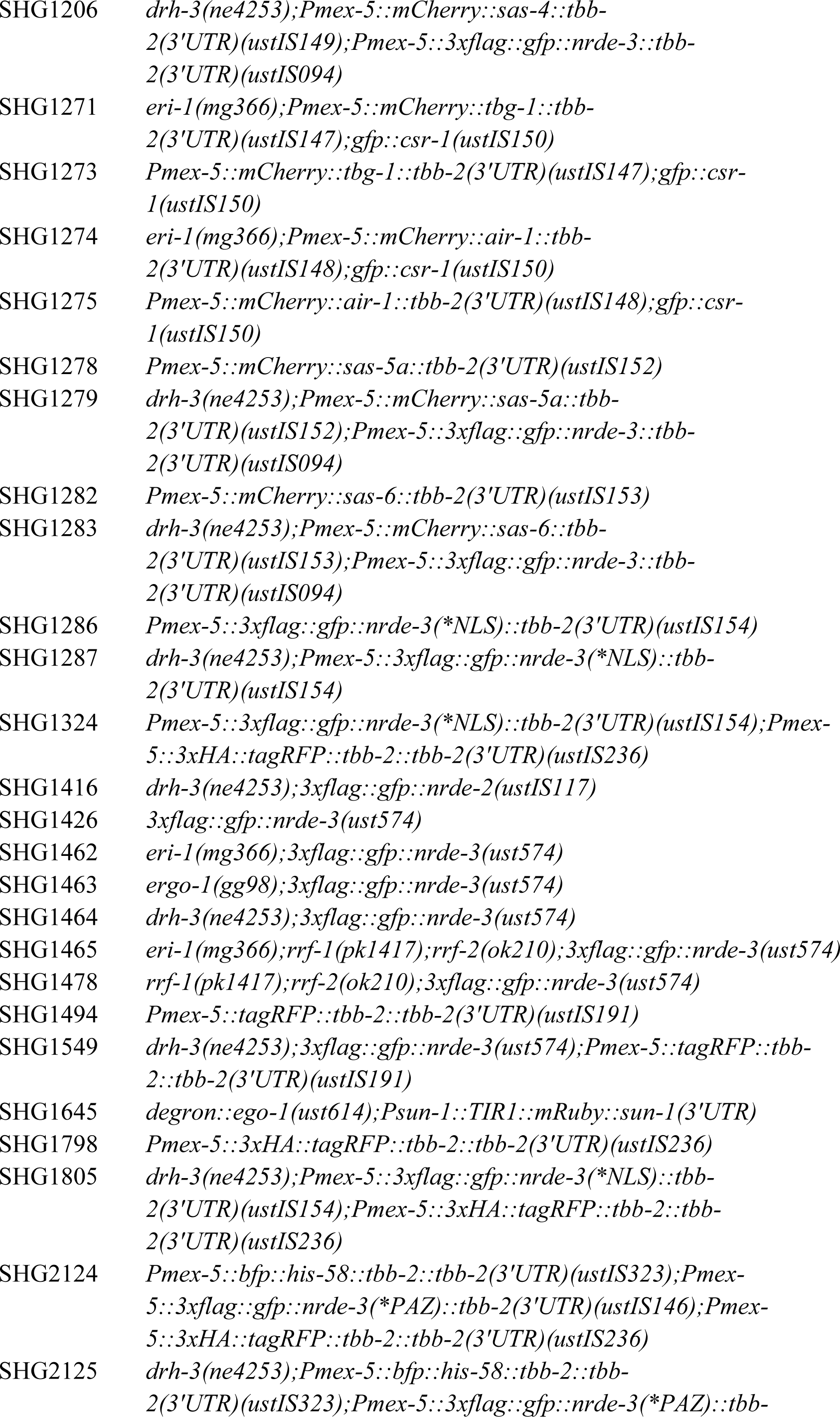
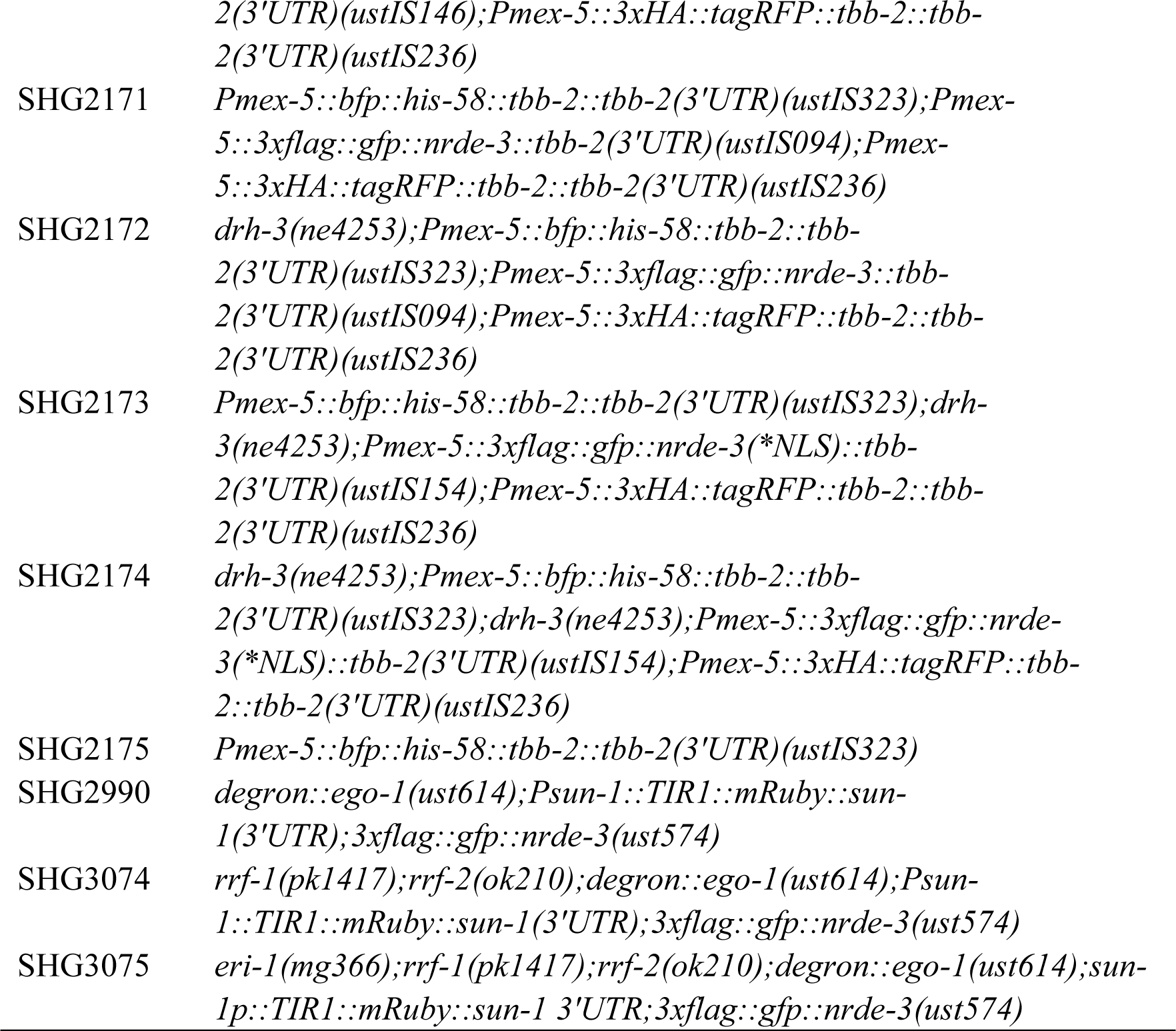
List of strains used in this study.

**Table S3.**
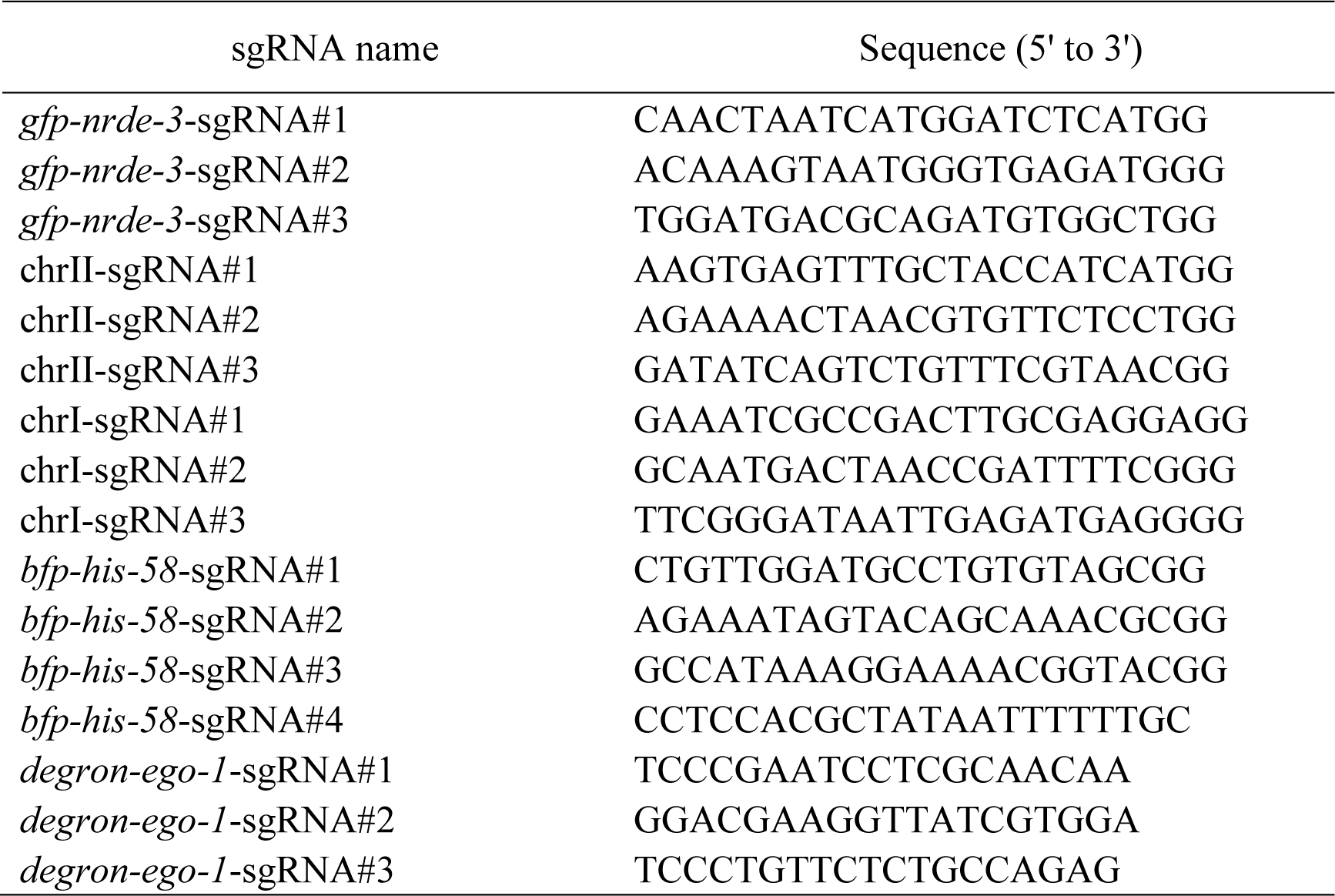
Sequence of sgRNAs for CRISPR/Cas9-mediated gene editing.

**Table S4.**
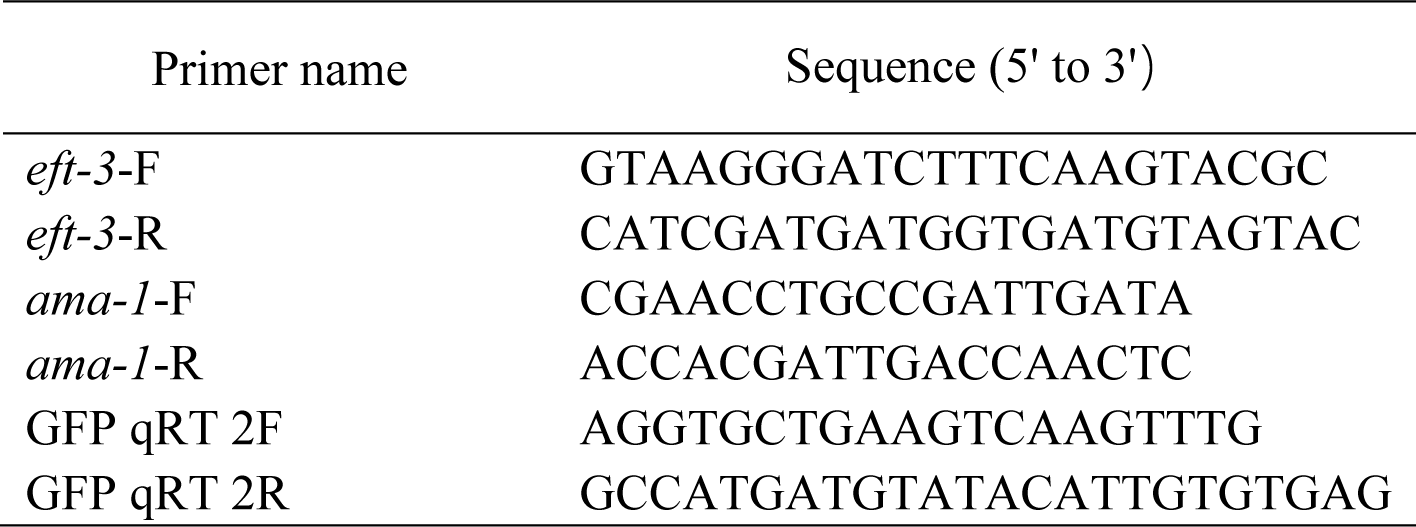
Sequences of the quantitative real-time PCR primers.

