## Supporting Information - SI Appendix for "Peri-centrosomal localization of small interfering RNAs in *C. elegans*"

**Supplemental materials and methods**

**Construction of plasmids**

All plasmids were generated through the recombinational cloning of PCR-amplified fragments. A ClonExpress MultiS One Step Cloning Kit (Vazyme) was used to connect these fragments to the vector.

For the *in situ* knock-in transgene 3xFLAG::GFP::NRDE-3(ust574, KI), the 3xFLAG::GFP sequence was PCR amplified with the primers 5'-ATGGACTACAAAGACCATGACGGT-3' and 5'-AGCTCCACCTCCACCTCC-3' from YY178 genomic DNA. The homologous left arm (1.2 kb) was PCR amplified with the primers 5'- ATAACAATTTCACAGGGCCCCCGTCGATCAAGTTTGCCGG -3' and 5'- ATAACAATTTCACAGGGCCCCCGTCGATCAAGTTTGCCGG -3' from N2 genomic DNA. The homologous right arms (1.3 kb) were PCR amplified from N2 genomic DNA with the primers AAGGAGGTGGAGGTGGAGCTATGGATCTCCTAGACAAAGTAATG-3' and CCAGTCACGACGTCACGTGAATCAGAGTAACCTCGTCGGG-3'. The backbone was PCR amplified with the primers 5'- CACGTGACGTCGTGACTGGG -3' and 5'-GGGCCCTGTGAAATTGTTATCC -3' from the plasmid pCFJ151.

For the *in situ* knock-in *degron::ego-1(ust614)* transgene, the homologous left arm (1.4 kb) was PCR amplified with the primers 5'-ATAACAATTTCACAGGGCCCCGTCCAAATCTTGGTTTCTGGC-3' and 5'-CGGTATGATCTCACCGGTGGCCATCCCACAACTTGTGCCTTGGCCGGAGGTTTGGCTGGATCTTTAGGCATTGTTGCGAGGATTCGGGATA-3' from N2 genomic DNA. The homologous right arms (1.5 kb) were PCR amplified with the primers 5'-CCACCGGTGAGATCATACCGGAAGAACGTGATGGTTTCCTGCCAAAAATCAAGCGGTGGCCCGGAGGCGGCGGCGTTCGTGAAGGGAGGTGGAGGTGGAGCTATGGGGGACGAAGGTTATCG- 3' and 5'-

CCCAGTCACGACGTCACGTGGATCCTTAACCACTGCTCCTCTC -3’ from N2 genomic DNA. The degron sequence was added to the abovementioned PCR primers. The backbone was PCR amplified with the primers 5’- CACGTGACGTCGTGACTGGG -3’ and 5’- GGGCCCTGTGAAATTGTTATCC -3’ from the plasmid pCFJ151.

For the ectopic transgene *mex-5p*:: 3xFLAG::GFP::NRDE-3(*PAZ):tbb-2 UTR inserted in chr II, the *mex-5p*::3xFLAG::GFP::NRDE-3(*PAZ)::tbb-2 UTR was amplified with the primers 5’- CTTCCATCACAGAGGCTGCCTTACAACGGTACAACTATCGTCTC -3’ and 5’- GGCAGCCTCTGTGATGGAAGTGAGCTTC -3’ from shg750 genomic DNA. The vector fragment was the same as that used for the chr II vector above.

For the ectopic transgene *mex-5p*::GFP::NRDE-3(m)(*NLS):tbb-2 UTR inserted in chr II, the *mex5-p*:: 3xFLAG::GFP::NRDE-3(*NLS)::tbb-2 UTR was amplified with the primers 5’- GCAGCAGCGCCACTGGGAGGATGTGGGT -3’ and 5’- CCTCCCAGTGGCGCTGCTGCTGGATCGGGACCTGTGCGAT -3’ from 750 genomic DNA. The vector fragment was the same as that used for the chr II vector above.

For the ectopic transgene *mex-5p*::tagRFP::TBB-2 inserted in chr I, the MEX-5 promoter was amplified with the primers 5’- CCTCCCTGTCAATTCCCAAAATACAAATATCAGTTTTTAAAAAATTAAACCAT -3’ and 5’- TCTTCTCCCTTGGACACCATTCTCTGTCTGAAACATTCAATTG -3’ from N2 genomic DNA. The tagRFP sequence was PCR amplified with the primers 5’-ATGGTGTCCAAGGGAGAAGA-3’ and 5’-GTTTGAGCTTGTGCCCGAGC-3’ from yy967 genomic DNA. The coding sequence (CDS) region and 3′ untranslated region (UTR) sequence were PCR amplified with the primers 5’- GCTCGGGCACAAGCTCAACGGAGGTGGAGGTGGAGCTATGAGAGAGATCGTCCACGT -3’ and 5’-

TTCAAAGAAATCGCCGACTTCCAACCACATATGTTTCTCTTAGGC -3’ from N2 genomic DNA. The chr I vector fragment was PCR amplified with the primers 5’-AAGTCGGCGATTTCTTTGAAGTT-3’ and 5’-GTATTTTGGGAATTGACAGGGAGG-3’ from the plasmid pSG274.

For *mex-5p*::3xHA::tagRFP::TBB-2 inserted in chr I, *mex-5p*::3xHA::tagRFP::TBB-2 was PCR amplified with the primers 5’-TCCATATGACGTGCCGGACTATGCATACCCATACGATGTTCCAGATTACGCTTACCCATACGATGTTCCAGATTACGCTATGGTGTCCAAGGGAGAAGA -3’ and 5’-AGTCCGGCACGTCATATGGATACATTCTCTGTCTGAAACATTCAATTG-3’ from shg1494 genomic DNA. The vector fragment used was the same as that used for the chrI vector above.

For *mex-5p*::Cherry::TBG-1::tbb-2 utr inserted in chr I, the MEX-5 promoter was amplified with the primers 5’-CCTCCCTGTCAATTCCCAAAATACAAATATCAGTTTTTAAAAAATTAAACCAT -3’ and 5’- TCCTCTCCCTTGGAGACCATTCTCTGTCTGAAACATTCAATTG -3’ from N2 genomic DNA. The mCherry sequence was PCR amplified with the primers 5’-ATGGTCTCCAAGGGAGAGGA-3’ and 5’-ATAGCTCCACCTCCACCTCCCTTGTAAAGCTCATCCATTCCTCC-3’ from the SX2650 genomic DNA. The coding sequence (CDS) region sequence was PCR amplified with the primers 5’-GGAGGTGGAGGTGGAGCTATGTCCGGTACGGGTGCC-3’ and 5’-TGCTTGAAAGGATCTTGCATCTAAAGCCCTCTTGTCAGATAG-3’ from N2 genomic DNA. The tbb-2 untranslated region (UTR) was PCR amplified with the primers 5’- ATGCAAGATCCTTTCAAGCA-3’ and 5’-TTCAAAGAAATCGCCGACTTCCAACCACATATGTTTCTCTTAGGC-3’ from N2 genomic DNA. The vector fragment used was the same as that used for the chrI vector above.

For *mex-5p*::Cherry::AIR-1::tbb-2 utr inserted in chr I, the coding sequence (CDS) region sequence was PCR amplified with the primers 5'-AGGGAGGTGGAGGTGGAGCTATGAGCGGAAAGGAAAATACTGC-3' and 5'-TGCTTGAAAGGATCTTGCATTTATTGATTGGCTGTAGAATTATTGCGAC-3' from N2 genomic DNA. The mex-5p::Cherry::tbb-2 utr vector was PCR amplified with the primers 5'-ATGCAAGATCCTTTCAAGCA-3' and 5'-ATAGCTCCACCTCCACCTCCCTTGTAAAGCTCATCCATTCCTCC-3' from shg1064 genomic DNA.

For *mex-5p*::mCherry::SAS-4::tbb-2 utr inserted in chr I, the coding sequence (CDS) region sequence was PCR amplified with the primers 5'-AGGGAGGTGGAGGTGGAGCTATGGCTTCCGATGAAAATATCGG -3' and 5'-TGCTTGAAAGGATCTTGCATTCATTTTTCCACTGGAACAAAGTTG -3' from N2 genomic DNA. The mex-5p::mCherry::tbb-2 utr backbone fragment is the same as that of *mex-5p*::mCherry::AIR-1::tbb-2 utr above.

For *mex-5p*::mCherry::SAS-5a::tbb-2 utr inserted in chr I, the coding sequence (CDS) region sequence was PCR amplified with the primers 5'-AGGGAGGTGGAGGTGGAGCTATGAATAATTACGACGACTTACCCT -3' and 5'-TGCTTGAAAGGATCTTGCATTCATTTCCTGCGAGCGTATTTTTC -3' from N2 genomic DNA. The mex-5p::mCherry::tbb-2 utr backbone fragment is the same as that of *mex-5p*::mCherry::AIR-1::tbb-2 utr above.

For *mex-5p*::mCherry::SAS-6::tbb-2 utr inserted in chr I, the coding sequence (CDS) region sequence was PCR amplified with the primers 5'-AGGGAGGTGGAGGTGGAGCTATGACTAGCAAAATTGCATTATTCG -3' and 5'-TGCTTGAAAGGATCTTGCATTTATCGTTGAGCGGGTGGGG -3' from N2 genomic DNA. The mex-5p::mCherry::tbb-2 utr backbone fragment is the same as that of *mex-5p*::mCherry::AIR-1::tbb-2 utr above.

For *mex-5p*::BFP::H2B::tbb-2 utr inserted in chr IV, the MEX-5 promoter was amplified with the primers 5'-CATCCCGTTAGAAACAATCTAAATATCAGTTTTTAAAAAATTAAACCAT -3' and 5'- TCCTTAATAAGCTCTGACATTCTCTGTCTGAAACATTCAATTG -3' from N2 genomic DNA. The BFP sequence was PCR amplified with the primers 5'-ATGTCAGAGCTTATTAAGGAGAATATG-3' and 5'-ATAGCTCCACCTCCACCTCCATTAAGCTTGTGACCCAGTTTGC-3' from shg1902 genomic DNA. The coding sequence (CDS) region and tbb-2 untranslated region (UTR) sequence were PCR amplified with the primers GGAGGTGGAGGTGGAGCTA-3 and GCTAACTTACATTTAGCTAGCCAACCACATATGTTTCTCTTAGGC-3 from shg366 genomic DNA. The chr IV vector fragment was PCR amplified with the primers 5'- CTAGCTAAATGTAAGTTAGCGACC -3' and 5'-AGATTGTTTCTAACGGGATGC -3' from the plasmid pCZGY2729.

**Supplemental figure legends**

**Figure S1. NRDE-3 is widely expressed in the germline, oocytes, early and late embryos and soma.** (A) The copy number of GFP in the indicated animals was quantified by real-time PCR. *eft-3* genomic DNA was used as an internal control for normalization. Mean ± SD. n = 3. (B) The percentage of larval arrest after feeding RNAi targeting the *lir-1* gene in the indicated animals (n > 50). Left: P0 generation, Right: F1 generation. (C) The percentage of dumpy animals after *dyp-11* RNAi in the indicated groups (n > 50). Left: P0 generation, Right: F1 generation. (D) Fluorescence micrographs of dissected gravid adult germlines from the indicated animals. The germline outline is marked. (E) Fluorescence micrographs of embryos from the indicated animals.

**Figure S2. NRDE-3 binds 22G RNAs.**

(A) Length and first letter distribution of NRDE-3-associated siRNAs in embryos from the indicated animals. (B) Venn diagram of the overlapping gene targets of GFP::NRDE-3(ggIS1, bombardment) and GFP::NRDE-3(ust574, KI) in embryos. Cutoff > 25 reads per million. (C-G) Venn diagram showing the overlap between GFP::NRDE-3(ust574, KI) targets and other known siRNA categories.

**Figure S3. siRNA binding was required for NRDE-3 accumulation in the perinuclear foci.** (A) Schematic of NRDE-3 and its variants. NLS, nuclear localization signal. (B, C) Fluorescence micrographs of the indicated embryos.

**Figure S4. Culturing animals at 25°C depleted the perinuclear accumulation of NRDE-3. (**A) Workflow of the temperature shift assay. Briefly, L4 stage animals grown at 20°C were divided onto new NGM plates and grown at 20°C or 25°C. Images were then taken 12 hours later, and the NRDE-3 foci were scored. (B, C, D) Left: Fluorescence micrographs of GFP::NRDE-3 embryos from the indicated animals. Right: The bar graph shows the percentages of NRDE-3 foci-positive embryos. An embryo with one or more NRDE-3 foci was defined as positive. At least 20 embryos were imaged for each condition.

**Figure S5. NRDE-3 accumulates in the peri-centrosomal foci.** (A, B) Images of *mex-5p::*GFP::NRDE-3 with the indicated mCherry-tagged centrosome proteins in the *eri-1* (A) *and ergo-1* (B) backgrounds.

**Figure S6. Culturing animals at 25°C depleted the peri-centrosomal accumulation of NRDE-3.** (A) Indicated animals were grown at 20°C and shifted to 25°C at the L4 stage. Images were taken after 12 hours of treatment. (B) Images of the indicated GFP::CSR-1 embryos.

**Figure S7. siRNA binding is essential for NRDE-3 accumulation in the peri-centrosomal foci and spindles during the cell cycle.** (A, B) Left: Fluorescence microscopy images of interphase and mitosis in the indicated animals. Right: Bar graph depicting the percentage of peri-centrosomal localization of NRDE-3(*PAZ) foci in each phase. Embryos at the 10-30-cell stage were selected for quantification. For each embryo, each cell was assigned to different mitosis phases using BFP::H2B as a marker. We defined one or more NRDE-3(*PAZ) foci in one cell as positive and then counted the percentage of positive cells in each phase. At least 50 cells were quantified for each phase.

**Figure S8. NRDE-3 (*NLS) accumulates in the peri-centrosomal foci and spindles during the cell cycle.** (A, B) Left: Fluorescence microscopy images of interphase and mitosis in the indicated animals. Right: Bar graph depicting the percentage of peri-centrosomal localization of NRDE-3(*NLS) foci in each phase. Embryos at the 10-30-cell stage were selected for quantification. For each embryo, each cell was assigned to different mitosis phases using BFP::H2B as a marker. We defined one or more NRDE-3(*NLS) foci in one cell as positive and then counted the percentage of positive cells in each phase. At least 50 cells were quantified for each phase.

**Figure S9. The peri-centrosomal-enriched siRNAs predominantly mapped to the 3' portion of the target genes.** (A-J) The distribution of normalized NRDE-3(*NLS)-associated small RNA reads across indicated genomic loci in wild type and *drh-3(ne4253)* animals.

**Figure S10. The integrity of the centrosome is required for the peri-centrosomal accumulation of NRDE-3.** (A, B) Images of *eri-1(-);*GFP::NRDE-3;mCherry::AIR-1 (A) and *ergo-1(-);*GFP::NRDE-3;mCherry::AIR-1 (B) animals after RNAi targeting of the centriole, and PCM genes. Synchronized embryos were hatched and cultured on NGM plates for 41 hours and then transferred to RNAi plates at the L4 stage. F1 embryos were imaged. (C) Bar graph quantifying the percentage of NRDE-3 foci-positive embryos. The embryos with one or more NRDE-3 foci were considered positive. At least 20 embryos were imaged for each RNAi experiment.

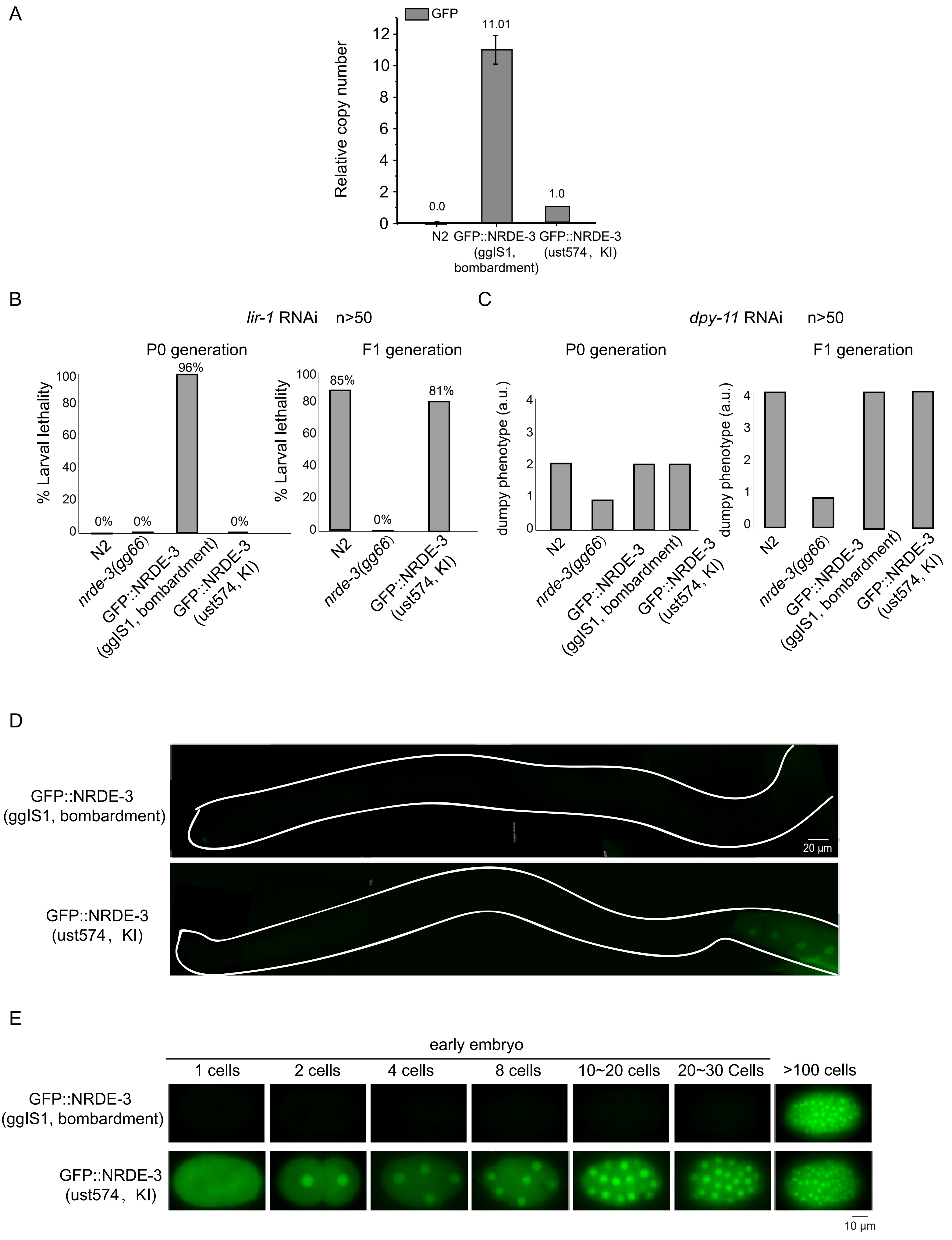
A
GFP
11.01
Relative copy number
0.0
1.0
N2
GFP::NRDE-3 (ggIS1, bombardment)
GFP::NRDE-3 (ust574，KI)
B
lir-1 RNAi n>50
P0 generation
F1 generation
% Larval lethality
96%
85%
81%
0%
nrde-3(gg66)
C
dpy-11 RNAi n>50
dumpy phenotype (a.u.)
D
20 μm
E
early embryo
1 cells
2 cells
4 cells
8 cells
10~20 cells
20~30 Cells
>100 cells
10 μm

Figure S1

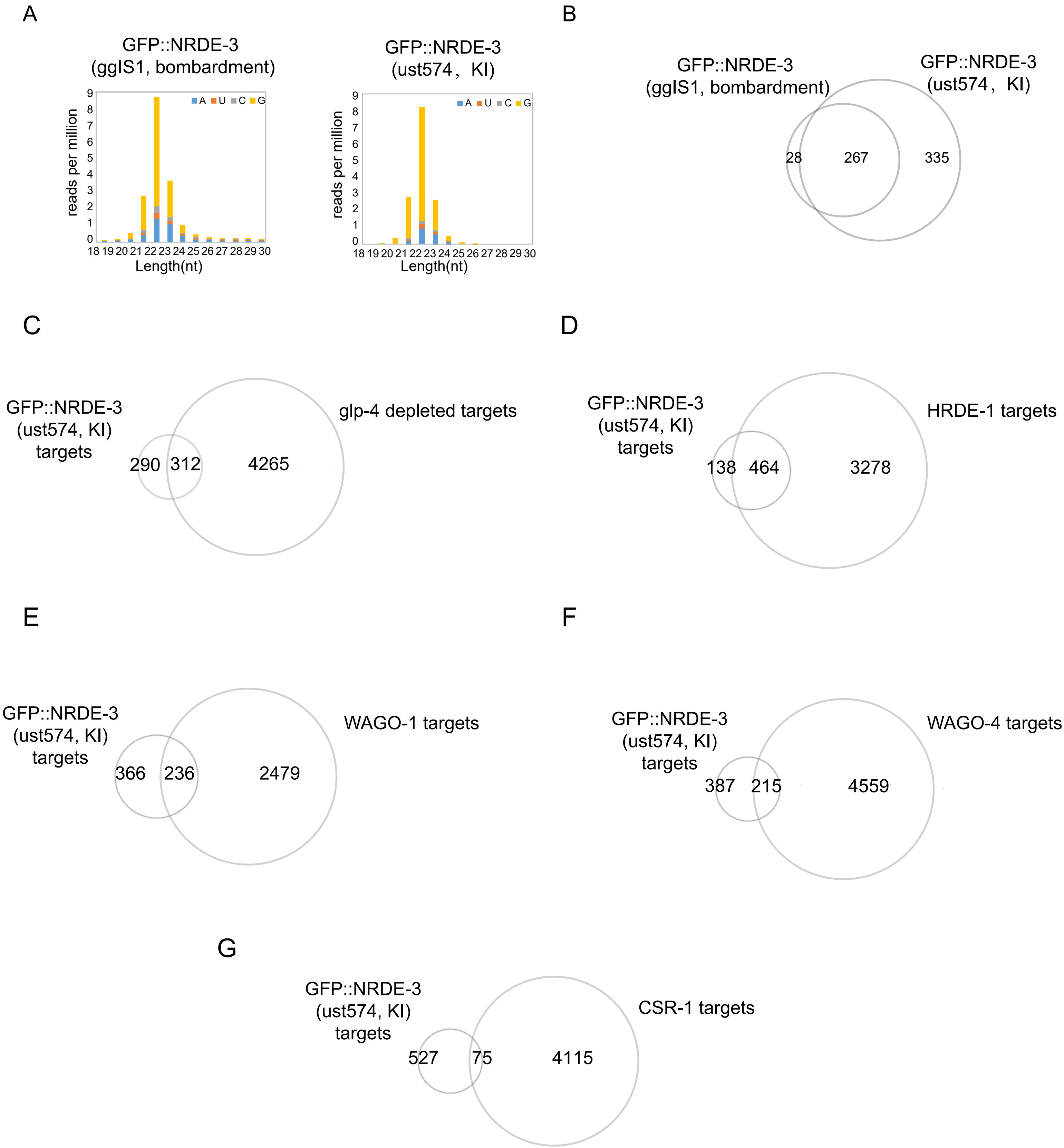
A
GFP::NRDE-3
(ggIS1, bombardment)
reads per million
Length(nt)
A U C G
GFP::NRDE-3
(ust574， KI)
B
GFP::NRDE-3
(ggIS1, bombardment)
GFP::NRDE-3
(ust574， KI)
28
267
335
C
GFP::NRDE-3
(ust574, KI)
targets
glp-4 depleted targets
290
312
4265
D
GFP::NRDE-3
(ust574, KI)
targets
HRDE-1 targets
138
464
3278
E
GFP::NRDE-3
(ust574, KI)
targets
WAGO-1 targets
366
236
2479
F
GFP::NRDE-3
(ust574, KI)
targets
WAGO-4 targets
387
215
4559
G
GFP::NRDE-3
(ust574, KI)
targets
CSR-1 targets
527
75
4115

Figure S2

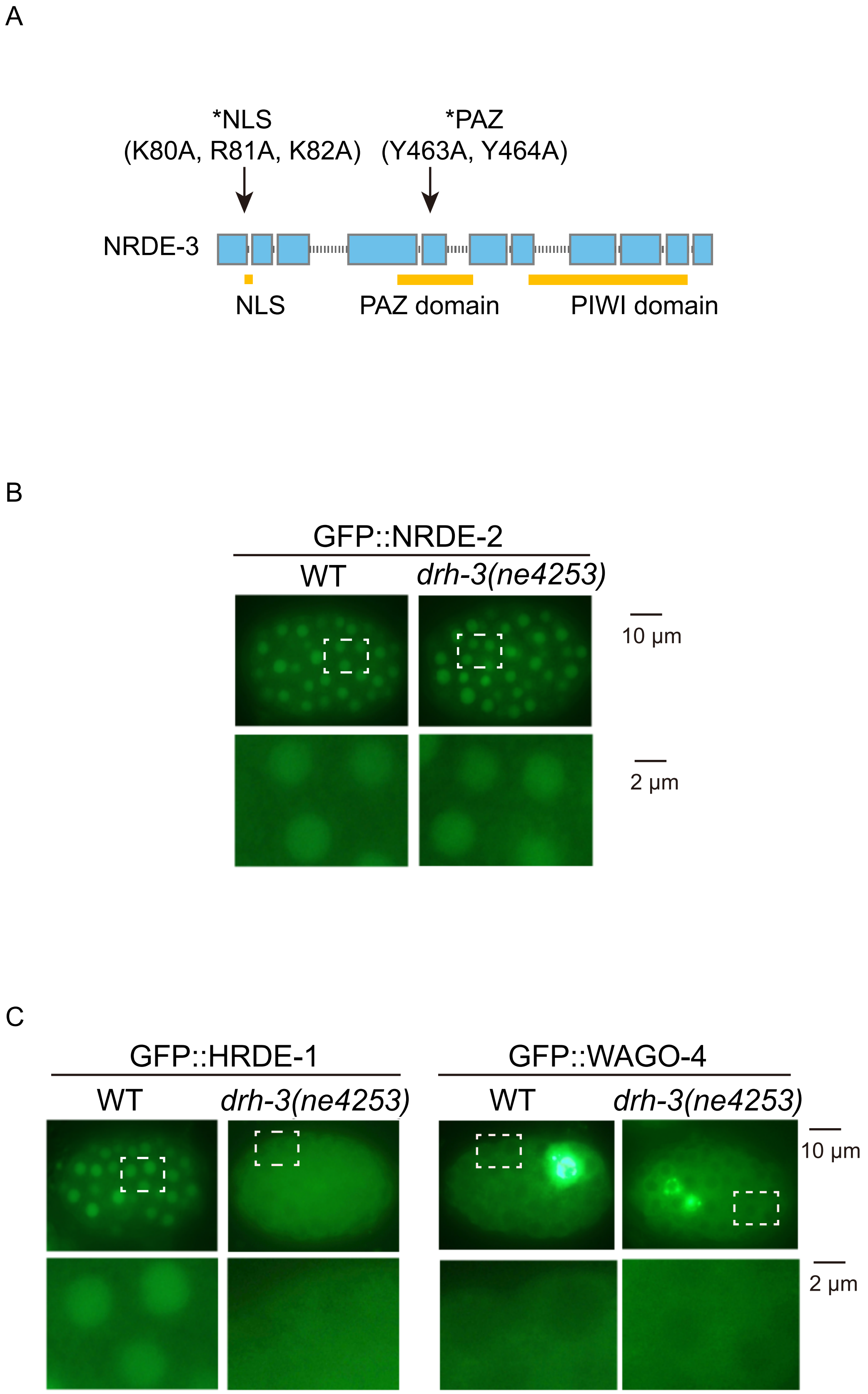
A
*NLS
(K80A, R81A, K82A)
*PAZ
(Y463A, Y464A)
NRDE-3
NLS
PAZ domain
PIWI domain
B
GFP::NRDE-2
WT
drh-3(ne4253)
10 μm
2 μm
C
GFP::HRDE-1
GFP::WAGO-4
WT
drh-3(ne4253)
WT
drh-3(ne4253)
10 μm
2 μm

Figure S3

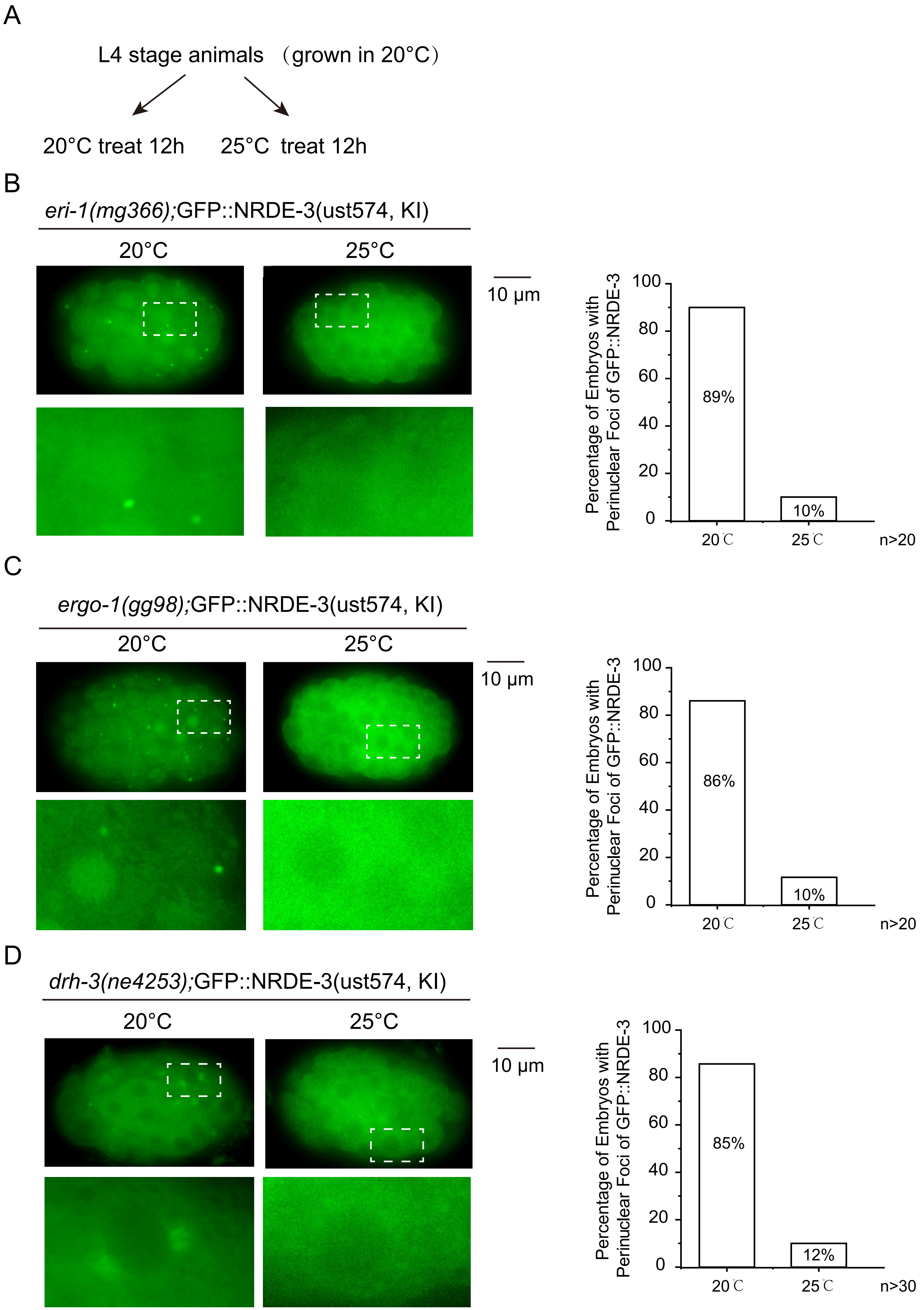
A
L4 stage animals (grown in 20°C)
20°C treat 12h
25°C treat 12h
B
eri-1(mg366);GFP::NRDE-3(ust574, KI)
20°C
25°C
10 μm
Percentage of Embryos with Perinuclear Foci of GFP::NRDE-3
100
80
60
40
20
0
89%
10%
20℃
25℃
n>20
C
ergo-1(gg98);GFP::NRDE-3(ust574, KI)
20°C
25°C
10 μm
Percentage of Embryos with Perinuclear Foci of GFP::NRDE-3
100
80
60
40
20
0
86%
10%
20℃
25℃
n>20
D
drh-3(ne4253);GFP::NRDE-3(ust574, KI)
20°C
25°C
10 μm
Percentage of Embryos with Perinuclear Foci of GFP::NRDE-3
100
80
60
40
20
0
85%
12%
20℃
25℃
n>30

Figure S4

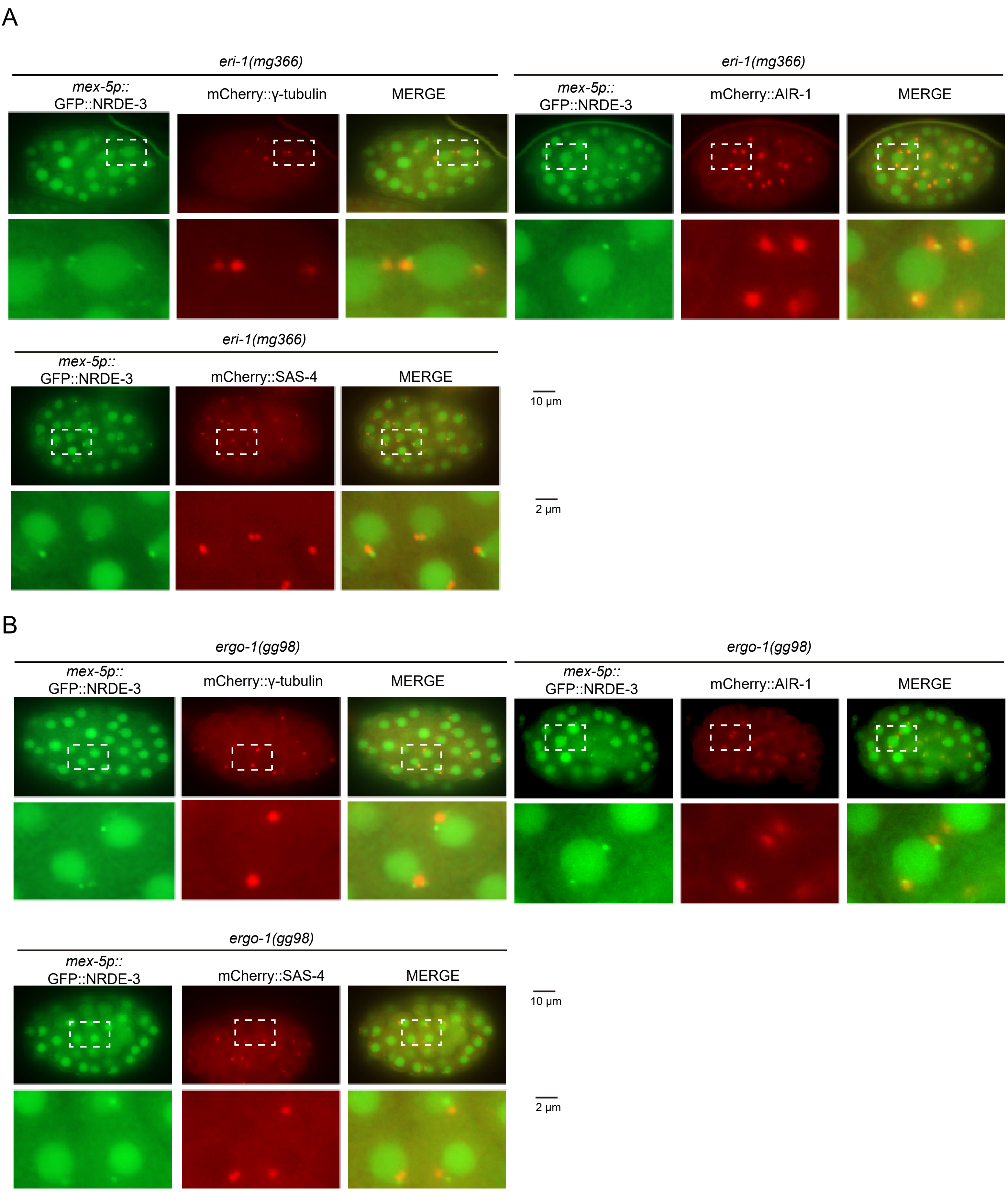
A
eri-1(mg366)
mex-5p::
GFP::NRDE-3
mCherry::γ-tubulin
MERGE
eri-1(mg366)
mex-5p::
GFP::NRDE-3
mCherry::AIR-1
MERGE
eri-1(mg366)
mex-5p::
GFP::NRDE-3
mCherry::SAS-4
MERGE
10 μm
2 μm
B
ergo-1(gg98)
mex-5p::
GFP::NRDE-3
mCherry::γ-tubulin
MERGE
ergo-1(gg98)
mex-5p::
GFP::NRDE-3
mCherry::AIR-1
MERGE
ergo-1(gg98)
mex-5p::
GFP::NRDE-3
mCherry::SAS-4
MERGE
10 μm
2 μm

Figure S5

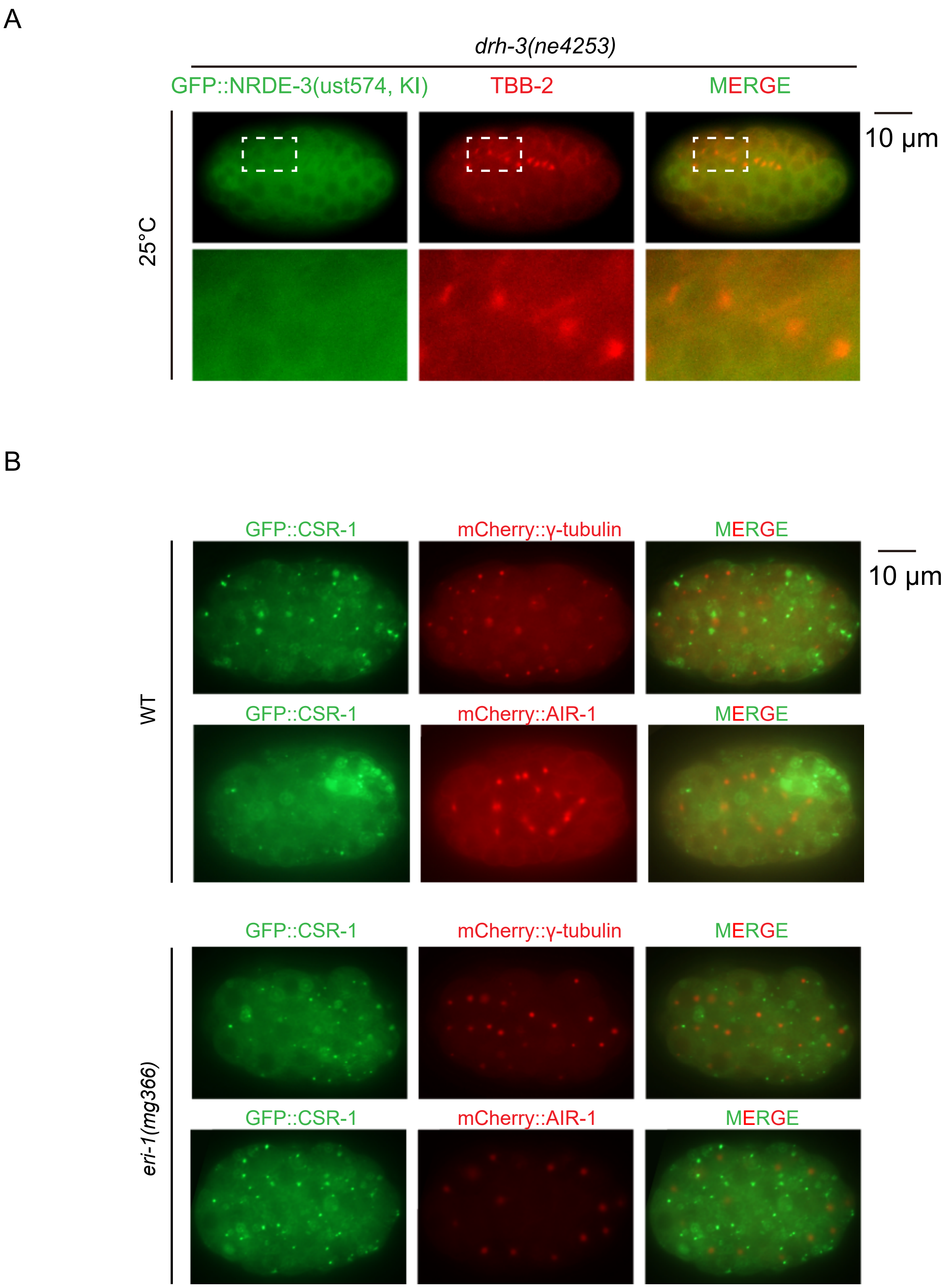
A
drh-3(ne4253)
GFP::NRDE-3(ust574, KI)
TBB-2
MERGE
10 μm
25°C
B
GFP::CSR-1
mCherry::γ-tubulin
MERGE
10 μm
WT
GFP::CSR-1
mCherry::AIR-1
MERGE
GFP::CSR-1
mCherry::γ-tubulin
MERGE
eri-1(mg366)
GFP::CSR-1
mCherry::AIR-1
MERGE

Figure S6

A

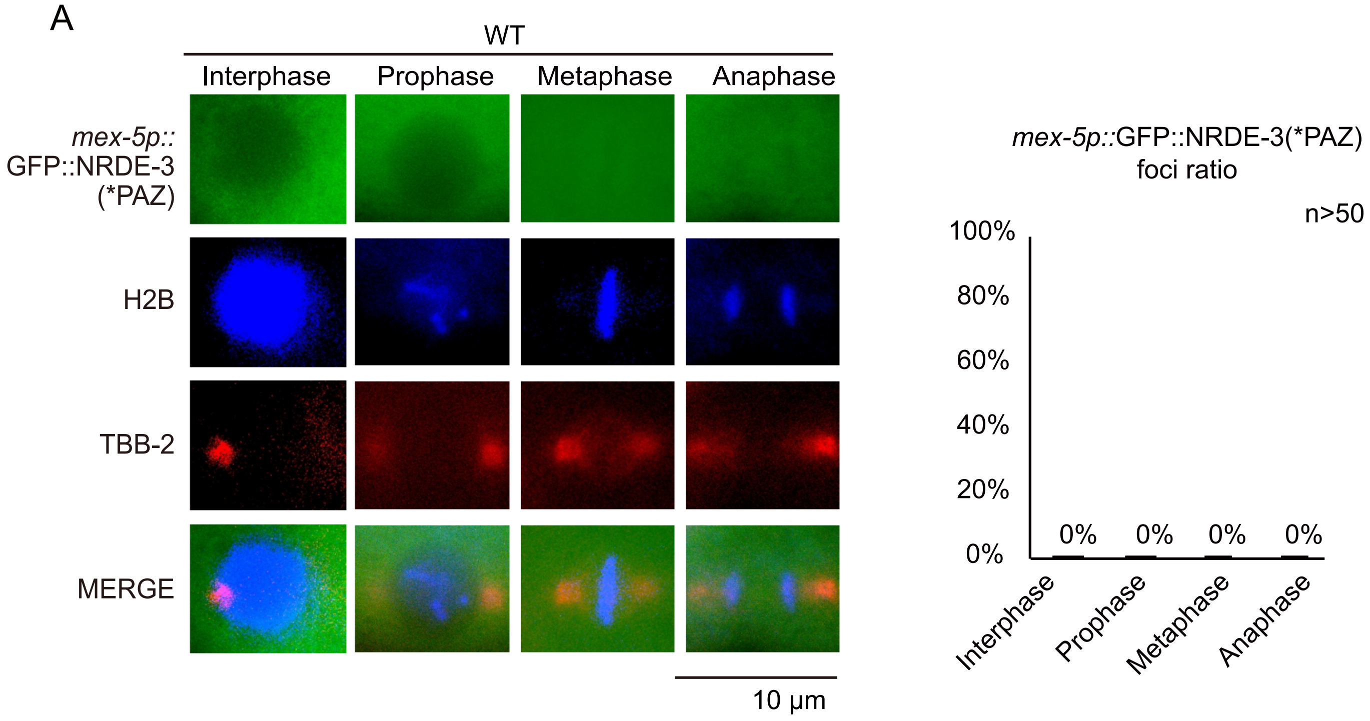
WT
Interphase
Prophase
Metaphase
Anaphase
mex-5p::
GFP::NRDE-3
(*PAZ)
H2B
TBB-2
MERGE
10 μm
mex-5p::GFP::NRDE-3(*PAZ)
foci ratio
n>50
100%
80%
60%
40%
20%
0%
0%
0%
0%
0%
Interphase
Prophase
Metaphase
Anaphase

B

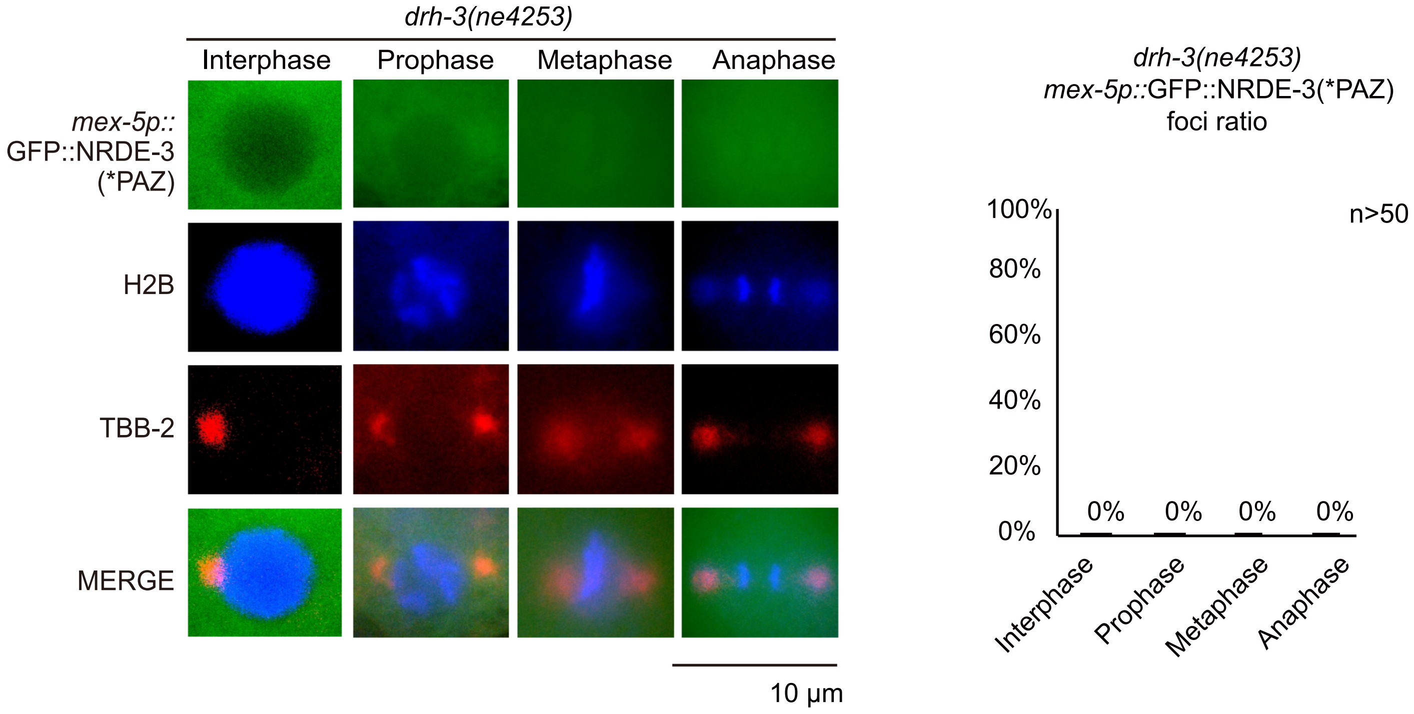
drh-3(ne4253)
Interphase
Prophase
Metaphase
Anaphase
mex-5p::
GFP::NRDE-3
(*PAZ)
H2B
TBB-2
MERGE
10 μm
drh-3(ne4253)
mex-5p::GFP::NRDE-3(*PAZ)
foci ratio
n>50
100%
80%
60%
40%
20%
0%
0%
0%
0%
0%
Interphase
Prophase
Metaphase
Anaphase

Figure S7

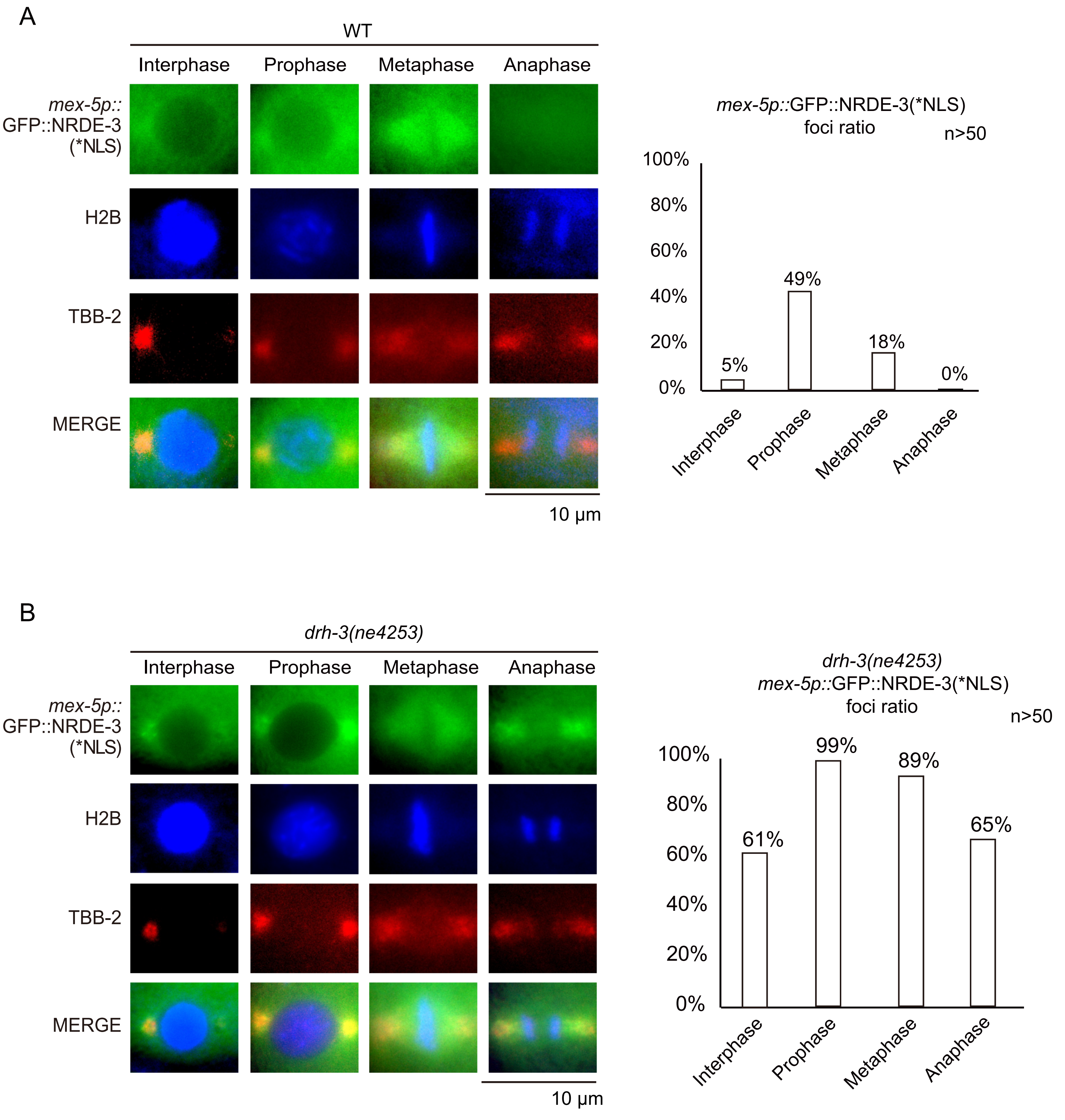
A
WT
Interphase
Prophase
Metaphase
Anaphase
mex-5p::
GFP::NRDE-3
(*NLS)
H2B
TBB-2
MERGE
10 µm
mex-5p::GFP::NRDE-3(*NLS)
foci ratio
n>50
100%
80%
60%
40%
20%
0%
5%
49%
18%
0%
B
drh-3(ne4253)
drh-3(ne4253)
mex-5p::GFP::NRDE-3(*NLS)
foci ratio
61%
99%
89%
65%

Figure S8

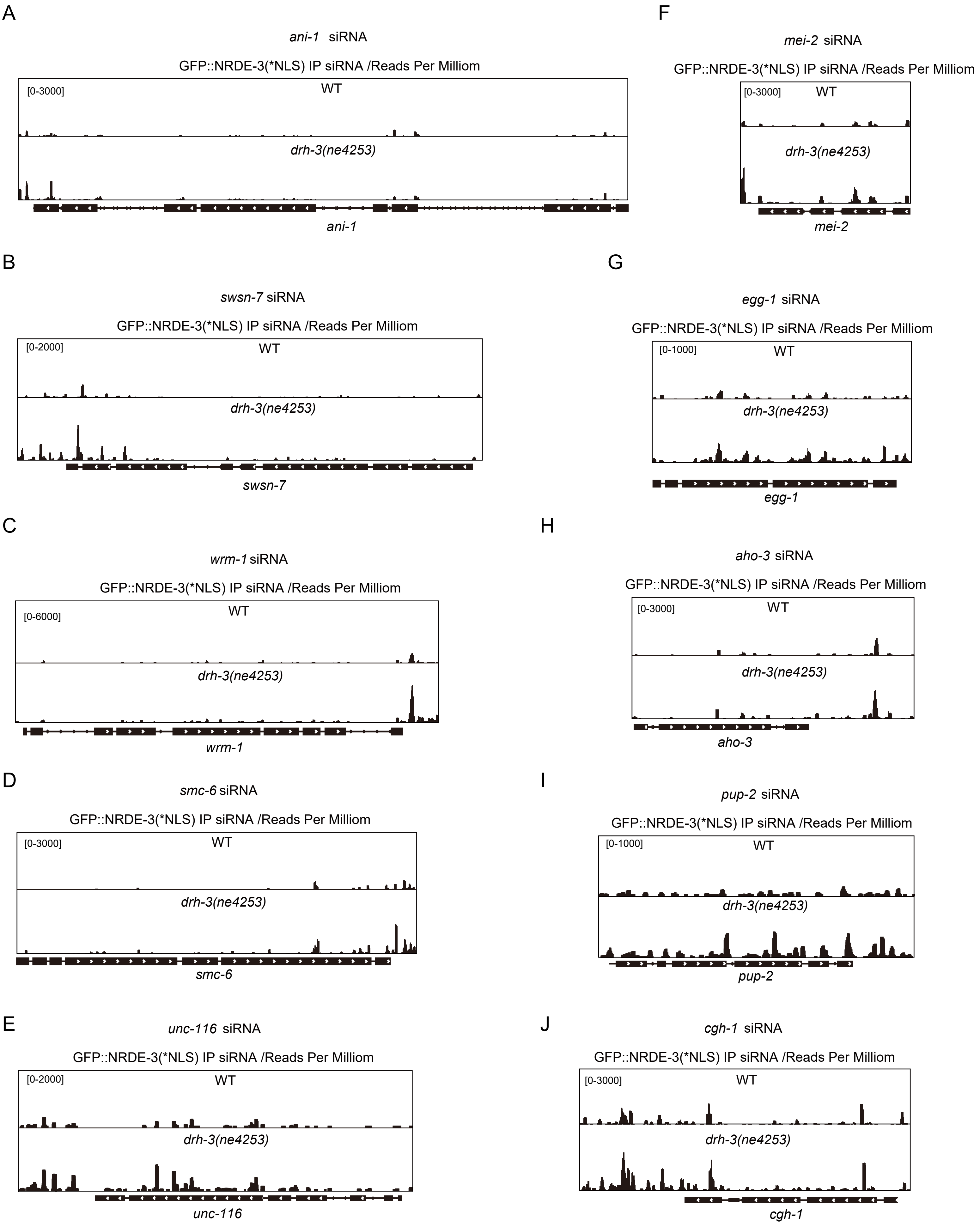
A
ani-1 siRNA
GFP::NRDE-3(*NLS) IP siRNA /Reads Per Milliom
[0-3000]
WT
drh-3(ne4253)
ani-1
B
swsn-7 siRNA
GFP::NRDE-3(*NLS) IP siRNA /Reads Per Milliom
[0-2000]
WT
drh-3(ne4253)
swsn-7
C
wrm-1 siRNA
GFP::NRDE-3(*NLS) IP siRNA /Reads Per Milliom
[0-6000]
WT
drh-3(ne4253)
wrm-1
D
smc-6 siRNA
GFP::NRDE-3(*NLS) IP siRNA /Reads Per Milliom
[0-3000]
WT
drh-3(ne4253)
smc-6
E
unc-116 siRNA
GFP::NRDE-3(*NLS) IP siRNA /Reads Per Milliom
[0-2000]
WT
drh-3(ne4253)
unc-116
F
mei-2 siRNA
GFP::NRDE-3(*NLS) IP siRNA /Reads Per Milliom
[0-3000]
WT
drh-3(ne4253)
mei-2
G
egg-1 siRNA
GFP::NRDE-3(*NLS) IP siRNA /Reads Per Milliom
[0-1000]
WT
drh-3(ne4253)
egg-1
H
aho-3 siRNA
GFP::NRDE-3(*NLS) IP siRNA /Reads Per Milliom
[0-3000]
WT
drh-3(ne4253)
aho-3
I
pup-2 siRNA
GFP::NRDE-3(*NLS) IP siRNA /Reads Per Milliom
[0-1000]
WT
drh-3(ne4253)
pup-2
J
cgh-1 siRNA
GFP::NRDE-3(*NLS) IP siRNA /Reads Per Milliom
[0-3000]
WT
drh-3(ne4253)
cgh-1

Figure S9

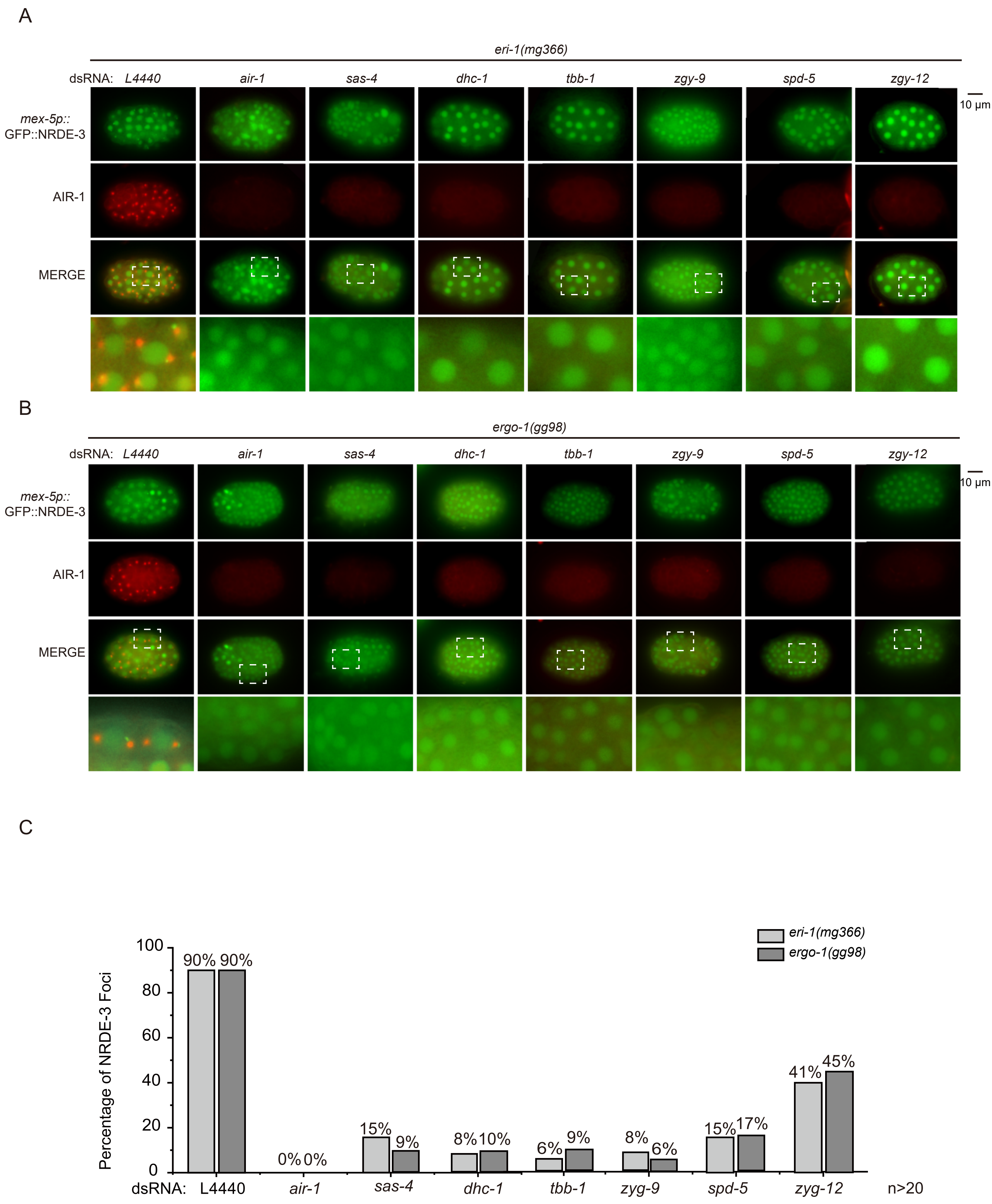
A
eri-1(mg366)
dsRNA: L4440
air-1
sas-4
dhc-1
tbb-1
zgy-9
spd-5
zgy-12
10 μm
mex-5p::
GFP::NRDE-3
AIR-1
MERGE
B
ergo-1(gg98)
dsRNA: L4440
air-1
sas-4
dhc-1
tbb-1
zgy-9
spd-5
zgy-12
10 μm
mex-5p::
GFP::NRDE-3
AIR-1
MERGE
C
eri-1(mg366)
ergo-1(gg98)
Percentage of NRDE-3 Foci
100
80
60
40
20
0
90%
90%
0%
0%
15%
9%
8%
10%
6%
9%
8%
6%
15%
17%
41%
45%
dsRNA: L4440
air-1
sas-4
dhc-1
tbb-1
zyg-9
spd-5
zyg-12
n>20

Figure S10

**Table S1. Peri-centrosomal-enriched siRNA targets.**

**Table S2. List of strains used in this study.**

**Table S3. Sequence of sgRNAs for CRISPR/Cas9-mediated gene editing.**

**Table S4. Sequence of quantitative real-time PCR primers.**

**Table S1. Peri-centrosomal-enriched siRNA targets.**

| Transcript | NRDE-3(*NLS) associated siRNA | | | TBB-2 associated siRNA | | |
|---|---|---|---|---|---|---|
| | WT / reads per million | *drh-3(ne4253)* / reads per million | *drh-3(ne4253)* / WT fold change | WT / reads per million | *drh-3(ne4253)* / reads per million | *drh-3(ne4253)* / WT fold change |
| ZK930.13 | 58.57 | 315.09 | 5.38 | 7001.50 | 13229.19 | 1.89 |
| F31C3.7 | 73.09 | 360.93 | 4.94 | 89.82 | 211.80 | 2.36 |
| F31C3.8 | 73.67 | 360.76 | 4.90 | 91.92 | 223.60 | 2.43 |
| D2045.6 | 162.44 | 630.62 | 3.88 | 27.54 | 49.38 | 1.79 |
| F39D8.7 | 398.06 | 1467.71 | 3.69 | 222.75 | 1106.83 | 4.97 |
| R09A1.1 | 84.51 | 297.55 | 3.52 | 26.65 | 47.20 | 1.77 |
| F29D11.2 | 287.42 | 948.98 | 3.30 | 41.02 | 113.98 | 2.78 |
| F31C3.11 | 47.24 | 153.50 | 3.25 | 25.15 | 96.58 | 3.84 |
| F01G4.1 | 171.73 | 553.65 | 3.22 | 37.43 | 72.98 | 1.95 |
| T10F2.3 | 296.42 | 939.26 | 3.17 | 25.15 | 91.93 | 3.66 |
| R10E8.6 | 207.74 | 641.54 | 3.09 | 317.37 | 1545.96 | 4.87 |
| ZK177.6 | 111.71 | 343.73 | 3.08 | 52.10 | 168.63 | 3.24 |
| Y71F9AL.18 | 126.23 | 387.76 | 3.07 | 51.80 | 117.08 | 2.26 |
| D1081.8 | 81.22 | 248.70 | 3.06 | 33.83 | 55.90 | 1.65 |
| K10D2.2 | 321.49 | 973.06 | 3.03 | 28.44 | 128.26 | 4.51 |
| R05D3.7 | 774.64 | 2321.32 | 3.00 | 77.25 | 413.35 | 5.35 |
| C08B11.3 | 440.66 | 1316.62 | 2.99 | 67.66 | 208.07 | 3.08 |
| W05F2.3 | 86.74 | 257.82 | 2.97 | 35.63 | 108.70 | 3.05 |
| Y42H9AR.4 | 267.57 | 791.43 | 2.96 | 34.13 | 109.94 | 3.22 |
| F57B10.12 | 352.95 | 1040.82 | 2.95 | 27.54 | 119.88 | 4.35 |
| F53H2.4 | 62.05 | 181.63 | 2.93 | 26.95 | 68.32 | 2.54 |
| K01F9.7 | 35.33 | 101.05 | 2.86 | 825.45 | 2232.92 | 2.71 |
| K09H9.7 | 208.52 | 595.27 | 2.85 | 40.42 | 54.04 | 1.34 |
| C36C9.10 | 348.40 | 981.31 | 2.82 | 78.74 | 403.73 | 5.13 |
| C07H6.5 | 854.11 | 2405.60 | 2.82 | 93.41 | 391.61 | 4.19 |
| F55A11.7 | 247.43 | 692.45 | 2.80 | 40.12 | 73.60 | 1.83 |
| C55B7.1 | 180.64 | 503.43 | 2.79 | 26.05 | 74.53 | 2.86 |
| K04G2.2 | 394.19 | 1089.76 | 2.76 | 36.53 | 101.55 | 2.78 |
| Y56A3A.20 | 113.55 | 313.63 | 2.76 | 38.02 | 80.75 | 2.12 |
| W03G9.2 | 116.94 | 321.11 | 2.75 | 54.79 | 86.96 | 1.59 |
| F01F1.4 | 257.41 | 706.21 | 2.74 | 48.50 | 184.16 | 3.80 |
| F26D10.10 | 105.71 | 289.81 | 2.74 | 35.33 | 66.77 | 1.89 |
| Y41D4B.19b | 80.54 | 216.80 | 2.69 | 36.23 | 55.28 | 1.53 |
| C43E11.1 | 94.00 | 252.83 | 2.69 | 66.17 | 106.52 | 1.61 |
| T09B4.14 | 54.60 | 146.71 | 2.69 | 413.77 | 934.47 | 2.26 |

| | | | | | | |
|---|---|---|---|---|---|---|
| R05D11.8 | 109.20 | 291.36 | 2.67 | 28.74 | 121.12 | 4.21 |
| B0244.8 | 318.59 | 843.80 | 2.65 | 27.84 | 160.87 | 5.78 |
| C32D5.10 | 313.36 | 829.61 | 2.65 | 58.98 | 104.35 | 1.77 |
| ZK675.1.1 | 214.91 | 568.27 | 2.64 | 32.04 | 126.09 | 3.94 |
| F23A7.8 | 151.69 | 397.05 | 2.62 | 55.69 | 111.18 | 2.00 |
| T01G9.5a | 118.10 | 307.10 | 2.60 | 27.25 | 48.76 | 1.79 |
| F54D5.14 | 616.94 | 1592.32 | 2.58 | 94.01 | 331.06 | 3.52 |
| Y66D12A.5 | 61.76 | 157.98 | 2.56 | 25.15 | 39.44 | 1.57 |
| VF39H2L.1 | 103.97 | 265.56 | 2.55 | 26.35 | 86.96 | 3.30 |
| C17H12.1 | 162.83 | 414.68 | 2.55 | 42.81 | 118.94 | 2.78 |
| T06D10.2 | 90.22 | 229.61 | 2.54 | 27.54 | 90.37 | 3.28 |
| Y49E10.19 | 304.45 | 774.58 | 2.54 | 54.19 | 199.69 | 3.68 |
| C01F1.2 | 55.66 | 141.21 | 2.54 | 25.15 | 50.93 | 2.03 |
| B0336.1 | 949.66 | 2401.98 | 2.53 | 73.95 | 236.65 | 3.20 |
| F10E7.8 | 504.16 | 1271.47 | 2.52 | 30.84 | 115.53 | 3.75 |
| F08B4.5 | 419.46 | 1057.59 | 2.52 | 56.59 | 121.74 | 2.15 |
| T24F1.2 | 245.60 | 617.29 | 2.51 | 49.10 | 82.61 | 1.68 |
| Y37A1B.1a | 154.11 | 387.24 | 2.51 | 30.54 | 63.66 | 2.08 |
| ZK675.1.2 | 174.73 | 435.23 | 2.49 | 25.75 | 124.84 | 4.85 |
| C36A4.4 | 110.07 | 272.09 | 2.47 | 30.54 | 108.70 | 3.56 |
| K06A5.7.1 | 245.40 | 606.19 | 2.47 | 29.04 | 95.65 | 3.29 |
| K06A5.7.2 | 246.47 | 607.66 | 2.47 | 36.53 | 96.58 | 2.64 |
| T07A9.6 | 335.91 | 824.11 | 2.45 | 70.06 | 205.90 | 2.94 |
| C10H11.9 | 463.02 | 1131.81 | 2.44 | 68.56 | 127.95 | 1.87 |
| F32A5.1b | 179.38 | 436.69 | 2.43 | 28.74 | 53.11 | 1.85 |
| M04B2.1 | 870.18 | 2097.04 | 2.41 | 130.54 | 399.07 | 3.06 |
| F25D7.4 | 584.90 | 1407.43 | 2.41 | 199.40 | 300.00 | 1.50 |
| M02B7.5a | 220.43 | 529.23 | 2.40 | 42.51 | 128.26 | 3.02 |
| R11A5.2 | 420.91 | 1003.24 | 2.38 | 84.43 | 213.04 | 2.52 |
| K01G5.5 | 115.20 | 272.78 | 2.37 | 33.23 | 81.99 | 2.47 |
| C02F5.1 | 354.70 | 839.59 | 2.37 | 102.10 | 161.80 | 1.58 |
| T11F8.3a.1 | 352.66 | 833.40 | 2.36 | 62.28 | 287.89 | 4.62 |
| F32A5.1a | 184.51 | 435.32 | 2.36 | 30.54 | 52.17 | 1.71 |
| Y65B4BL.5a | 84.51 | 199.00 | 2.35 | 28.44 | 55.90 | 1.97 |
| T07C4.3b.2 | 127.01 | 298.67 | 2.35 | 28.74 | 85.71 | 2.98 |
| Y53C12A.1 | 606.00 | 1419.73 | 2.34 | 61.68 | 193.17 | 3.13 |
| C28A5.2 | 265.54 | 620.47 | 2.34 | 75.75 | 105.28 | 1.39 |
| F10G7.2 | 319.07 | 744.65 | 2.33 | 80.84 | 269.88 | 3.34 |
| Y47G6A.18 | 57.99 | 135.27 | 2.33 | 45.21 | 67.39 | 1.49 |
| T07C4.3a.2 | 133.20 | 309.25 | 2.32 | 27.84 | 82.61 | 2.97 |
| T11F8.3b.1 | 342.21 | 792.55 | 2.32 | 61.98 | 289.75 | 4.68 |
| M7.2b | 173.86 | 401.00 | 2.31 | 26.05 | 54.66 | 2.10 |
| EEED8.5 | 125.75 | 289.98 | 2.31 | 29.34 | 44.72 | 1.52 |
| C16A3.7 | 369.12 | 850.00 | 2.30 | 51.50 | 143.17 | 2.78 |

| | | | | | | |
|---|---|---|---|---|---|---|
| T05H10.5b | 201.36 | 463.61 | 2.30 | 29.64 | 47.83 | 1.61 |
| CD4.17 | 104.16 | 239.16 | 2.30 | 113.17 | 229.81 | 2.03 |
| F56F3.1 | 167.18 | 382.09 | 2.29 | 53.29 | 77.64 | 1.46 |
| B0379.3a | 115.68 | 263.41 | 2.28 | 32.34 | 85.40 | 2.64 |
| T04A8.14 | 459.05 | 1044.26 | 2.27 | 92.22 | 193.17 | 2.09 |
| Y75B12B.2.2 | 117.23 | 266.59 | 2.27 | 64.07 | 155.28 | 2.42 |
| Y75B12B.2.1 | 125.94 | 284.91 | 2.26 | 65.27 | 174.22 | 2.67 |
| Y47G6A.28 | 579.96 | 1310.42 | 2.26 | 62.87 | 225.78 | 3.59 |
| C30B5.1 | 80.15 | 180.85 | 2.26 | 28.74 | 62.11 | 2.16 |
| C28H8.9a | 81.12 | 182.83 | 2.25 | 29.34 | 43.48 | 1.48 |
| F20G4.3 | 482.77 | 1085.63 | 2.25 | 125.45 | 309.01 | 2.46 |
| T07C4.3b.1 | 132.14 | 296.69 | 2.25 | 30.24 | 87.58 | 2.90 |
| F10E9.8 | 524.49 | 1174.63 | 2.24 | 56.59 | 113.35 | 2.00 |
| Y111B2A.11 | 134.75 | 301.59 | 2.24 | 33.53 | 50.31 | 1.50 |
| B0379.3b | 115.39 | 257.91 | 2.24 | 35.93 | 86.65 | 2.41 |
| F54E12.2.2 | 278.32 | 622.02 | 2.23 | 64.07 | 176.71 | 2.76 |
| W09D10.1.1 | 187.03 | 417.09 | 2.23 | 28.14 | 58.39 | 2.07 |
| T11F8.3b.2 | 355.37 | 789.02 | 2.22 | 63.17 | 281.06 | 4.45 |
| K12D12.5 | 259.05 | 574.37 | 2.22 | 35.33 | 81.06 | 2.29 |
| D1037.1 | 45.21 | 100.01 | 2.21 | 32.04 | 43.48 | 1.36 |
| F55A3.3 | 339.69 | 751.10 | 2.21 | 72.46 | 121.74 | 1.68 |
| T26A8.4 | 394.68 | 870.72 | 2.21 | 49.70 | 72.05 | 1.45 |
| F33H2.1.2 | 159.63 | 350.78 | 2.20 | 41.62 | 59.63 | 1.43 |
| T05C3.5 | 143.95 | 315.78 | 2.19 | 41.32 | 67.39 | 1.63 |
| F35G12.8 | 295.84 | 648.76 | 2.19 | 108.98 | 167.70 | 1.54 |
| T19E7.3a | 137.46 | 300.90 | 2.19 | 47.31 | 84.78 | 1.79 |
| F54E12.2.1 | 281.80 | 616.77 | 2.19 | 57.49 | 183.85 | 3.20 |
| M02B7.5b | 218.49 | 477.97 | 2.19 | 28.74 | 125.78 | 4.38 |
| T11F8.3a.2 | 368.25 | 805.45 | 2.19 | 51.80 | 302.48 | 5.84 |
| F33H1.4 | 136.40 | 298.15 | 2.19 | 40.12 | 66.77 | 1.66 |
| C02F4.1 | 148.60 | 323.87 | 2.18 | 38.32 | 75.47 | 1.97 |
| M176.2 | 155.66 | 338.66 | 2.18 | 29.64 | 62.73 | 2.12 |
| T05F1.6 | 161.86 | 351.30 | 2.17 | 26.95 | 62.11 | 2.31 |
| R144.2a | 426.04 | 921.89 | 2.16 | 39.22 | 113.66 | 2.90 |
| T07C4.3a.1 | 139.59 | 302.02 | 2.16 | 25.15 | 85.09 | 3.38 |
| Y110A7A.17a | 566.31 | 1219.44 | 2.15 | 79.64 | 309.32 | 3.88 |
| R08D7.6b | 485.00 | 1043.83 | 2.15 | 38.62 | 117.39 | 3.04 |
| T05H10.5a | 211.42 | 454.24 | 2.15 | 26.95 | 50.31 | 1.87 |
| Y24D9A.2 | 501.16 | 1074.71 | 2.14 | 56.59 | 144.72 | 2.56 |
| F57C9.7 | 139.21 | 297.98 | 2.14 | 25.15 | 39.75 | 1.58 |
| C18G1.5 | 183.83 | 393.35 | 2.14 | 64.67 | 106.21 | 1.64 |
| W09G10.4a | 64.86 | 138.46 | 2.13 | 26.95 | 53.11 | 1.97 |
| T25E12.5 | 128.36 | 273.90 | 2.13 | 55.39 | 90.68 | 1.64 |

| | | | | | | |
|---|---|---|---|---|---|---|
| W09D10.1.2 | 196.61 | 419.32 | 2.13 | 29.34 | 58.70 | 2.00 |
| F33H2.1.1 | 164.38 | 350.52 | 2.13 | 38.32 | 53.42 | 1.39 |
| C50F2.2 | 126.23 | 268.83 | 2.13 | 70.36 | 130.43 | 1.85 |
| F28B3.7a | 214.42 | 455.87 | 2.13 | 32.63 | 90.06 | 2.76 |
| Y76B12C.2 | 151.50 | 322.06 | 2.13 | 34.13 | 62.73 | 1.84 |
| ZK1098.8 | 310.07 | 659.00 | 2.13 | 64.37 | 106.83 | 1.66 |
| Y71F9AL.9 | 254.11 | 539.98 | 2.12 | 76.35 | 181.99 | 2.38 |
| K10B2.1 | 546.56 | 1159.50 | 2.12 | 42.81 | 158.39 | 3.70 |
| C27D9.1 | 349.56 | 739.83 | 2.12 | 61.38 | 97.52 | 1.59 |
| C17E4.3 | 129.91 | 274.76 | 2.11 | 29.94 | 59.01 | 1.97 |
| ZK328.2 | 113.94 | 240.62 | 2.11 | 35.33 | 56.52 | 1.60 |
| ZK265.6 | 33.30 | 70.26 | 2.11 | 26.35 | 36.02 | 1.37 |
| T24D1.3 | 669.12 | 1409.49 | 2.11 | 95.81 | 302.17 | 3.15 |
| C28C12.2 | 2220.23 | 4674.97 | 2.11 | 440.12 | 620.81 | 1.41 |
| VW02B12L.3 | 216.36 | 455.53 | 2.11 | 36.23 | 76.40 | 2.11 |
| F58A4.3 | 406.20 | 853.87 | 2.10 | 143.41 | 301.55 | 2.10 |
| C01C7.1b | 204.65 | 430.16 | 2.10 | 47.60 | 74.84 | 1.57 |
| C32F10.5 | 300.00 | 630.44 | 2.10 | 29.64 | 77.64 | 2.62 |
| F23A7.4.1 | 108.23 | 227.38 | 2.10 | 40.42 | 66.46 | 1.64 |
| K04G2.8a | 313.55 | 657.62 | 2.10 | 37.13 | 113.04 | 3.04 |
| R06A4.2 | 566.70 | 1188.39 | 2.10 | 76.35 | 123.60 | 1.62 |
| R08D7.6a | 497.00 | 1042.11 | 2.10 | 41.32 | 120.81 | 2.92 |
| F25H2.8 | 107.84 | 225.05 | 2.09 | 34.13 | 54.66 | 1.60 |
| T19E10.1b | 174.15 | 362.91 | 2.08 | 36.23 | 69.88 | 1.93 |
| C32D5.11 | 83.74 | 174.49 | 2.08 | 25.75 | 34.16 | 1.33 |
| F30A10.10 | 386.25 | 804.42 | 2.08 | 54.49 | 126.09 | 2.31 |
| F26H9.2 | 207.94 | 432.65 | 2.08 | 34.43 | 66.15 | 1.92 |
| K04G2.8b | 318.97 | 662.52 | 2.08 | 33.23 | 107.45 | 3.23 |
| F56C9.11 | 87.12 | 179.82 | 2.06 | 27.54 | 40.06 | 1.45 |
| T19E10.1a | 173.86 | 358.44 | 2.06 | 30.84 | 69.57 | 2.26 |
| W09C5.2 | 195.64 | 402.98 | 2.06 | 28.14 | 56.83 | 2.02 |
| F40F11.2 | 654.11 | 1346.46 | 2.06 | 119.46 | 313.04 | 2.62 |
| C07G2.1a | 202.61 | 415.71 | 2.05 | 85.93 | 212.42 | 2.47 |
| ZK632.12 | 175.70 | 360.33 | 2.05 | 66.17 | 194.41 | 2.94 |
| C56E6.3a | 117.72 | 240.88 | 2.05 | 25.45 | 38.20 | 1.50 |
| R03D7.4 | 146.18 | 298.75 | 2.04 | 34.13 | 68.63 | 2.01 |
| F56D2.6a | 189.16 | 386.38 | 2.04 | 32.63 | 43.48 | 1.33 |
| F08F8.8 | 297.97 | 607.14 | 2.04 | 50.00 | 217.08 | 4.34 |
| C29E4.2 | 168.83 | 343.99 | 2.04 | 31.44 | 68.32 | 2.17 |
| D1044.6 | 243.37 | 495.17 | 2.03 | 36.83 | 122.05 | 3.31 |
| C26E6.4 | 226.23 | 460.26 | 2.03 | 35.03 | 81.68 | 2.33 |
| F01G10.1.2 | 92.55 | 188.16 | 2.03 | 33.23 | 87.27 | 2.63 |
| W01B6.9 | 178.61 | 362.82 | 2.03 | 30.84 | 53.11 | 1.72 |

| | | | | | | |
|---|---|---|---|---|---|---|
| W07E6.4 | 130.59 | 265.04 | 2.03 | 30.54 | 40.68 | 1.33 |
| C06A5.1 | 183.45 | 371.77 | 2.03 | 26.65 | 38.82 | 1.46 |
| H25P06.2b | 136.30 | 276.14 | 2.03 | 37.43 | 76.71 | 2.05 |
| F43G6.1b | 283.06 | 571.88 | 2.02 | 44.91 | 110.25 | 2.45 |
| C16A3.3 | 194.48 | 392.23 | 2.02 | 39.22 | 57.14 | 1.46 |
| F45F2.10 | 282.67 | 569.90 | 2.02 | 43.71 | 79.50 | 1.82 |
| K08E3.4 | 305.61 | 616.08 | 2.02 | 38.62 | 162.11 | 4.20 |
| C43E11.10 | 502.71 | 1010.29 | 2.01 | 29.94 | 65.84 | 2.20 |
| K07H8.10 | 174.35 | 350.09 | 2.01 | 63.77 | 87.27 | 1.37 |
| Y71F9B.7 | 457.70 | 918.45 | 2.01 | 53.29 | 86.96 | 1.63 |
| F54D10.5 | 176.09 | 353.19 | 2.01 | 57.78 | 145.96 | 2.53 |
| C08B11.1 | 742.69 | 1488.70 | 2.00 | 73.65 | 187.27 | 2.54 |
| F31C3.9 | 869.41 | 1732.33 | 1.99 | 828.14 | 3253.42 | 3.93 |
| Y54E10A.15 | 125.56 | 250.08 | 1.99 | 28.14 | 37.58 | 1.34 |
| F28C6.2 | 359.34 | 715.67 | 1.99 | 45.81 | 142.24 | 3.11 |
| T23B12.4 | 116.94 | 232.62 | 1.99 | 28.74 | 56.83 | 1.98 |
| F32E10.4 | 268.73 | 534.39 | 1.99 | 34.13 | 55.90 | 1.64 |
| T21C9.13 | 71.83 | 142.67 | 1.99 | 27.54 | 43.48 | 1.58 |
| F28D9.1 | 70.67 | 140.35 | 1.99 | 26.95 | 52.80 | 1.96 |
| Y87G2A.5 | 189.25 | 375.21 | 1.98 | 83.83 | 159.32 | 1.90 |
| F23A7.4.2 | 136.30 | 269.77 | 1.98 | 51.20 | 90.68 | 1.77 |
| H38K22.1 | 139.79 | 276.65 | 1.98 | 29.34 | 64.91 | 2.21 |
| ZK507.6 | 416.94 | 824.97 | 1.98 | 29.94 | 90.99 | 3.04 |
| K01C8.5 | 142.69 | 282.33 | 1.98 | 43.71 | 67.70 | 1.55 |
| F43G6.1a | 284.32 | 561.30 | 1.97 | 41.62 | 117.39 | 2.82 |
| D1043.1 | 186.54 | 368.24 | 1.97 | 26.65 | 60.87 | 2.28 |
| W07G4.4 | 208.33 | 411.15 | 1.97 | 27.54 | 86.02 | 3.12 |
| K07C5.8 | 300.77 | 593.21 | 1.97 | 61.08 | 188.20 | 3.08 |
| F01G10.1.1 | 93.32 | 183.69 | 1.97 | 41.32 | 88.51 | 2.14 |
| T21C9.12 | 36.98 | 72.58 | 1.96 | 34.73 | 138.20 | 3.98 |
| R05D3.2 | 461.28 | 905.21 | 1.96 | 41.02 | 73.91 | 1.80 |
| T24D1.2 | 259.54 | 508.50 | 1.96 | 28.44 | 53.73 | 1.89 |
| F11H8.4a | 219.46 | 428.52 | 1.95 | 49.40 | 90.37 | 1.83 |
| Y75B8A.25 | 209.39 | 408.74 | 1.95 | 53.89 | 87.58 | 1.63 |
| Y105E8B.4 | 613.26 | 1195.02 | 1.95 | 73.35 | 193.48 | 2.64 |
| D1081.7b | 357.50 | 694.94 | 1.94 | 32.93 | 89.75 | 2.73 |
| C30C11.1 | 30.01 | 58.31 | 1.94 | 31.14 | 63.35 | 2.03 |
| R07B5.9b.1 | 167.76 | 325.93 | 1.94 | 26.05 | 71.43 | 2.74 |
| D2096.12 | 325.75 | 632.85 | 1.94 | 32.34 | 126.09 | 3.90 |
| F45E12.3 | 319.75 | 620.73 | 1.94 | 77.25 | 147.83 | 1.91 |
| F56A3.4 | 339.30 | 656.76 | 1.94 | 54.49 | 95.65 | 1.76 |
| D1014.8 | 471.35 | 912.00 | 1.93 | 40.12 | 114.91 | 2.86 |
| C26C6.1a | 120.81 | 233.74 | 1.93 | 26.35 | 43.79 | 1.66 |
| Y51H1A.4 | 158.28 | 306.15 | 1.93 | 36.53 | 58.07 | 1.59 |

| | | | | | | |
|---|---|---|---|---|---|---|
| F33E11.3 | 87.61 | 169.33 | 1.93 | 36.83 | 65.22 | 1.77 |
| B0280.5 | 443.76 | 857.48 | 1.93 | 79.34 | 287.89 | 3.63 |
| Y46G5A.31 | 211.33 | 407.71 | 1.93 | 32.04 | 91.93 | 2.87 |
| K07D4.3 | 92.84 | 178.70 | 1.92 | 39.82 | 65.22 | 1.64 |
| T05H10.1 | 222.94 | 428.35 | 1.92 | 73.35 | 97.52 | 1.33 |
| M03C11.7 | 124.10 | 238.30 | 1.92 | 29.34 | 73.60 | 2.51 |
| T21B10.3 | 235.91 | 452.69 | 1.92 | 26.95 | 55.90 | 2.07 |
| H02I12.1 | 707.45 | 1357.29 | 1.92 | 180.54 | 312.11 | 1.73 |
| C27F2.7 | 251.21 | 481.50 | 1.92 | 31.44 | 81.99 | 2.61 |
| B0523.5 | 298.64 | 572.22 | 1.92 | 28.74 | 72.98 | 2.54 |
| C08H9.2a | 107.45 | 205.88 | 1.92 | 33.53 | 78.88 | 2.35 |
| H27M09.3 | 113.94 | 217.74 | 1.91 | 39.82 | 73.91 | 1.86 |
| T05B9.1 | 84.51 | 161.50 | 1.91 | 32.04 | 54.66 | 1.71 |
| C38C3.5a.1 | 57.79 | 110.25 | 1.91 | 31.44 | 65.22 | 2.07 |
| K05C4.1 | 56.53 | 107.75 | 1.91 | 26.35 | 56.21 | 2.13 |
| F11H8.4b | 229.43 | 437.12 | 1.91 | 52.69 | 77.95 | 1.48 |
| F10G7.3 | 327.49 | 623.05 | 1.90 | 50.30 | 111.18 | 2.21 |
| Y18D10A.17 | 279.09 | 529.66 | 1.90 | 106.29 | 182.61 | 1.72 |
| ZC404.3a | 356.63 | 676.71 | 1.90 | 25.45 | 36.02 | 1.42 |
| Y48E1B.12 | 123.43 | 234.08 | 1.90 | 25.15 | 50.31 | 2.00 |
| F18E2.3 | 392.35 | 743.70 | 1.90 | 50.90 | 165.84 | 3.26 |
| H05L14.2 | 292.55 | 554.25 | 1.89 | 46.11 | 194.10 | 4.21 |
| B0365.3.2 | 76.48 | 144.56 | 1.89 | 28.44 | 65.22 | 2.29 |
| F58E10.4 | 370.86 | 700.62 | 1.89 | 33.83 | 109.01 | 3.22 |
| R119.7a | 152.95 | 288.95 | 1.89 | 26.95 | 77.64 | 2.88 |
| C06G3.2 | 1681.12 | 3173.12 | 1.89 | 304.49 | 517.70 | 1.70 |
| K08B12.5.2 | 159.24 | 300.56 | 1.89 | 36.23 | 61.49 | 1.70 |
| T18H9.7a | 421.78 | 793.50 | 1.88 | 34.43 | 118.01 | 3.43 |
| W02F12.3 | 96.42 | 179.91 | 1.87 | 27.25 | 92.86 | 3.41 |
| T09A5.10.1 | 309.10 | 576.35 | 1.86 | 32.34 | 64.60 | 2.00 |
| T12E12.1 | 329.53 | 614.11 | 1.86 | 78.74 | 189.75 | 2.41 |
| ZK328.5b | 319.36 | 595.01 | 1.86 | 35.93 | 126.71 | 3.53 |
| C38C3.5a.2 | 58.28 | 108.53 | 1.86 | 31.14 | 76.09 | 2.44 |
| R151.8 | 227.20 | 422.85 | 1.86 | 53.89 | 124.22 | 2.31 |
| F58G1.1 | 1803.39 | 3354.32 | 1.86 | 310.48 | 477.33 | 1.54 |
| K07A12.2 | 876.19 | 1627.58 | 1.86 | 129.64 | 278.57 | 2.15 |
| F12F6.6 | 349.47 | 649.02 | 1.86 | 46.11 | 110.56 | 2.40 |
| Y38A10A.5.2 | 75.41 | 140.00 | 1.86 | 44.01 | 79.19 | 1.80 |
| D1081.7a | 359.73 | 667.17 | 1.85 | 32.34 | 81.99 | 2.54 |
| F20D12.2 | 174.54 | 323.52 | 1.85 | 35.63 | 58.70 | 1.65 |
| T21E3.1 | 430.88 | 796.85 | 1.85 | 109.28 | 218.63 | 2.00 |
| ZC518.2 | 352.95 | 652.29 | 1.85 | 56.29 | 126.40 | 2.25 |
| F22B3.4 | 273.86 | 505.49 | 1.85 | 69.76 | 127.02 | 1.82 |

| | | | | | | |
|---|---|---|---|---|---|---|
| B0303.9 | 231.95 | 427.92 | 1.84 | 40.42 | 70.81 | 1.75 |
| C08B6.7b | 49.18 | 90.55 | 1.84 | 43.71 | 98.45 | 2.25 |
| Y43F8C.14 | 57.50 | 105.69 | 1.84 | 26.05 | 56.21 | 2.16 |
| ZC404.9 | 412.20 | 756.26 | 1.83 | 35.93 | 55.90 | 1.56 |
| T07C4.1.1 | 79.09 | 144.99 | 1.83 | 34.13 | 111.80 | 3.28 |
| Y71F9B.3 | 57.12 | 104.66 | 1.83 | 28.74 | 75.47 | 2.63 |
| F18C5.3 | 171.35 | 313.63 | 1.83 | 40.12 | 89.13 | 2.22 |
| F26B1.3 | 383.25 | 701.31 | 1.83 | 75.45 | 243.79 | 3.23 |
| Y55B1AR.2a | 297.58 | 543.42 | 1.83 | 29.94 | 73.91 | 2.47 |
| C38C10.4 | 309.39 | 563.37 | 1.82 | 84.43 | 118.94 | 1.41 |
| T22D1.5 | 504.45 | 918.02 | 1.82 | 86.23 | 136.34 | 1.58 |
| Y53F4B.15 | 37.37 | 67.94 | 1.82 | 33.53 | 45.96 | 1.37 |
| K03H1.2 | 279.38 | 507.81 | 1.82 | 54.19 | 100.00 | 1.85 |
| B0240.4 | 124.39 | 225.66 | 1.81 | 27.84 | 63.98 | 2.30 |
| ZK1055.1 | 1231.17 | 2230.76 | 1.81 | 277.54 | 505.90 | 1.82 |
| T27F2.1 | 84.03 | 152.13 | 1.81 | 26.95 | 37.27 | 1.38 |
| R12E2.10 | 433.40 | 784.12 | 1.81 | 117.07 | 241.30 | 2.06 |
| T07C4.1.2 | 80.15 | 144.91 | 1.81 | 28.74 | 95.34 | 3.32 |
| C47E8.5.1 | 217.42 | 391.89 | 1.80 | 48.20 | 124.22 | 2.58 |
| Y32H12A.5 | 566.60 | 1019.32 | 1.80 | 51.20 | 113.35 | 2.21 |
| F22B7.6b | 55.28 | 99.41 | 1.80 | 25.75 | 49.69 | 1.93 |
| H37A05.1 | 241.63 | 434.54 | 1.80 | 41.32 | 78.57 | 1.90 |
| C41G7.4 | 125.36 | 225.31 | 1.80 | 36.83 | 48.45 | 1.32 |
| Y39G10AR.13b.2 | 209.49 | 376.32 | 1.80 | 59.28 | 90.06 | 1.52 |
| Y17G9B.9 | 388.19 | 696.40 | 1.79 | 60.48 | 83.23 | 1.38 |
| F25D7.2 | 126.14 | 226.26 | 1.79 | 38.32 | 68.63 | 1.79 |
| C27F2.8 | 126.14 | 226.17 | 1.79 | 51.50 | 69.88 | 1.36 |
| T10B5.6 | 341.34 | 610.84 | 1.79 | 90.42 | 145.03 | 1.60 |
| C26D10.1 | 127.49 | 227.98 | 1.79 | 33.83 | 46.89 | 1.39 |
| K06A5.4 | 446.27 | 797.80 | 1.79 | 73.65 | 98.76 | 1.34 |
| C38D4.3 | 583.83 | 1043.14 | 1.79 | 147.60 | 427.33 | 2.90 |
| C25A1.10b | 74.25 | 132.52 | 1.78 | 25.45 | 46.89 | 1.84 |
| F46F11.2 | 244.34 | 435.92 | 1.78 | 68.26 | 130.43 | 1.91 |
| W09D10.2 | 698.94 | 1245.07 | 1.78 | 90.42 | 171.12 | 1.89 |
| W02A2.7 | 230.59 | 410.46 | 1.78 | 28.74 | 115.84 | 4.03 |
| C27A2.6 | 202.23 | 359.64 | 1.78 | 38.32 | 64.60 | 1.69 |
| Y79H2A.11 | 415.59 | 738.80 | 1.78 | 71.86 | 110.25 | 1.53 |
| C08B6.7a | 43.56 | 77.40 | 1.78 | 50.90 | 92.55 | 1.82 |
| M4.1b | 140.37 | 249.39 | 1.78 | 39.52 | 88.20 | 2.23 |
| C05D2.5 | 113.26 | 201.06 | 1.78 | 31.44 | 57.45 | 1.83 |
| C47E8.5.3 | 219.46 | 389.14 | 1.77 | 56.59 | 124.53 | 2.20 |
| F44B9.3b | 395.45 | 700.79 | 1.77 | 30.54 | 48.45 | 1.59 |
| C01H6.9 | 302.42 | 535.50 | 1.77 | 54.49 | 113.35 | 2.08 |

| | | | | | | |
|---|---|---|---|---|---|---|
| Y34D9A.10 | 85.48 | 151.01 | 1.77 | 27.54 | 50.00 | 1.82 |
| Y110A7A.18 | 559.54 | 987.07 | 1.76 | 97.01 | 158.07 | 1.63 |
| Y55F3AM.12 | 280.93 | 495.26 | 1.76 | 41.92 | 55.90 | 1.33 |
| T03F1.9 | 615.39 | 1083.91 | 1.76 | 104.19 | 236.34 | 2.27 |
| F23C8.4 | 138.72 | 243.80 | 1.76 | 39.52 | 81.99 | 2.07 |
| Y39G10AR.13a | 223.43 | 392.49 | 1.76 | 62.57 | 100.00 | 1.60 |
| T09A5.10.2 | 320.43 | 562.08 | 1.75 | 34.43 | 60.25 | 1.75 |
| C07H4.2 | 245.01 | 429.73 | 1.75 | 33.83 | 50.00 | 1.48 |
| F07F6.4 | 125.85 | 220.67 | 1.75 | 30.24 | 53.11 | 1.76 |
| F22B7.13 | 362.63 | 635.26 | 1.75 | 80.84 | 142.86 | 1.77 |
| T20D3.11b | 242.69 | 424.57 | 1.75 | 30.54 | 47.20 | 1.55 |
| F57B9.6a.2 | 115.39 | 201.75 | 1.75 | 31.14 | 49.69 | 1.60 |
| C38D4.4 | 340.85 | 595.36 | 1.75 | 38.62 | 69.57 | 1.80 |
| M4.1a | 127.98 | 223.33 | 1.75 | 32.04 | 69.25 | 2.16 |
| F25B4.6 | 160.50 | 280.09 | 1.75 | 31.74 | 74.53 | 2.35 |
| C23G10.8.1 | 74.83 | 130.54 | 1.74 | 32.04 | 46.27 | 1.44 |
| ZK637.7b | 346.85 | 603.79 | 1.74 | 37.43 | 108.39 | 2.90 |
| F35F11.1 | 77.93 | 135.36 | 1.74 | 29.94 | 48.45 | 1.62 |
| ZC404.8.1 | 135.14 | 234.69 | 1.74 | 45.21 | 100.31 | 2.22 |
| T23D8.4 | 146.37 | 253.86 | 1.73 | 37.43 | 67.39 | 1.80 |
| F22B7.6a | 58.47 | 101.39 | 1.73 | 25.45 | 48.45 | 1.90 |
| ZK637.7a | 341.14 | 590.46 | 1.73 | 32.34 | 109.63 | 3.39 |
| Y62E10A.14 | 446.08 | 771.74 | 1.73 | 55.69 | 112.73 | 2.02 |
| H02I12.5 | 626.62 | 1083.82 | 1.73 | 66.17 | 111.49 | 1.68 |
| F58B3.4 | 76.77 | 132.69 | 1.73 | 27.25 | 37.89 | 1.39 |
| Y38A10A.5.1 | 77.35 | 133.30 | 1.72 | 37.13 | 90.06 | 2.43 |
| B0365.3.1 | 87.22 | 149.89 | 1.72 | 37.13 | 65.22 | 1.76 |
| F58G11.6 | 153.24 | 263.32 | 1.72 | 28.14 | 62.42 | 2.22 |
| B0035.6 | 129.91 | 222.90 | 1.72 | 47.01 | 61.49 | 1.31 |
| C47E8.5.4 | 219.65 | 376.75 | 1.72 | 46.11 | 133.54 | 2.90 |
| T01C3.3 | 449.47 | 769.67 | 1.71 | 83.53 | 187.27 | 2.24 |
| C23G10.8.2 | 76.48 | 130.72 | 1.71 | 30.54 | 45.34 | 1.48 |
| K08E3.6.2 | 174.35 | 297.98 | 1.71 | 29.04 | 65.53 | 2.26 |
| C52E12.4 | 412.20 | 704.32 | 1.71 | 45.21 | 70.19 | 1.55 |
| F54F2.2a.2 | 199.90 | 341.41 | 1.71 | 31.74 | 53.42 | 1.68 |
| C25G4.5 | 337.17 | 575.32 | 1.71 | 52.99 | 104.66 | 1.97 |
| Y105E8A.9 | 165.34 | 281.12 | 1.70 | 29.04 | 65.22 | 2.25 |
| F57B9.2 | 418.49 | 711.45 | 1.70 | 71.56 | 192.86 | 2.70 |
| C06E1.10 | 166.80 | 283.19 | 1.70 | 32.63 | 63.04 | 1.93 |
| Y57A10A.31 | 345.79 | 585.64 | 1.69 | 46.41 | 114.60 | 2.47 |
| F52B5.3 | 308.33 | 521.92 | 1.69 | 41.32 | 57.14 | 1.38 |

| | | | | | | |
|---|---|---|---|---|---|---|
| K04F10.7 | 90.71 | 153.50 | 1.69 | 42.22 | 54.97 | 1.30 |
| F25H5.5a | 124.10 | 209.57 | 1.69 | 37.72 | 56.83 | 1.51 |
| R144.7b | 137.66 | 232.45 | 1.69 | 28.74 | 51.55 | 1.79 |
| Y110A7A.19 | 63.60 | 107.15 | 1.68 | 32.63 | 69.25 | 2.12 |
| T24A11.1b | 50.53 | 85.05 | 1.68 | 25.15 | 33.54 | 1.33 |
| R119.4 | 249.76 | 419.75 | 1.68 | 72.16 | 112.73 | 1.56 |
| Y39G10AR.13b.1 | 229.91 | 385.27 | 1.68 | 65.57 | 97.83 | 1.49 |
| F25B3.1 | 196.71 | 328.68 | 1.67 | 35.33 | 77.95 | 2.21 |
| D2013.6 | 131.17 | 218.95 | 1.67 | 37.72 | 69.88 | 1.85 |
| T09F3.2 | 369.60 | 616.34 | 1.67 | 40.42 | 80.43 | 1.99 |
| F38B7.5 | 870.67 | 1451.46 | 1.67 | 55.99 | 125.16 | 2.24 |
| C50F4.11 | 458.95 | 763.31 | 1.66 | 73.05 | 151.86 | 2.08 |
| R05D3.11 | 348.69 | 579.53 | 1.66 | 90.42 | 132.61 | 1.47 |
| C29E4.4 | 325.17 | 540.06 | 1.66 | 27.54 | 56.52 | 2.05 |
| C06A8.5 | 556.24 | 922.92 | 1.66 | 81.74 | 135.09 | 1.65 |
| T20G5.11 | 130.11 | 215.85 | 1.66 | 26.35 | 41.61 | 1.58 |
| K09H11.3 | 1414.52 | 2344.45 | 1.66 | 109.58 | 191.93 | 1.75 |
| T20D3.11a | 252.95 | 419.24 | 1.66 | 29.94 | 47.52 | 1.59 |
| F19F10.11a | 71.83 | 118.85 | 1.65 | 28.14 | 44.10 | 1.57 |
| C47E8.5.2 | 239.01 | 395.33 | 1.65 | 86.23 | 127.33 | 1.48 |
| R09E12.3 | 92.84 | 153.50 | 1.65 | 28.44 | 44.41 | 1.56 |
| Y105E8A.19 | 122.94 | 202.95 | 1.65 | 27.84 | 44.10 | 1.58 |
| F15D4.1 | 631.56 | 1041.42 | 1.65 | 62.57 | 121.74 | 1.95 |
| R144.10 | 132.14 | 217.83 | 1.65 | 32.04 | 50.62 | 1.58 |
| R01H10.8 | 232.04 | 380.45 | 1.64 | 47.31 | 85.09 | 1.80 |
| ZK154.5 | 169.70 | 277.94 | 1.64 | 41.02 | 81.68 | 1.99 |
| F56D2.6b | 140.95 | 230.73 | 1.64 | 26.35 | 40.37 | 1.53 |
| C50B6.3b | 104.16 | 170.27 | 1.63 | 29.64 | 41.93 | 1.41 |
| F54D11.2.2 | 199.81 | 326.62 | 1.63 | 27.84 | 38.20 | 1.37 |
| F57B9.6a.1 | 118.78 | 194.01 | 1.63 | 30.24 | 59.32 | 1.96 |
| F56D12.5a.2 | 180.25 | 294.37 | 1.63 | 34.73 | 49.07 | 1.41 |
| T01G1.3 | 382.19 | 622.88 | 1.63 | 40.12 | 71.12 | 1.77 |
| C10H11.10 | 1247.34 | 2032.63 | 1.63 | 133.83 | 349.07 | 2.61 |
| M04F3.5 | 506.68 | 825.31 | 1.63 | 64.07 | 149.69 | 2.34 |
| C25H3.8 | 236.88 | 385.70 | 1.63 | 38.62 | 74.22 | 1.92 |
| Y2H9A.1 | 227.59 | 369.70 | 1.62 | 34.13 | 50.31 | 1.47 |
| C01C7.1a | 176.48 | 286.63 | 1.62 | 47.60 | 62.42 | 1.31 |
| T20G5.1 | 270.76 | 439.53 | 1.62 | 38.92 | 85.09 | 2.19 |
| K12D12.2.1 | 163.41 | 265.04 | 1.62 | 51.50 | 136.96 | 2.66 |
| T06E4.1 | 218.39 | 354.22 | 1.62 | 47.90 | 75.78 | 1.58 |
| R06C7.8 | 506.49 | 821.36 | 1.62 | 76.65 | 110.87 | 1.45 |
| Y67H2A.1 | 179.67 | 291.27 | 1.62 | 39.82 | 67.08 | 1.68 |
| F21G4.2 | 512.58 | 830.90 | 1.62 | 62.87 | 151.86 | 2.42 |

| | | | | | | |
|---|---|---|---|---|---|---|
| Y6D1A.1 | 332.14 | 536.88 | 1.62 | 52.99 | 150.62 | 2.84 |
| W10C6.1 | 318.20 | 513.83 | 1.61 | 43.71 | 57.45 | 1.31 |
| ZC404.8.2 | 142.88 | 230.56 | 1.61 | 52.10 | 98.14 | 1.88 |
| F55H2.6 | 231.95 | 374.17 | 1.61 | 25.45 | 38.82 | 1.53 |
| C32E8.8 | 270.67 | 434.80 | 1.61 | 50.00 | 103.11 | 2.06 |
| F39B2.11 | 66.70 | 106.89 | 1.60 | 30.24 | 49.38 | 1.63 |
| Y32H12A.8 | 59.54 | 94.94 | 1.59 | 35.93 | 60.25 | 1.68 |
| Y18D10A.1 | 165.73 | 263.93 | 1.59 | 30.54 | 59.63 | 1.95 |
| F28B3.1.2 | 122.75 | 195.47 | 1.59 | 27.25 | 41.61 | 1.53 |
| K02F2.4 | 285.09 | 452.26 | 1.59 | 60.78 | 93.48 | 1.54 |
| F54C8.3 | 175.99 | 278.63 | 1.58 | 29.64 | 44.10 | 1.49 |
| C14B9.8 | 212.78 | 336.33 | 1.58 | 56.29 | 79.50 | 1.41 |
| Y47G6A.23 | 133.30 | 210.69 | 1.58 | 25.75 | 50.31 | 1.95 |
| Y45F10A.2 | 563.60 | 890.41 | 1.58 | 73.65 | 231.37 | 3.14 |
| F21H12.6 | 129.14 | 203.99 | 1.58 | 31.44 | 68.63 | 2.18 |
| Y48G1BL.2b | 129.82 | 204.16 | 1.57 | 41.62 | 72.98 | 1.75 |
| W04A8.7 | 128.65 | 202.01 | 1.57 | 32.04 | 50.62 | 1.58 |
| C50B6.3a | 111.13 | 172.34 | 1.55 | 26.35 | 54.66 | 2.07 |
| F53H1.3a | 156.44 | 242.00 | 1.55 | 29.94 | 50.31 | 1.68 |
| T07G12.11 | 64.28 | 99.07 | 1.54 | 30.54 | 45.65 | 1.49 |
| F20D12.1b | 455.37 | 701.65 | 1.54 | 39.52 | 60.56 | 1.53 |
| K08H10.7 | 146.95 | 226.26 | 1.54 | 25.45 | 60.25 | 2.37 |
| F32D1.10 | 264.18 | 406.16 | 1.54 | 32.34 | 58.39 | 1.81 |
| F31E3.4 | 323.33 | 495.17 | 1.53 | 42.22 | 75.16 | 1.78 |
| T22F3.3b.2 | 104.45 | 159.87 | 1.53 | 37.43 | 62.73 | 1.68 |
| F10G7.9a | 29.62 | 45.32 | 1.53 | 27.84 | 41.93 | 1.51 |
| F44B9.3a | 498.35 | 761.16 | 1.53 | 37.43 | 72.67 | 1.94 |
| ZK1067.2a | 264.09 | 402.21 | 1.52 | 41.62 | 64.91 | 1.56 |
| T22F3.3a | 104.94 | 159.70 | 1.52 | 42.22 | 66.15 | 1.57 |
| Y110A2AR.1 | 119.65 | 181.37 | 1.52 | 33.83 | 45.96 | 1.36 |
| C37H5.8 | 302.52 | 458.19 | 1.51 | 59.88 | 92.24 | 1.54 |
| C28A5.1 | 283.93 | 429.81 | 1.51 | 67.66 | 90.06 | 1.33 |
| F38E11.5 | 428.94 | 649.11 | 1.51 | 75.15 | 137.27 | 1.83 |
| F28D1.10 | 540.37 | 817.57 | 1.51 | 86.23 | 114.29 | 1.33 |
| F20D12.1a | 460.21 | 695.97 | 1.51 | 50.90 | 67.08 | 1.32 |
| T22F3.3b.1 | 107.07 | 161.85 | 1.51 | 37.13 | 72.05 | 1.94 |
| Y54G11A.3 | 127.49 | 192.72 | 1.51 | 46.71 | 64.91 | 1.39 |
| C14B9.4a | 200.10 | 302.28 | 1.51 | 49.40 | 64.60 | 1.31 |
| C28H8.3 | 283.74 | 428.44 | 1.51 | 37.43 | 72.36 | 1.93 |
| Y37E11B.4 | 248.89 | 374.60 | 1.51 | 36.53 | 83.85 | 2.30 |
| ZK973.2.2 | 121.10 | 182.23 | 1.50 | 32.93 | 48.76 | 1.48 |
| F25B5.4a.2 | 30.78 | 46.27 | 1.50 | 26.65 | 50.62 | 1.90 |
| C45G3.1 | 886.25 | 1328.57 | 1.50 | 104.79 | 257.14 | 2.45 |

| | | | | | | |
|---|---|---|---|---|---|---|
| T24C4.7 | 164.47 | 245.26 | 1.49 | 30.84 | 45.03 | 1.46 |
| T22F3.3b.3 | 105.32 | 156.26 | 1.48 | 40.72 | 70.19 | 1.72 |
| Y73E7A.2 | 108.33 | 160.64 | 1.48 | 25.45 | 39.13 | 1.54 |
| C14B9.4b | 202.81 | 300.65 | 1.48 | 48.50 | 70.19 | 1.45 |
| T23D8.9a | 629.04 | 927.48 | 1.47 | 29.94 | 65.84 | 2.20 |
| ZK973.2.1 | 123.62 | 182.23 | 1.47 | 35.03 | 56.21 | 1.60 |
| F25B5.4a.1 | 30.78 | 45.32 | 1.47 | 27.25 | 51.86 | 1.90 |
| Y43F4B.3 | 416.55 | 611.78 | 1.47 | 51.80 | 104.35 | 2.01 |
| C39E9.12 | 637.66 | 934.62 | 1.47 | 204.19 | 420.19 | 2.06 |
| ZK1251.9 | 306.49 | 448.99 | 1.46 | 57.49 | 77.64 | 1.35 |
| ZC308.1a | 102.23 | 149.46 | 1.46 | 26.65 | 46.89 | 1.76 |
| Y46G5A.5 | 693.13 | 1012.70 | 1.46 | 43.41 | 95.34 | 2.20 |
| Y47G6A.11 | 165.25 | 241.39 | 1.46 | 50.30 | 87.58 | 1.74 |
| Y48C3A.7 | 132.24 | 192.55 | 1.46 | 63.17 | 84.16 | 1.33 |
| VF13D12L.1 | 264.96 | 385.18 | 1.45 | 38.62 | 71.74 | 1.86 |
| F57C9.5 | 223.91 | 324.98 | 1.45 | 41.62 | 65.22 | 1.57 |
| Y54E5A.4 | 392.64 | 569.13 | 1.45 | 31.74 | 41.93 | 1.32 |
| F48E8.6 | 707.36 | 1022.51 | 1.45 | 76.35 | 139.75 | 1.83 |
| F18C5.2 | 322.36 | 465.16 | 1.44 | 40.42 | 75.16 | 1.86 |
| T08A11.1 | 167.67 | 241.82 | 1.44 | 45.81 | 73.29 | 1.60 |
| F13B12.6 | 136.30 | 195.82 | 1.44 | 31.74 | 53.11 | 1.67 |
| C07E3.2 | 100.97 | 144.65 | 1.43 | 27.25 | 39.75 | 1.46 |
| C50C3.6 | 309.39 | 442.28 | 1.43 | 64.67 | 96.89 | 1.50 |
| T12D8.1 | 690.32 | 983.72 | 1.43 | 166.47 | 349.69 | 2.10 |
| F26D10.3.2 | 466.02 | 661.83 | 1.42 | 110.18 | 285.09 | 2.59 |
| Y75B12B.5 | 46.47 | 65.96 | 1.42 | 44.61 | 90.06 | 2.02 |
| Y66D12A.15 | 229.82 | 325.76 | 1.42 | 60.48 | 131.99 | 2.18 |
| Y43F4B.6 | 1401.45 | 1982.92 | 1.41 | 245.21 | 427.33 | 1.74 |
| F54E7.8 | 130.30 | 184.03 | 1.41 | 26.05 | 40.99 | 1.57 |
| F26D10.3.1 | 465.54 | 657.45 | 1.41 | 116.47 | 304.97 | 2.62 |
| F55A11.3 | 96.90 | 136.82 | 1.41 | 29.64 | 45.65 | 1.54 |
| Y25C1A.5 | 161.67 | 228.15 | 1.41 | 38.32 | 55.28 | 1.44 |
| F22B5.7 | 665.15 | 937.54 | 1.41 | 170.36 | 498.14 | 2.92 |
| W09B6.3 | 110.75 | 155.74 | 1.41 | 48.80 | 73.60 | 1.51 |
| F32D1.1 | 366.51 | 515.21 | 1.41 | 62.87 | 97.52 | 1.55 |
| C47D12.8 | 258.66 | 363.60 | 1.41 | 37.43 | 87.27 | 2.33 |
| Y39G10AR.7b | 242.88 | 340.72 | 1.40 | 32.63 | 55.59 | 1.70 |
| W02D3.2 | 72.70 | 101.91 | 1.40 | 43.41 | 92.24 | 2.12 |
| Y37E11AL.8 | 322.94 | 452.60 | 1.40 | 29.04 | 44.72 | 1.54 |
| F53G2.6 | 257.12 | 358.78 | 1.40 | 75.75 | 141.93 | 1.87 |
| Y71F9B.10a | 89.25 | 124.52 | 1.40 | 44.91 | 86.02 | 1.92 |
| R07H5.8 | 101.16 | 139.83 | 1.38 | 31.74 | 95.03 | 2.99 |
| ZK1067.3 | 311.04 | 428.87 | 1.38 | 43.11 | 59.63 | 1.38 |

| | | | | | | |
|---|---|---|---|---|---|---|
| C30G12.7 | 511.42 | 702.25 | 1.37 | 93.41 | 144.72 | 1.55 |
| C03D6.4 | 259.15 | 355.68 | 1.37 | 35.63 | 101.86 | 2.86 |
| Y39G10AR.7a | 243.66 | 334.18 | 1.37 | 29.64 | 54.66 | 1.84 |
| T06E4.3b | 235.04 | 321.11 | 1.37 | 26.65 | 38.51 | 1.45 |
| T01C3.1 | 770.09 | 1048.30 | 1.36 | 33.23 | 78.88 | 2.37 |
| Y34D9A.4 | 738.33 | 1004.45 | 1.36 | 134.73 | 204.66 | 1.52 |
| ZK430.1 | 349.27 | 473.41 | 1.36 | 104.19 | 153.73 | 1.48 |
| Y18D10A.13 | 377.44 | 510.22 | 1.35 | 35.93 | 62.42 | 1.74 |
| F28H1.3.2 | 107.26 | 144.39 | 1.35 | 25.75 | 39.44 | 1.53 |
| C44B9.4 | 189.64 | 255.07 | 1.34 | 25.15 | 43.48 | 1.73 |
| T22B11.5b.2 | 71.83 | 96.49 | 1.34 | 27.54 | 37.89 | 1.38 |
| T24H10.1 | 136.30 | 183.00 | 1.34 | 26.65 | 42.24 | 1.59 |
| F36D4.3b | 478.90 | 641.71 | 1.34 | 108.38 | 151.24 | 1.40 |
| Y81G3A.3b | 271.35 | 362.39 | 1.34 | 31.44 | 71.12 | 2.26 |
| F36D4.3a | 476.86 | 635.60 | 1.33 | 89.52 | 161.80 | 1.81 |
| C18G1.4b | 104.94 | 139.49 | 1.33 | 35.33 | 47.83 | 1.35 |
| C29H12.5 | 290.71 | 385.61 | 1.33 | 47.90 | 83.85 | 1.75 |
| F14B4.3 | 227.49 | 301.08 | 1.32 | 40.12 | 52.80 | 1.32 |
| F23H11.1 | 230.01 | 304.09 | 1.32 | 61.98 | 291.93 | 4.71 |
| Y73B6BL.38 | 569.80 | 752.04 | 1.32 | 96.41 | 231.68 | 2.40 |
| Y81G3A.3a | 266.21 | 349.32 | 1.31 | 27.25 | 67.08 | 2.46 |
| T23G7.4 | 118.97 | 156.00 | 1.31 | 50.60 | 73.91 | 1.46 |
| T06E4.3a | 238.53 | 311.57 | 1.31 | 27.54 | 36.65 | 1.33 |
| F20H11.3 | 159.83 | 208.03 | 1.30 | 26.95 | 54.66 | 2.03 |

**Table S2. List of strains used in this study.**

| Strains | Genotype |
|---|---|
| N2 | |
| YY174 | *3xflag::gfp::nrde-3(ggIS1)* |
| YY158 | *nrde-3(gg66)* |
| SHG487 | *3xflag::gfp::wago-4(ustIS052)* |
| SHG492 | *drh-3(ne4253);3xflag::gfp::wago-4(ustIS052)* |
| SHG498 | *3xflag::gfp::hrde-1(ustIS068)* |
| SHG674 | *drh-3(ne4253);3xflag::gfp::hrde-1(ustIS068)* |
| SHG750 | *Pmex-5::3xflag::gfp::nrde-3::tbb-2(3'UTR)(ustIS094)* |
| SHG852 | *3xflag::gfp::nrde-2(ustIS117)* |
| SHG1059 | *drh-3(ne4253);Pmex-5::3xflag::gfp::nrde-3::tbb-2(3'UTR)(ustIS094)* |
| SHG1060 | *Pmex-5::3xflag::gfp::nrde-3(*PAZ)::tbb-2(3'UTR)(ustIS146)* |
| SHG1063 | *drh-3(ne4253);Pmex-5::3xflag::gfp::nrde-3(*PAZ)::tbb-2(3'UTR)(ustIS146)* |
| SHG1064 | *Pmex-5::mCherry::tbg-1::tbb-2(3'UTR)(ustIS147)* |
| SHG1070 | *ergo-1(gg98);Pmex-5::mCherry::air-1::tbb-2(3'UTR)(ustIS148);Pmex-5::3xflag::gfp::nrde-3::tbb-2(3'UTR)(ustIS094)* |
| SHG1074 | *eri-1(mg366);Pmex-5::mCherry::tbg-1::tbb-2(3'UTR)(ustIS147);Pmex-5::3xflag::gfp::nrde-3::tbb-2(3'UTR)(ustIS094)* |
| SHG1075 | *ergo-1(gg98);Pmex-5::mCherry::tbg-1::tbb-2(3'UTR)(ustIS147);Pmex-5::3xflag::gfp::nrde-3::tbb-2(3'UTR)(ustIS094)* |
| SHG1076 | *drh-3(ne4253);Pmex-5::mCherry::tbg-1::tbb-2(3'UTR)(ustIS147);Pmex-5::3xflag::gfp::nrde-3::tbb-2(3'UTR)(ustIS094)* |
| SHG1080 | *Pmex-5::mCherry::air-1::tbb-2(3'UTR)(ustIS148)* |
| SHG1083 | *eri-1(mg366);Pmex-5::mCherry::air-1::tbb-2(3'UTR)(ustIS148);Pmex-5::3xflag::gfp::nrde-3::tbb-2(3'UTR)(ustIS094)* |
| SHG1084 | *Pmex-5::mCherry::sas-4::tbb-2(3'UTR)(ustIS149)* |
| SHG1107 | *drh-3(ne4253);Pmex-5::mCherry::air-1::tbb-2(3'UTR)(ustIS148);Pmex-5::3xflag::gfp::nrde-3::tbb-2(3'UTR)(ustIS094)* |
| SHG1204 | *eri-1(mg366);Pmex-5::mCherry::sas-4::tbb-2(3'UTR)(ustIS149);Pmex-5::3xflag::gfp::nrde-3::tbb-2(3'UTR)(ustIS094)* |
| SHG1205 | *ergo-1(gg98);Pmex-5::mCherry::sas-4::tbb-2(3'UTR)(ustIS149);Pmex-5::3xflag::gfp::nrde-3::tbb-2(3'UTR)(ustIS094)* |

| | |
|---|---|
| SHG1206 | *drh-3(ne4253);Pmex-5::mCherry::sas-4::tbb-2(3'UTR)(ustIS149);Pmex-5::3xflag::gfp::nrde-3::tbb-2(3'UTR)(ustIS094)* |
| SHG1271 | *eri-1(mg366);Pmex-5::mCherry::tbg-1::tbb-2(3'UTR)(ustIS147);gfp::csr-1(ustIS150)* |
| SHG1273 | *Pmex-5::mCherry::tbg-1::tbb-2(3'UTR)(ustIS147);gfp::csr-1(ustIS150)* |
| SHG1274 | *eri-1(mg366);Pmex-5::mCherry::air-1::tbb-2(3'UTR)(ustIS148);gfp::csr-1(ustIS150)* |
| SHG1275 | *Pmex-5::mCherry::air-1::tbb-2(3'UTR)(ustIS148);gfp::csr-1(ustIS150)* |
| SHG1278 | *Pmex-5::mCherry::sas-5a::tbb-2(3'UTR)(ustIS152)* |
| SHG1279 | *drh-3(ne4253);Pmex-5::mCherry::sas-5a::tbb-2(3'UTR)(ustIS152);Pmex-5::3xflag::gfp::nrde-3::tbb-2(3'UTR)(ustIS094)* |
| SHG1282 | *Pmex-5::mCherry::sas-6::tbb-2(3'UTR)(ustIS153)* |
| SHG1283 | *drh-3(ne4253);Pmex-5::mCherry::sas-6::tbb-2(3'UTR)(ustIS153);Pmex-5::3xflag::gfp::nrde-3::tbb-2(3'UTR)(ustIS094)* |
| SHG1286 | *Pmex-5::3xflag::gfp::nrde-3(*NLS)::tbb-2(3'UTR)(ustIS154)* |
| SHG1287 | *drh-3(ne4253);Pmex-5::3xflag::gfp::nrde-3(*NLS)::tbb-2(3'UTR)(ustIS154)* |
| SHG1324 | *Pmex-5::3xflag::gfp::nrde-3(*NLS)::tbb-2(3'UTR)(ustIS154);Pmex-5::3xHA::tagRFP::tbb-2::tbb-2(3'UTR)(ustIS236)* |
| SHG1416 | *drh-3(ne4253);3xflag::gfp::nrde-2(ustIS117)* |
| SHG1426 | *3xflag::gfp::nrde-3(ust574)* |
| SHG1462 | *eri-1(mg366);3xflag::gfp::nrde-3(ust574)* |
| SHG1463 | *ergo-1(gg98);3xflag::gfp::nrde-3(ust574)* |
| SHG1464 | *drh-3(ne4253);3xflag::gfp::nrde-3(ust574)* |
| SHG1465 | *eri-1(mg366);rrf-1(pk1417);rrf-2(ok210);3xflag::gfp::nrde-3(ust574)* |
| SHG1478 | *rrf-1(pk1417);rrf-2(ok210);3xflag::gfp::nrde-3(ust574)* |
| SHG1494 | *Pmex-5::tagRFP::tbb-2::tbb-2(3'UTR)(ustIS191)* |
| SHG1549 | *drh-3(ne4253);3xflag::gfp::nrde-3(ust574);Pmex-5::tagRFP::tbb-2::tbb-2(3'UTR)(ustIS191)* |
| SHG1645 | *degron::ego-1(ust614);Psun-1::TIR1::mRuby::sun-1(3'UTR)* |
| SHG1798 | *Pmex-5::3xHA::tagRFP::tbb-2::tbb-2(3'UTR)(ustIS236)* |
| SHG1805 | *drh-3(ne4253);Pmex-5::3xflag::gfp::nrde-3(*NLS)::tbb-2(3'UTR)(ustIS154);Pmex-5::3xHA::tagRFP::tbb-2::tbb-2(3'UTR)(ustIS236)* |
| SHG2124 | *Pmex-5::bfp::his-58::tbb-2::tbb-2(3'UTR)(ustIS323);Pmex-5::3xflag::gfp::nrde-3(*PAZ)::tbb-2(3'UTR)(ustIS146);Pmex-5::3xHA::tagRFP::tbb-2::tbb-2(3'UTR)(ustIS236)* |
| SHG2125 | *drh-3(ne4253);Pmex-5::bfp::his-58::tbb-2::tbb-2(3'UTR)(ustIS323);Pmex-5::3xflag::gfp::nrde-3(*PAZ)::tbb-* |

| | |
|---|---|
| | *2(3'UTR)(ustIS146);Pmex-5::3xHA::tagRFP::tbb-2::tbb-2(3'UTR)(ustIS236)* |
| SHG2171 | *Pmex-5::bfp::his-58::tbb-2::tbb-2(3'UTR)(ustIS323);Pmex-5::3xflag::gfp::nrde-3::tbb-2(3'UTR)(ustIS094);Pmex-5::3xHA::tagRFP::tbb-2::tbb-2(3'UTR)(ustIS236)* |
| SHG2172 | *drh-3(ne4253);Pmex-5::bfp::his-58::tbb-2::tbb-2(3'UTR)(ustIS323);Pmex-5::3xflag::gfp::nrde-3::tbb-2(3'UTR)(ustIS094);Pmex-5::3xHA::tagRFP::tbb-2::tbb-2(3'UTR)(ustIS236)* |
| SHG2173 | *Pmex-5::bfp::his-58::tbb-2::tbb-2(3'UTR)(ustIS323);drh-3(ne4253);Pmex-5::3xflag::gfp::nrde-3(*NLS)::tbb-2(3'UTR)(ustIS154);Pmex-5::3xHA::tagRFP::tbb-2::tbb-2(3'UTR)(ustIS236)* |
| SHG2174 | *drh-3(ne4253);Pmex-5::bfp::his-58::tbb-2::tbb-2(3'UTR)(ustIS323);drh-3(ne4253);Pmex-5::3xflag::gfp::nrde-3(*NLS)::tbb-2(3'UTR)(ustIS154);Pmex-5::3xHA::tagRFP::tbb-2::tbb-2(3'UTR)(ustIS236)* |
| SHG2175 | *Pmex-5::bfp::his-58::tbb-2::tbb-2(3'UTR)(ustIS323)* |
| SHG2990 | *degron::ego-1(ust614);Psun-1::TIR1::mRuby::sun-1(3'UTR);3xflag::gfp::nrde-3(ust574)* |
| SHG3074 | *rrf-1(pk1417);rrf-2(ok210);degron::ego-1(ust614);Psun-1::TIR1::mRuby::sun-1(3'UTR);3xflag::gfp::nrde-3(ust574)* |
| SHG3075 | *eri-1(mg366);rrf-1(pk1417);rrf-2(ok210);degron::ego-1(ust614);sun-1p::TIR1::mRuby::sun-1 3'UTR;3xflag::gfp::nrde-3(ust574)* |

**Table S3. Sequence of sgRNAs for CRISPR/Cas9-mediated gene editing.**

| sgRNA name | Sequence (5' to 3') |
|---|---|
| *gfp-nrde-3*-sgRNA#1 | CAACTAATCATGGATCTCATGG |
| *gfp-nrde-3*-sgRNA#2 | ACAAAGTAATGGGTGAGATGGG |
| *gfp-nrde-3*-sgRNA#3 | TGGATGACGCAGATGTGGCTGG |
| chrII-sgRNA#1 | AAGTGAGTTTGCTACCATCATGG |
| chrII-sgRNA#2 | AGAAAACTAACGTGTTCTCCTGG |
| chrII-sgRNA#3 | GATATCAGTCTGTTTCGTAACGG |
| chrI-sgRNA#1 | GAAATCGCCGACTTGCGAGGAGG |
| chrI-sgRNA#2 | GCAATGACTAACCGATTTTCGGG |
| chrI-sgRNA#3 | TTCGGGATAATTGAGATGAGGGG |
| *bfp-his-58*-sgRNA#1 | CTGTTGGATGCCTGTGTAGCGG |
| *bfp-his-58*-sgRNA#2 | AGAAATAGTACAGCAAACGCGG |
| *bfp-his-58*-sgRNA#3 | GCCATAAAGGAAAACGGTACGG |
| *bfp-his-58*-sgRNA#4 | CCTCCACGCTATAATTTTTTGC |
| *degron-ego-1*-sgRNA#1 | TCCCGAATCCTCGCAACAA |
| *degron-ego-1*-sgRNA#2 | GGACGAAGGTTATCGTGGA |
| *degron-ego-1*-sgRNA#3 | TCCCTGTTCTCTGCCAGAG |

**Table S4. Sequences of the quantitative real-time PCR primers.**

| Primer name | Sequence (5' to 3') |
|---|---|
| *eft-3*-F | GTAAGGGATCTTTCAAGTACGC |
| *eft-3*-R | CATCGATGATGGTGATGTAGTAC |
| *ama-1*-F | CGAACCTGCCGATTGATA |
| *ama-1*-R | ACCACGATTGACCAACTC |
| GFP qRT 2F | AGGTGCTGAAGTCAAGTTTG |
| GFP qRT 2R | GCCATGATGTATACATTGTGTGAG |
